# Red Algae-Derived Mineral Intervention to Counter Pro-inflammatory Activity in Human Colon Organoids

**DOI:** 10.1101/2022.12.15.520649

**Authors:** James Varani, Shannon D McClintock, Daniyal M Nadeem, Isabelle Harber, Dania Zeidan, Muhammad N Aslam

## Abstract

Human colon organoids were maintained in culture medium alone or exposed to lipopolysaccharide with a combination of three pro-inflammatory cytokines (tumor necrosis factor-α, interleukin-1β and interferon-γ [LPS-cytokines]) to mimic the environment in the inflamed colon. Untreated organoids and those exposed to LPS-cytokines were concomitantly treated with a multimineral product that has previously been shown to improve barrier structure/function. The organoids were subjected to proteomic analysis to obtain a broad view of the protein changes induced by these interventions. In parallel, confocal fluorescence microscopy and trans-epithelial electrical resistance measurements were used to assess barrier structure/function. The LPS-cytokines altered expression of multiple proteins that influence innate immunity and promote inflammation. Most of these were unaffected by the multi-mineral intervention, though a subset of inflammation-related proteins including fibrinogen-β and -γ chains, phospholipase A2 and SPARC was down-regulated in the presence of the multi-mineral intervention; another subset of proteins with anti-inflammatory, antioxidant or antimicrobial activity was up-regulated by multi-mineral treatment. When used alone, the multi-mineral intervention strongly up-regulated proteins that contribute to barrier formation and tissue strength. Concomitant treatment with LPS-cytokines did not inhibit barrier formation in response to the multimineral intervention. Altogether, the findings suggest that mineral intervention may provide a novel approach to combating inflammation in the colon by improving barrier structure/function as well as by directly altering expression of pro-inflammatory proteins.

## INTRODUCTION

Ulcerative colitis (UC) is an inflammatory disease, and the therapeutic armamentarium currently used for UC is aimed, primarily, at controlling inflammation (1). Agents that target inflammation broadly including steroids and non-steroidal anti-inflammatories such as mesalamine have long been treatment options. More recently, biological agents targeting specific components of the process have become part of the treatment mix. Among the biological agents currently in use are tumor necrosis factor-α (TNF-α) inhibitors, interleukin-blocking molecules, agents that interfere with leukocyte trafficking as well as agents that interfere with up-stream signaling pathways that lead to cytokine generation and immune function (2). The use of biologicals has dramatically expanded in recent years, but clinical remission rates continue to be approximately 20% over placebo in clinical trials (3), suggesting that additional aspects of UC patho-physiology need to be addressed.

Abnormal barrier function in the gastrointestinal tract with increased mucosal permeability is a well-described pathologic feature of UC (4–8), but there are no therapies that directly address this fundamental disease element (9). The colonic barrier is a complex, multi-faceted entity. Tight junctional complexes are present at the apical surface of colonic epithelial cells. Tight junctions can open and close to regulate inter-cellular transport of soluble moieties (10–13). Adherens junctions are closely associated with tight junctional complexes and help organize these structures into functional units (14). Adherens junctions and, especially, desmosomes are also largely responsible for tissue cohesion and strength (15–17). Effective control of permeability in a mechanically active tissue such as the gastrointestinal tract cannot be maintained if tissue strength and cohesion are compromised. An additional contributor to barrier function is the mucinous layer covering the apical surface. This layer consists of mucin proteins organized with trefoils; the mucinous layer contributes to barrier function by trapping bacteria and other particulates in the colonic stream and preventing their reaching the epithelial surface (18,19).

Past work in our laboratory has shown that a red algae-derived multi-mineral product (Aquamin^®^) increases the elaboration of numerous proteins that contribute to barrier structure. These studies have been conducted using human colon tissue in organoid culture (20–22) and validated in a small interventional trial with healthy adult subjects (23). In the organoid culture studies (using colon tissue from healthy individuals or lesional tissue from individuals with UC), we demonstrated modest changes in tight junctional protein expression with the multi-mineral intervention but much stronger upregulation of several cadherins and desmosomal proteins. Also, strongly up-regulated were mucin proteins, trefoils and members of the carcinoembryonic antigen cell adhesion molecule (CEACAM) family. Improved permeability control and increased organoid cohesion accompanied mineral-induced changes in barrier protein expression. These findings highlight the importance of inorganic minerals to barrier formation in the gastrointestinal tract. They do not, however, indicate whether an effective barrier can be maintained in the face of an inflammatory attack.

As a way to begin addressing this issue, human colon organoids were exposed to lipopolysaccharide (LPS) in combination with three pro-inflammatory cytokines (LPS-cytokines) to mimic the environment in the inflamed colon (24). Untreated organoids and those exposed to the pro-inflammatory stimulus were concomitantly treated with the same multi-mineral natural product as used previously or kept as control. The colon organoids were then subjected to proteomic analysis to obtain a broad view of the protein changes induced by the two interventions (i.e., the pro-inflammatory mix and the multi-mineral product) alone and in combination. In parallel, confocal fluorescence microscopy and trans-epithelial electrical resistance (TEER) measurements were used to assess barrier structure/function. The results are presented here.

## MATERIALS AND METHODS

### LPS-cytokine mix

A combination of lipopolysaccharide (LPS) from *E. coli (Escherichia coli*) and three pro-inflammatory cytokines was used to mimic the environment in the chronically-inflamed colon (24). The stock solution for the mix consisted of LPS (1 μg/mL, Sigma), tumor necrosis factor-α (TNF-α; 50 ng/mL, Sigma), interleukin 1-β (IL-1β; 25 ng/mL, Shenandoah Biotech), and interferon-γ (IFN-γ; 50 ng/mL, Sigma). The pro-inflammatory mix was initially assessed over a wide range of concentrations for organoid toxicity at day-14 with cultures established from two subjects. Toxicity was defined based on morphological evidence of organoid growth suppression, organoid failure to demonstrate features of differentiation (formation of thick walls and decreased budding structures) and loss of tissue integrity. High concentrations (≤1:50 dilution) of the pro-inflammatory mix were toxic, based on these criteria. Findings were similar with organoids from either source, and based on the preliminary results, subsequent experiments were carried out using a 1:250 dilution of the LPS-cytokines stock solution.

### Aquamin^®^

Aquamin^®^ is a calcium-, magnesium- and multi-trace element-rich product harvested from the skeletal remains of the red marine algae, *Lithothamnion sp* (25) (Marigot Ltd, Cork, Ireland). The calcium and magnesium ratio in Aquamin^®^ is approximately (12:1); Aquamin^®^ also contains measurable levels of seventy-two trace minerals. Mineral composition of Aquamin^®^ was established via an independent laboratory (Advanced Laboratories; Salt Lake City, Utah) using Inductively Coupled Plasma Optical Emission Spectrometry. Supplement Table 1 is a list of elements detected in Aquamin^®^ and their relative amounts. Aquamin^®^ is sold as a dietary supplement (GRAS 000028) and is used in various products for human consumption in Europe, Asia, Australia, and North America. In the studies reported on here, Aquamin^®^ was used at an amount providing a final calcium concentration of 3 mM.

### Colon organoid culture

Normal appearing tissue obtained endoscopically from the sigmoid colon from four subjects was utilized. Demographic characteristics (age, gender, ethnicity and site) of the subjects providing tissue are present in Supplement Table 2. The collection and use of human colonic tissue was approved by the Institutional Review Board (IRBMED) at the University of Michigan and all subjects provided written informed consent prior to biopsy. This study was conducted according to the principles stated in the Declaration of Helsinki.

For the present work, cryopreserved samples from previous studies (20–22) were cultured in Matrigel (Corning) and expanded over a 3-4 week period. During the expansion phase, culture medium consisted of a 1:1 mix of Advanced DMEM/F12 (Invitrogen) and the same medium previously conditioned by the growth of L cells genetically modified to produce recombinant forms of Wnt3a, R-spondin-3 and Noggin (i.e., L-WRN) (26). The growth medium also contained 10% fetal bovine serum (Gibco) and the final calcium concentration was 1.0 mM. The medium was supplemented with 1X N2 (Invitrogen), 1X B-27 without vitamin A (Invitrogen), 1 mM N-Acetyl-L-cysteine (Sigma), 10 mM HEPES (Invitrogen), 2 mM Glutamax (Invitrogen), 100 μg/mL Primocin (InvivoGen) and small molecule inhibitors: 10 μM Y27632 (Tocris); as a ROCK inhibitor, 500 nM A83-01 (Tocris); a TGF-β inhibitor, 10 μM SB202190 (Sigma); a p38 inhibitor, along with 100 ng/mL EGF (R&D). For the first 10 days of expansion of culture, and for two days at each passage the medium was also supplemented with 2.5 μM CHIR99021 (Tocris).

During the experimental phase, established organoids were interrogated in L-WRN plus 10 μM Y27632 (but without the additional small molecules) diluted 1:4 with KGM Gold. KGM-Gold is a serum-free, calcium-free culture medium optimized for epithelial cell growth (Lonza). The final serum concentration in the L-WRN – KGM Gold culture medium was 2.5% and the calcium concentration was 0.25 mM. This control treatment medium was compared to the same medium supplemented with the pro-inflammatory (LPS-cytokine) mix at a 1:250 dilution of the stock material. Control organoid cultures and those exposed to the pro-inflammatory mix were maintained without additional treatment or treated with Aquamin^®^ in an amount to provide a final calcium concentration of 3.0 mM. Organoids were evaluated by phase-contrast microscopy (Hoffman Modulation Contrast - Olympus IX70 with a DP71 digital camera) for change in size and shape during the 14-day in-life portion of the study and at harvest. At harvest, organoids were prepared for either proteomic assessment or for confocal microscopy as described below.

### Organoid culture on transwell filters

For some experiments, colon organoids were dissociated into small cell aggregates (less than 40μm in size) and plated onto collagen IV (Sigma)-coated transwells (0.4 μm pore size, 0.33cm^2^, PET, Corning Costar) at 200,000 cell aggregates as described previously (21). Cells were seeded for attachment and initial growth in growth medium (as above). The medium was also supplemented with 10 nM Gastrin (Sigma), 50 ng/mL Noggin (R&D), 50 ng/mL EGF, and 2.5 μM Y27632. After 2 days, Y27632 was removed. After 24 hours, growth medium was replaced with L-WRN – KGM Gold along with the LPS-cytokine mix alone, Aquamin^®^ alone or the combination of the two. Fresh culture medium was added every two days during the incubation period. TEER values were determined daily with an Epithelial Volt ohm meter 2 (EVOM2) epithelial volt/ohm meter (World Precision Instruments) and STX2 series chopstick electrodes.

### Confocal fluorescence microscopy

Once the TEER assay was completed, membranes were prepared for confocal fluorescence microscopy. The membranes were fixed for 15 minutes at −20°C in 100% methanol. After fixation, membranes were washed three times in PBS before blocking in 3% BSA (Sigma) in PBS for 1 hour. Following this, membranes were stained with antibodies to occludin (331594; Invitrogen; 1:400), desmoglein-2 (53-9159-80; eBioscience; 1:200) and cadherin-17 (NBP2-12065AF488; Novus Biologicals; 1:200) for 1 hour in 1% BSA in PBS. Stained membranes were rinsed three times (5 minutes each) in PBS, stained with DAPI for 5 minutes to detect nuclei and washed with PBS three more times. Lastly, the membranes were gently cut from the transwell insert and mounted apical side up on Superfrost Plus glass slides (Fisher Scientific, Pittsburgh, PA) with Prolong Gold (P36930; Life Technologies Molecular Probes). The stained specimens were visualized and imaged with a Leica Stellaris, an inverted confocal microscope system (at University of Michigan Medical School Biomedical Research Core Facility).

### Differential proteomic analysis

Colon organoids were isolated from Matrigel using 2mM EDTA for 15 minutes and then exposed to Radioimmunoprecipitation assay (RIPA) - lysis and extraction buffer (Pierce, # 89901; ThermoFisher Scientific) for protein isolation, as described in our previous reports (20–22). Proteomic assessments were conducted by using mass spectrometry-based Tandem Mass Tagging system (TMT, ThermoFisher Scientific) in the Proteomics Resource Facility (PRF) housed in the Department of Pathology at the University of Michigan. For this, four subjects were assessed separately using TMT six-plex kits for each subject. First experiment was conducted to assess proteomic signature with LPS-cytokines alone, without any other intervention. Subsequent three experiments were conducted with complete set of samples. Fifty micrograms of organoid protein from each condition was digested separately with trypsin and individual samples labeled with one of six isobaric mass tags according to the manufacturer’s protocol. After labeling, equal amounts of peptide from each condition were mixed together. In order to achieve in-depth characterization of the proteome, the labeled peptides were fractionated using 2D-LC (basic pH reverse-phase separation followed by acidic pH reverse phase) and analyzed on a high-resolution, tribrid mass spectrometer (Orbitrap Fusion Tribrid, ThermoFisher Scientific) using conditions optimized at the PRF. MultiNotch MS3 analysis was employed to obtain accurate quantitation of the identified proteins/peptides (27). Data analysis was performed using Proteome Discoverer (v2.4, ThermoFisher Scientific). MS2 spectra were searched against UniProt human protein database (20353 sequences; downloaded on 06/20/2019) using the following search parameters: MS1 and MS2 tolerance were set to 10 ppm and 0.6 Da, respectively; carbamidomethylation of cysteines (57.02146 Da) and TMT labeling of lysine and N-termini of peptides (229.16293 Da) were considered static modifications; oxidation of methionine (15.9949 Da) and deamidation of asparagine and glutamine (0.98401 Da) were considered variable. Identified proteins and peptides were filtered to retain only those that passed ≤2% false discovery rate (FDR) threshold of detection. Quantitation was performed using high-quality MS3 spectra (Average signal-to-noise ratio of 6 and <40% isolation interference). Differential protein expression between conditions, normalizing to control (0.25mM calcium) for each subject’s specimens separately was established. Next, resulting data from the three datasets were combined. Proteins names were retrieved using Uniprot.org, and Reactome v82 (reactome.org) was used for pathway enrichment analyses (28). Only Proteins with a ≤2% FDR confidence of detection were included in the analyses. The initial analysis was based on an unbiased proteome-wide screen of all proteins modified by LPS-cytokines and Aquamin^®^ intervention in relation to the control. Follow-up analysis involved a targeted approach towards inflammation, differentiation, barrier-related, cell-cell and cell adhesion proteins. Mass spectrometry-based proteomics data were deposited to the ProteomeXchange Consortium via the PRIDE partner repository with the dataset identifier is pending.

### Statistical analysis

Means and standard deviations were obtained for individual proteins and TEER measurements. Groups were analyzed by one-way analysis of variance (ANOVA) followed by unpaired t-test (two-tailed) using GraphPad Prism (v 9.1). Pathways enrichment data reflect Reactome-generated p-values based on the number of entities identified in a given pathway as compared to total proteins involved in that pathway. A binomial test was applied within Reactome to calculate the probability shown for each result, and the p-values were corrected for the multiple testing (Benjamini– Hochberg procedure) that arises from evaluating the submitted list of identifiers against every pathway. Data were considered significant at p<0.05.

## RESULTS

### Effects of LPS-cytokine mix, Aquamin^®^ and the combination of the two interventions on organoid morphology

Colon organoids from three subjects were incubated for a two-week period (with subculture at the end of week-one) under control conditions (i.e., in 25% LWRN – KGM Gold medium alone) or in the same medium treated either with LPS and the mix of pro-inflammatory cytokines, with Aquamin^®^ alone or with a combination of both interventions. At the beginning of the treatment phase, and also immediately after the first subculture, organoids were approximately 40 μm in size (on average). At the time of harvest (either after the first week or at the end of the 14-day culture period), individual organoids had increased to approximately 500 μm in diameter. Phase-contrast images taken at the end of the two-week incubation period identified no gross morphological changes attributable to either intervention alone or with the combination of interventions. Individual organoids appeared as thick-walled round or oval-shaped structures. Some of the structures were multi-lobed (Figure 1).

**Figure 1.**
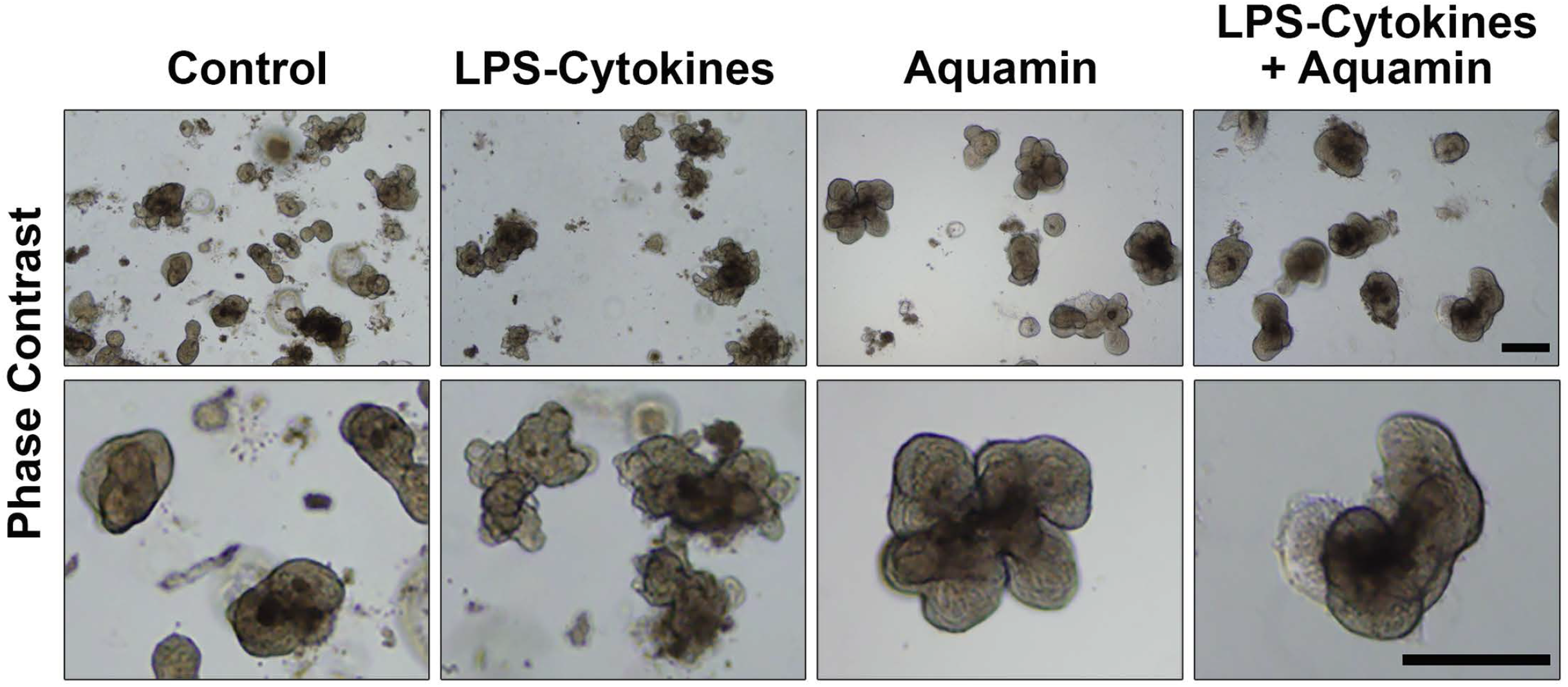
Colon organoid appearance assessed by phase-contrast microscopy: At the end of the incubation period, intact colon organoids were examined by phase-contrast microscopy. Organoids were present as thick-walled structures with few surface buds. A wide range of sizes and shapes were seen under all conditions. Images in the upper panels presented at lesser magnification and the lower panels present higher magnification. Scale bars=500μm.

### Effects of the LPS-cytokine mix, Aquamin^®^ and the combination of the two interventions on proteomic signature: Overview

At the end of the two-week incubation period, protein was prepared from each of the three treatment groups (n=3 subjects evaluated separately) and evaluated by proteomic analysis. From these separate evaluations, a total of 4700 unique proteins was identified with <2% FDR. Using 1.8-fold difference from control and <2% FDR as criteria, these studies identified a total of 219 proteins altered with the LPS-cytokine mix, 869 proteins altered with Aquamin^®^ alone and 457 proteins altered with the combination of interventions (Figure 2). Not surprisingly, the two interventions were very different in the proteins affected (upper Venn plot). Even among the common proteins, there was little overlap between the two interventions (upper correlation curve). In contrast, when proteins altered in response to the LPS-cytokine mix with or without Aquamin^®^ were compared, there was significant overlap (middle Venn plot and correlation curve). Likewise, when proteins altered in response to Aquamin^®^ treatment with or without the LPS-cytokine mix were compared, significant overlap was observed (lower Venn plot and correlation curve).

**Figure 2.**
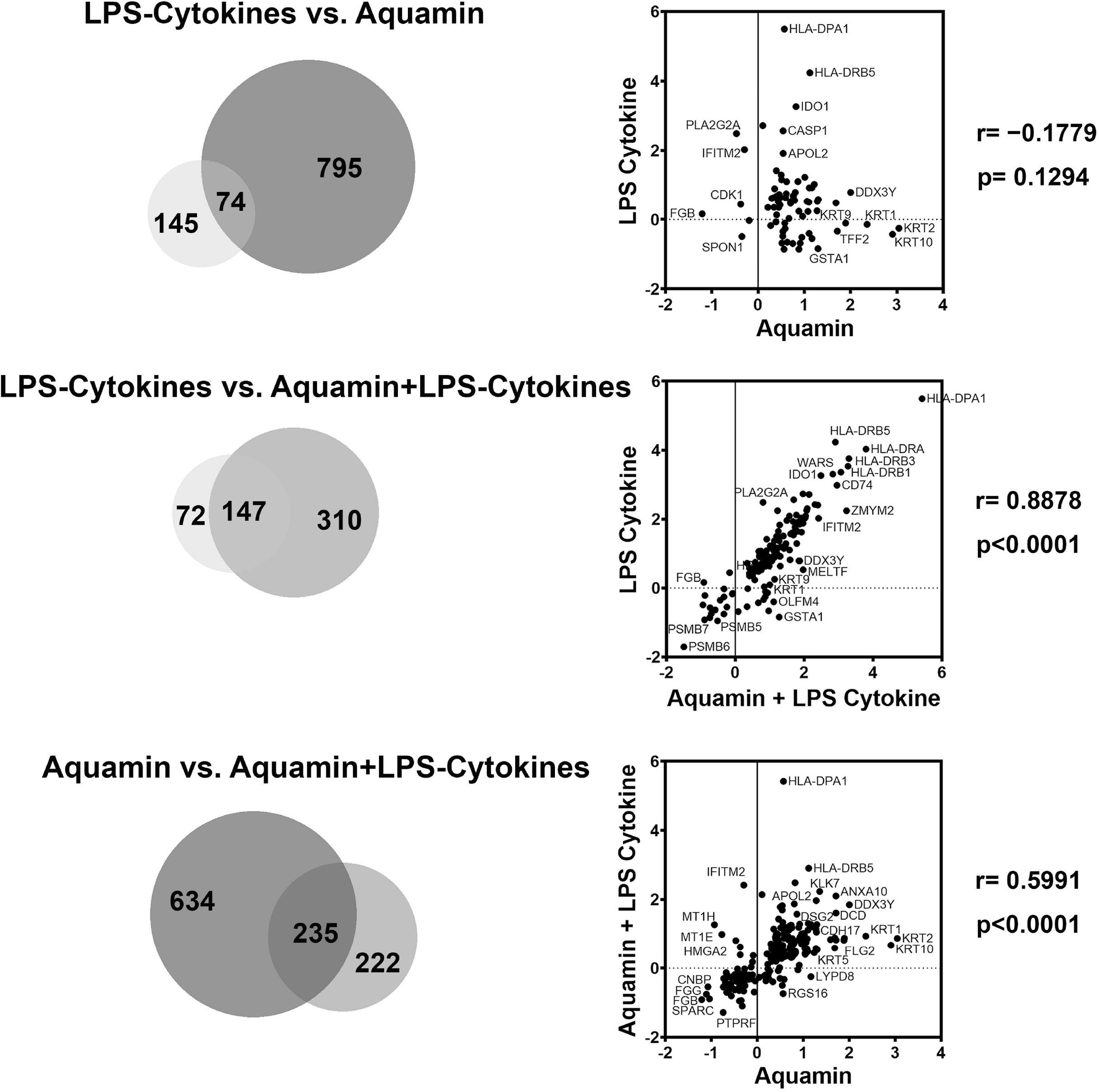
Proteomic analysis of human colon organoids: Effects of the LPS-cytokine mix and Aquamin^®^ alone and in combination. At the end of the incubation period, lysates were subjected to TMT mass spectroscopy-based proteomic analysis. <u>Upper left</u>: Venn plot showing proteins altered (increased or decreased) by an average of 1.8-fold or greater, comparing the LPS-cytokine mix with Aquamin^®^. The data are based on all unique proteins identified across organoid cultures from three separate specimens with each of the two interventions compared to control. <u>Upper right</u>: Correlation analysis showing relative expression levels of the seventy-four proteins common to both interventions. <u>Middle left</u>: Venn plot comparing proteins altered by the LPS-cytokine mix alone to those altered by the LPS-cytokine mix in the presence of Aquamin^®^. <u>Middle right</u>: Correlation analysis showing relative expression levels of the 147 proteins common to both interventions. <u>Lower left</u>: Venn plot comparing proteins altered in the presence of Aquamin^®^ alone to the proteins altered by Aquamin^®^ in combination with the LPS-cytokine mix. <u>Lower right</u>: Correlation analysis showing relative expression levels of the 235 proteins common to both interventions.

After evaluating each specimen separately, data from the three specimens were merged to obtain mean abundance ratio values. Using the same criteria (1.8-fold change from control with <2% FDR), a total of 94 proteins were identified as responsive to the pro-inflammatory mix while 91 proteins were responsive to Aquamin^®^ (alone) and 145 proteins were responsive to the combination of these two interventions (Figure 3). The majority of proteins were up-regulated (independent of intervention) with only a small fraction of the total being down-regulated. Venn plots show the minimal overlap among the three treatment groups. The complete list of common proteins and proteins unique to each group is presented in Supplement Table 3.

**Figure 3.**
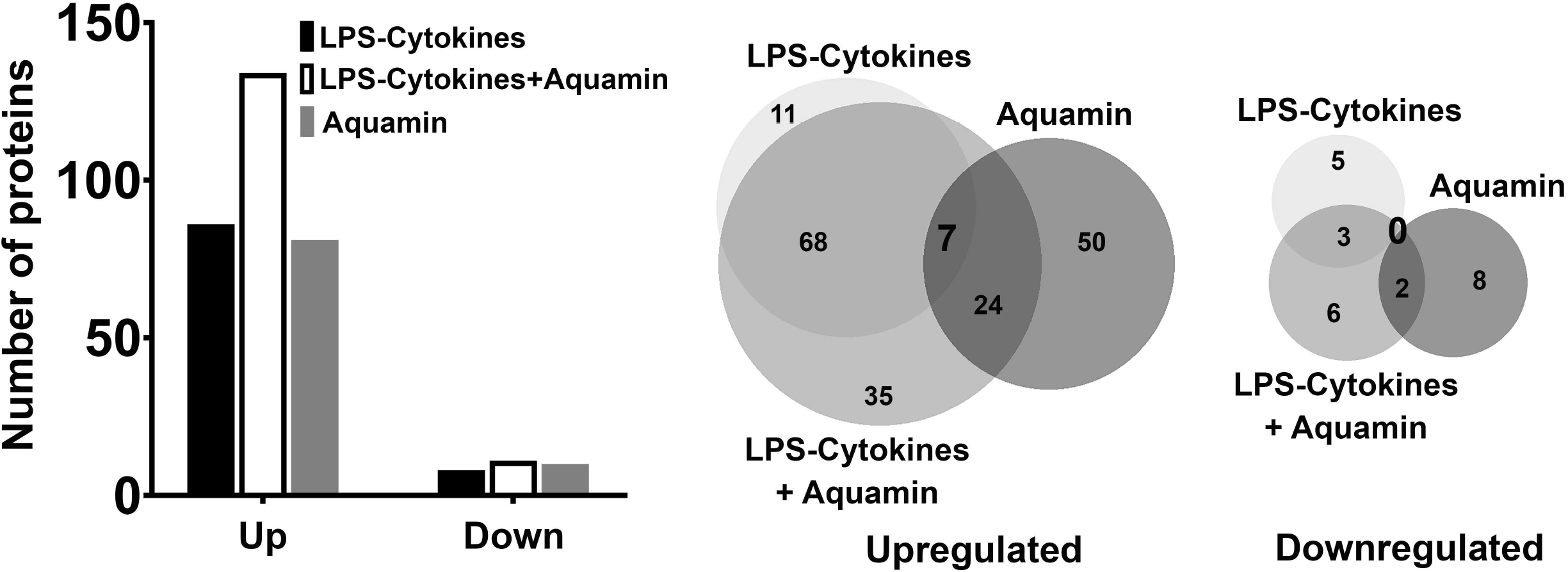
Proteomic analysis of human colon organoids exposed to the LPS-cytokine mix, Aquamin^®^ and the combination of both interventions: Data from three subjects merged. Proteomic data from the three separate colon organoid cultures (analyzed in Figure 2 above) were merged to obtain mean values and re-analyzed. <u>Left</u>: Bar graph showing proteins (merged data) increased or decreased by an average of 1.8-fold or greater with each of the three interventions compared to control. <u>Middle and right</u>: Venn plots showing overlap among the three interventions.

### Effects of the LPS-cytokine mix on proteomic signature: Alone and in combination with Aquamin^®^

Table 1 provides the complete list of the 86 individual proteins up-regulated in response to the LPS-cytokines stimulus. Table 2 is a list of relevant pathways most affected by the protein changes. The complete list of 182 pathways affected by this pro-inflammatory stimulus (with p<0.05) is presented in Supplement Table 4. Among the most highly up-regulated proteins were several HLA class II histocompatibility antigens along with certain HLA class I antigens. Additional proteins that support antigen presentation, inflammation, cytokine signaling, PD-1 signaling and immune cell recognition were induced along with HLA moieties. Not surprisingly, up-regulation of immune cell signaling pathways accompanied these changes.

**Table 1.**
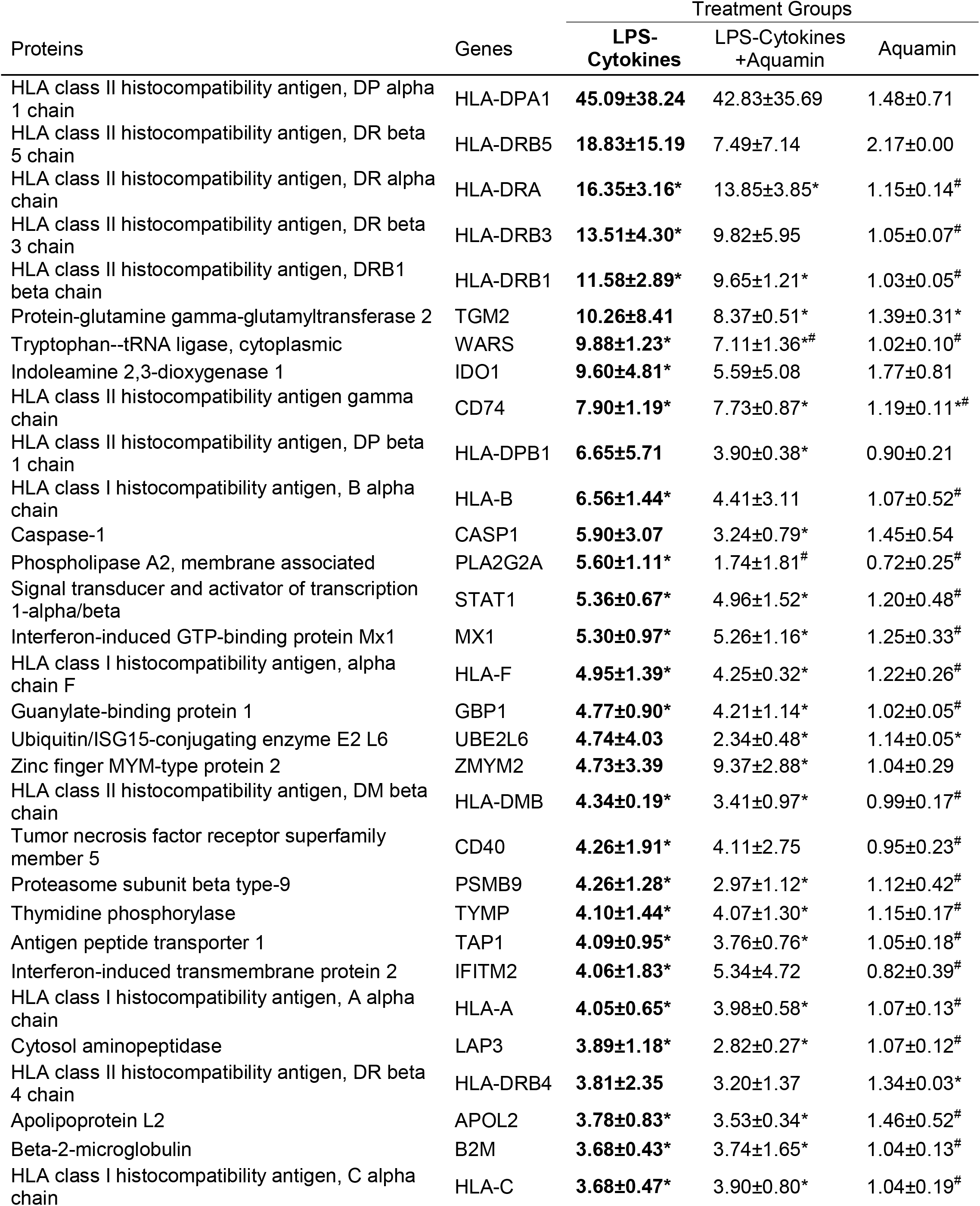
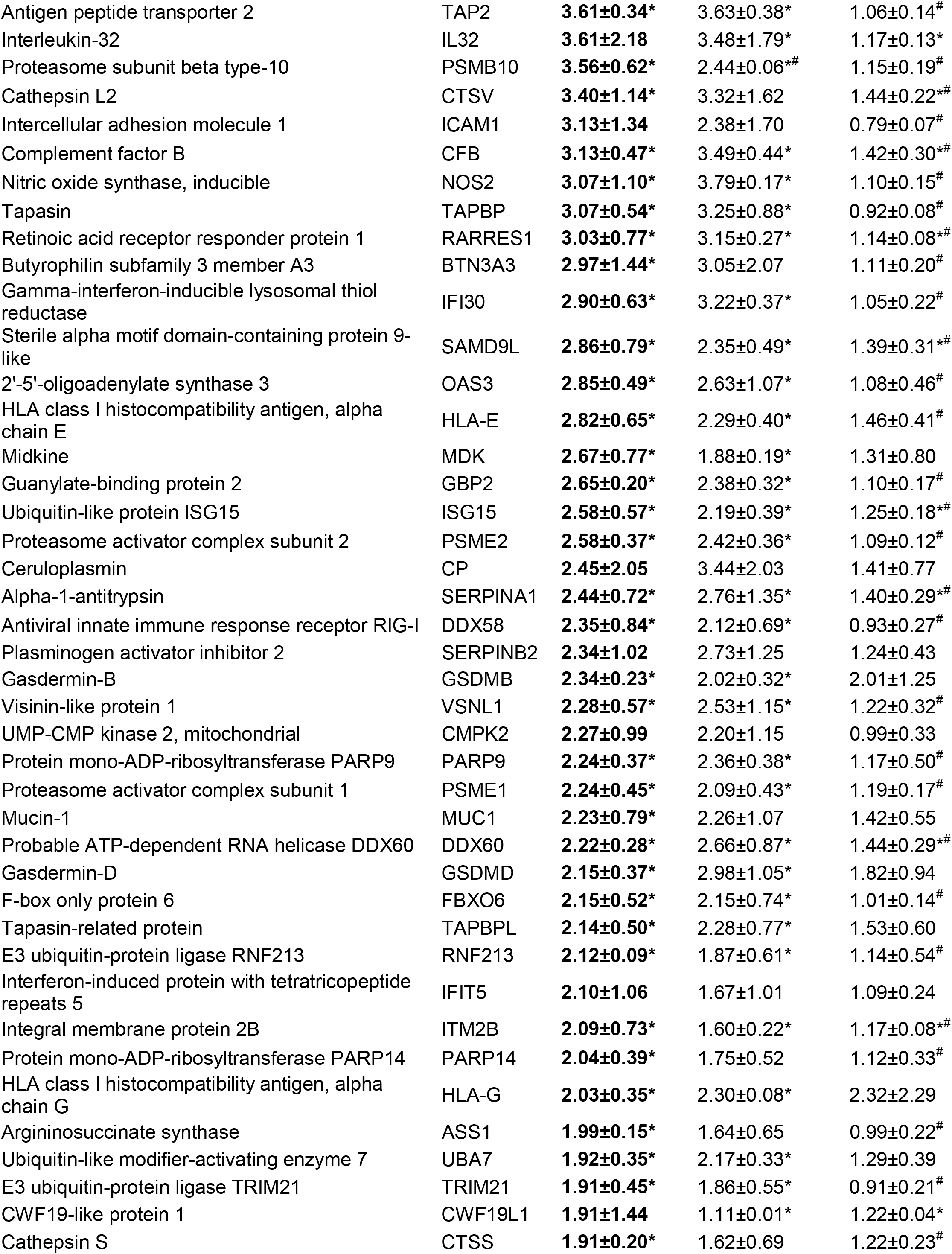
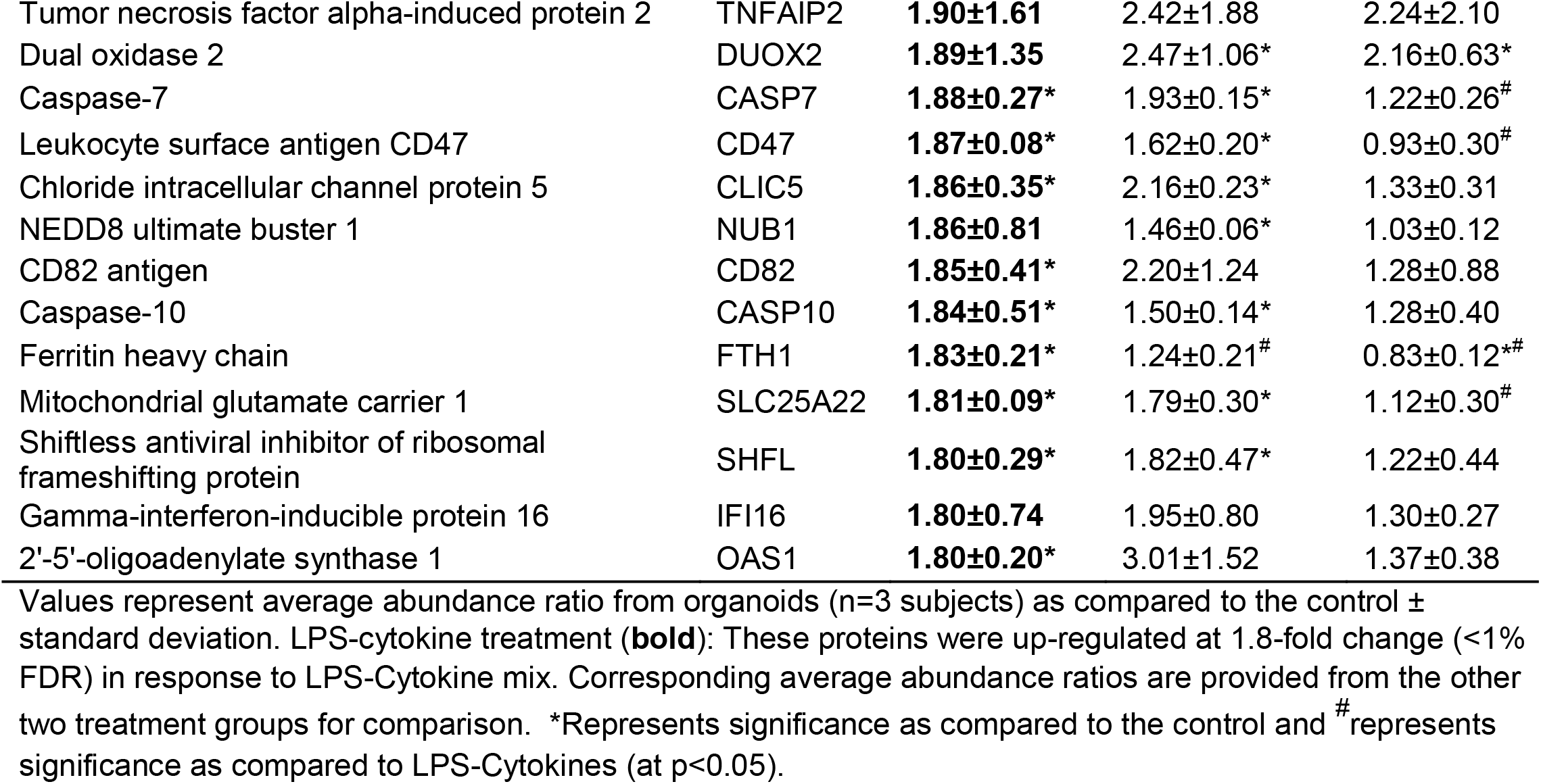
Up-regulated proteins: The effect of LPS-Cytokines on the proteomic expression.

**Table 2.** Top pathways associated with up-regulated proteins altered with LPS-Cytokines.

| Pathways | Entities pValue |
| --- | --- |
| Interferon Signaling | $1.11 \times 10^{-16}$ |
| Interferon gamma signaling | $1.11 \times 10^{-16}$ |
| Immune System | $1.11 \times 10^{-16}$ |
| Cytokine Signaling in Immune system | $1.11 \times 10^{-16}$ |
| Adaptive Immune System | $6.44 \times 10^{-15}$ |
| Interferon alpha/beta signaling | $1.90 \times 10^{-14}$ |
| Antigen processing-Cross presentation | $4.80 \times 10^{-13}$ |
| ER-Phagosome pathway | $2.57 \times 10^{-11}$ |
| Class I MHC mediated antigen processing & presentation | $2.58 \times 10^{-11}$ |
| MHC class II antigen presentation | $1.11 \times 10^{-9}$ |
| Downstream TCR signaling | $1.76 \times 10^{-9}$ |
| Antigen Presentation: Folding, assembly and peptide loading of class I MHC | $5.08 \times 10^{-9}$ |
| TCR signaling | $1.04 \times 10^{-8}$ |
| Endosomal/Vacuolar pathway | $1.21 \times 10^{-8}$ |
| Translocation of ZAP-70 to Immunological synapse | $2.19 \times 10^{-8}$ |
| Phosphorylation of CD3 and TCR zeta chains | $3.66 \times 10^{-8}$ |
| PD-1 signaling | $4.30 \times 10^{-8}$ |
| Antiviral mechanism by IFN-stimulated genes | $4.84 \times 10^{-7}$ |
| Innate Immune System | $6.93 \times 10^{-7}$ |
| Signaling by Interleukins | $6.13 \times 10^{-5}$ |
| Cross-presentation of soluble exogenous antigens (endosomes) | $6.27 \times 10^{-4}$ |
| Regulation of ornithine decarboxylase (ODC) | $6.75 \times 10^{-4}$ |
| p53-Independent G1/S DNA damage checkpoint | $7.25 \times 10^{-4}$ |
| TNFR2 non-canonical NF-kB pathway | 0.001 |
| Dectin-1 mediated noncanonical NF-kB signaling | 0.001 |
| Interleukin-1 family signaling | 0.001 |
| Defective CFTR causes cystic fibrosis | 0.001 |
| Programmed Cell Death | 0.001 |
| Interleukin-1 processing | 0.002 |
| Cell recruitment (pro-inflammatory response) | 0.019 |
| The AIM2 inflammasome | 0.023 |
| Beta-catenin independent WNT signaling | 0.025 |
| The IPAF inflammasome | 0.038 |

Several proteins involved in intracellular protein processing / breakdown (degradative enzymes and oxidant generators) were also responsive to the LPS-cytokine mix, as were several moieties involved in ubiquitinating reactions and multiple proteasome subunits. Of interest, the most highly up-regulated proteasome subunits (PSMB9, PSMB10, PSME2 and PSME1) are found in the specialized proteasome complex known as the immunoproteasome (29), which is responsible for modification of peptides for binding by major histocompatibility complex molecules and presentation to T cells. These proteins are also involved in ornithine decarboxylase up-regulation. Increased colonic ornithine decarboxylase activity occurs with mucosal inflammation (30) and is thought to play roles in both injury and repair. Interestingly, mucin-1 (muc1), a cell surface mucin was upregulated in response to LPS-cytokines. It has been shown previously that pro-inflammatory cytokines increase muc1 expression in the oral mucosa as part of the host response (31).

For comparison, Table 1 also shows how the concomitant presence of Aquamin^®^ along with the LPS-cytokine mix affected the same eighty-six proteins. As seen from the table, the majority of proteins that responded to the LPS-cytokine mix alone also responded to the pro-inflammatory challenge when Aquamin^®^ was present. This suggests that Aquamin^®^ does not act, primarily, to suppress protein changes that occur in response to LPS-cytokine stimulation. In further support for this idea, pathway analysis conducted using proteins up-regulated by the LPS-cytokine mix in the presence of Aquamin^®^ demonstrated a similar spectrum of target pathways affected as seen with the LPS-cytokine mix alone. Specifically, a total of 175 pathways (p<0.05; Supplement Table 5) were likely to be affected by the concomitant presence of Aquamin^®^ along with the LPS-cytokine mix, compared to 182 pathways affected by the LPS-cytokine mix alone (Table 2 above and Supplement Table 4). Among these, 144 pathways were common between the two groups. Although the overall effect of Aquamin^®^ on organoid response to LPS-cytokine stimulation was not widespread, there was some effect. Specifically, among the LPS-cytokine up-regulated proteins, twenty-seven were inhibited by at least 20% when Aquamin^®^ was concomitantly present. Some of these proteins play a role in modifying the pro-inflammatory environment (See following sections for detailed analysis).

Finally, Table 1 shows how the same 86 (LPS-cytokine stimulated) proteins responded to Aquamin^®^ in the absence of the pro-inflammatory stimulus. As predicted from Figures 2 and 3 above, there was little effect. Although the majority of LPS-cytokines-responsive proteins showed minimal change with Aquamin^®^ alone, there were a few important exceptions. These included TNF-α – induced protein 2 (TNFAIP2) and dual oxidase 2 (an NADPH-like protein found in non-phagocytic cells) as well as two members of the gasdermin family (GSDMB and GSDMD). The significance of the overlap in this small group of mostly unrelated proteins is unclear. While each of these moieties has multiple biological roles, all four have been suggested to participate in processes that counteract inflammation (32–35).

Table 3 presents the eight proteins down-regulated (≥1.8-fold and <2% FDR) in response to the LPS-cytokines stimulus. How these proteins responded to Aquamin^®^ alone or when Aquamin^®^ was provided along with the pro-inflammatory mix is presented for comparison. The three most highly down-regulated proteins were proteasome subunits; all three of these proteins are present in the “classical” proteasome complex (29); their down-regulation is consistent with their replacement by analogues found in the immunoproteasome (Table 1 above). These proteins were not altered with Aquamin^®^ alone, and Aquamin^®^ did not affect their response to the LPS-cytokine mix. Another down-regulated protein shown in Table 3 was glutathione S-transferase A1 (GSTA1). This protein is part of the glutathione antioxidant mechanism and a critical component of detoxification pathways (36). It is of interest that Aquamin^®^ alone substantially up-regulated GSTA1 as compared to control, and when Aquamin^®^ was present along with the pro-inflammatory stimulus, the GSTA1 level was still strongly up-regulated. Supplement Table 6 provides a list of 160 pathways (with p<0.05) likely to be affected by proteins down-regulated with the pro-inflammatory mix.

**Table 3.** Down-regulated proteins: The effect of LPS-Cytokines on the proteomic expression.

| Proteins | Genes | Treatment Groups |  |  |
| --- | --- | --- | --- | --- |
|  |  | LPS-Cytokines | LPS-Cytokines + Aquamin | Aquamin |
| Proteasome subunit beta type-6 | PSMB6 | **0.31±0.28\*** | 0.36±0.20* | 1.03±0.22<sup>#</sup> |
| Proteasome subunit beta type-5 | PSMB5 | **0.52±0.40** | 0.70±0.35 | 1.01±0.03<sup>#</sup> |
| Proteasome subunit beta type-7 | PSMB7 | **0.53±0.25\*** | 0.54±0.20* | 0.99±0.18<sup>#</sup> |
| Antileukoproteinase | SLPI | **0.55±0.37\*** | 0.60±0.32* | 0.91±0.27 |
| Regulator of G-protein signaling 16 | RGS16 | **0.55±0.28\*** | 0.60±0.27* | 1.48±0.88 |
| Elafin | PI3 | **0.55±0.07\*** | 0.46±0.02* | 1.17±0.34<sup>#</sup> |
| Phosphoenolpyruvate carboxykinase, cytosolic [GTP] | PCK1 | **0.55±0.24\*** | 1.10±0.42 | 1.84±1.02<sup>#</sup> |
| Glutathione S-transferase A1 | GSTA1 | **0.56±0.24\*** | 2.42±3.12 | 2.45±1.87 |

Values represent average abundance ratio from organoids (n=3 subjects) as compared to the control ± standard deviation. LPS-cytokine treatment (**bold**): These proteins were down-regulated at 1.8-fold change (<1% FDR) in response to LPS-Cytokine mix. Corresponding average abundance ratios are provided from the other two treatment groups for comparison. \*Represents significance as compared to the control and <sup>#</sup>represents significance as compared to LPS-Cytokines (at p<0.05).

### Effects of Aquamin^®^ on proteomic signature: Alone and in combination with the LPS-cytokine mix

In addition to assessing the effects of the LPS-cytokine mix on the colon organoid proteomic signature and determining how the concomitant presence of Aquamin^®^ modified these effects, the inverse analysis was also carried out. That is, Aquamin^®^ was examined (independently) for ability to modify the proteome. In parallel, how concomitant exposure to the LPS-cytokine mix modified the response to the multi-mineral intervention was determined. Table 4 shows the individual proteins up-regulated in response to Aquamin^®^ alone (≥1.8-fold and <2% FDR) and the effects of the LPS-cytokine mix (alone and in the presence of Aquamin^®^, respectively) on these same proteins. Consistent with what we have reported previously in colon organoids derived from tissue of healthy individuals (20) or from UC lesional tissue (22), Aquamin^®^ alone caused up-regulation of multiple proteins involved in epithelial differentiation, tissue strength and barrier formation. Included were several keratins and other differentiation-related proteins. Two signature proteins related to epithelial cell cohesion and tissue strength (i.e., desmoglein-2 and cadherin-17) were strongly up-regulated by the multi-mineral product as was trefoil factor-2 (component of the mucinous layer) and carcinoembryonic antigen related cell adhesion molecule-5 (CEACAM-5). The most important pathways associated with Aquamin^®^-induced protein up-regulation are presented in Supplement Table 7. Not surprisingly, these included differentiation, cell-cell junctional complexes and tissue barrier-related pathways.

**Table 4.**
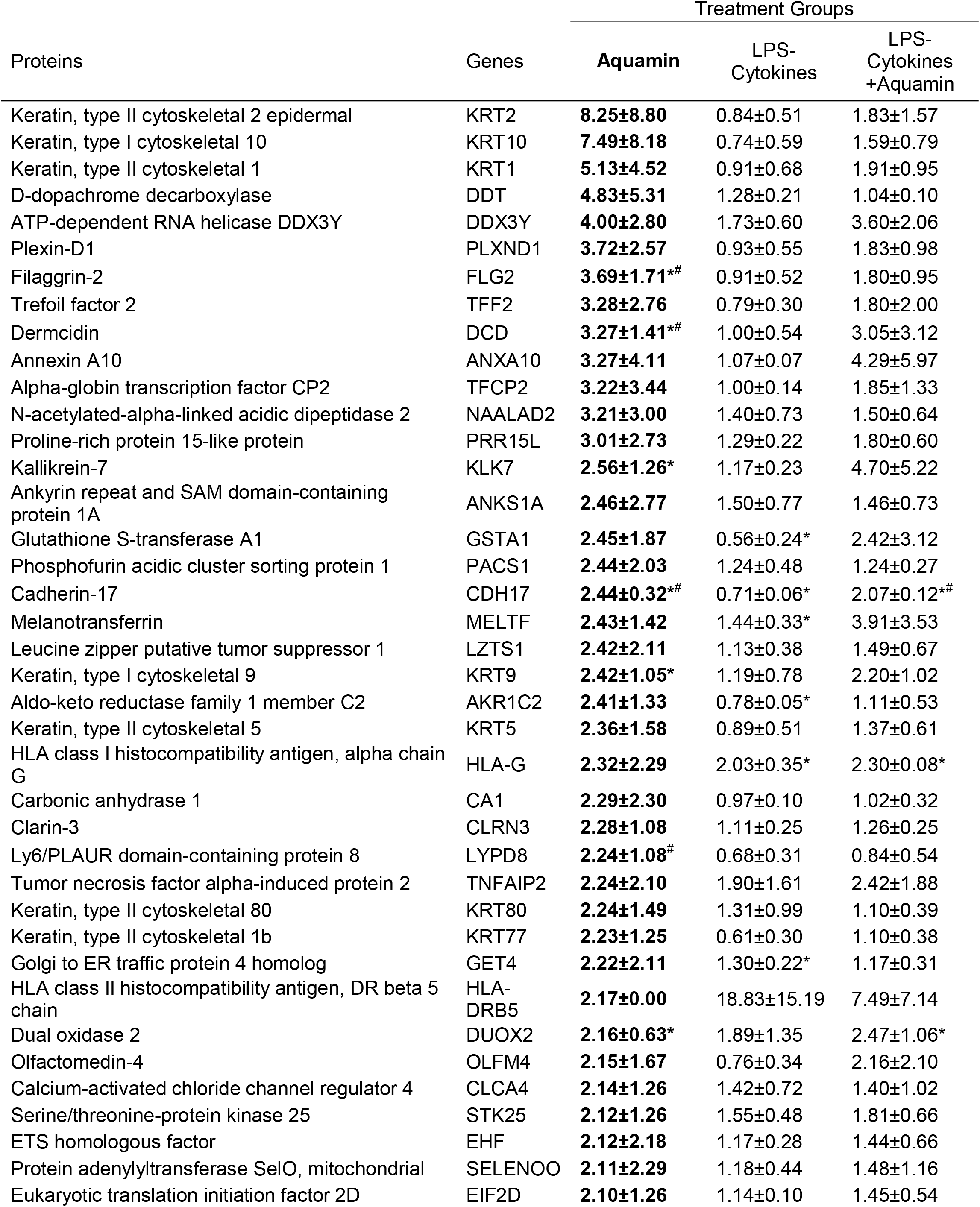
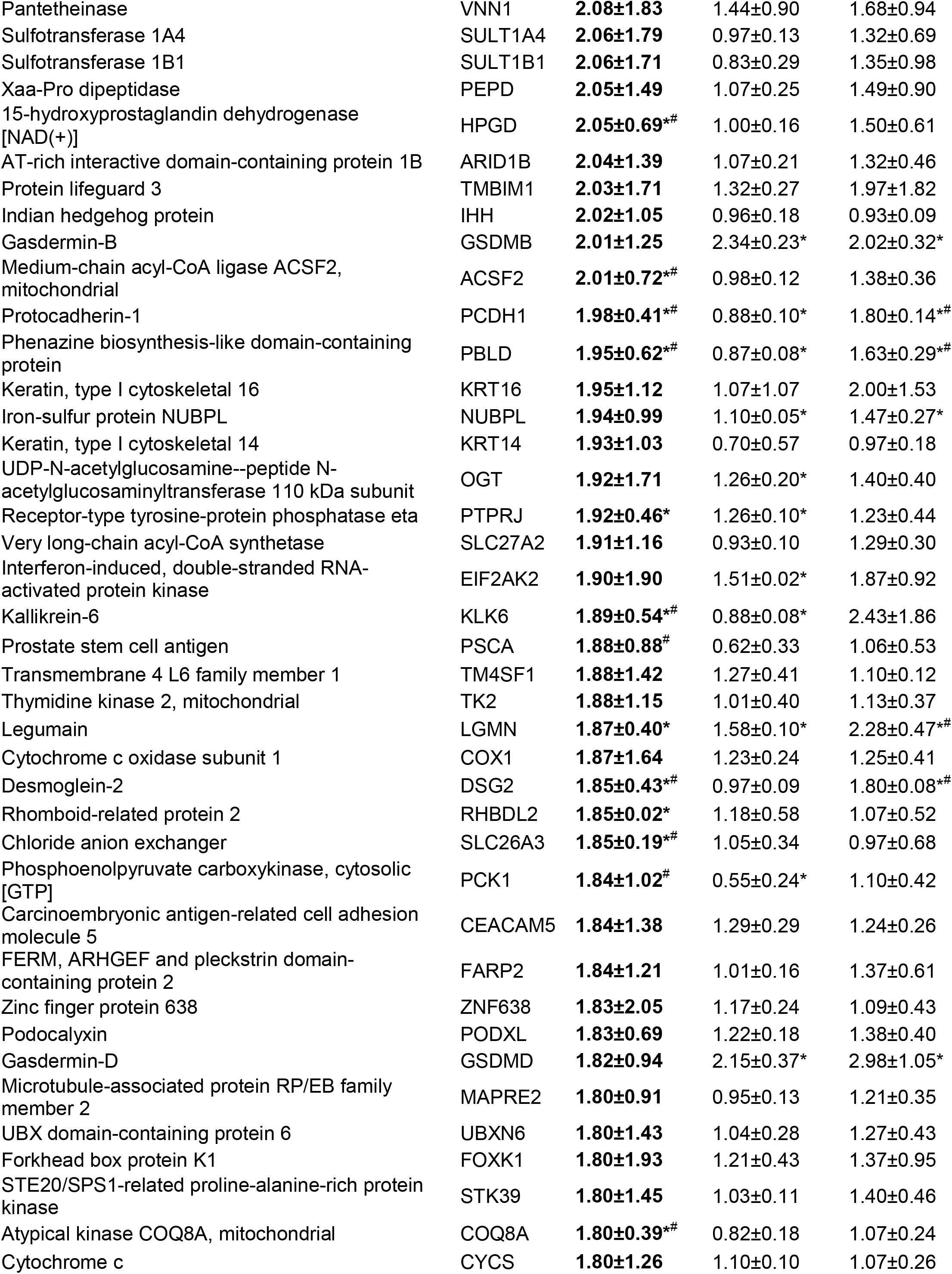
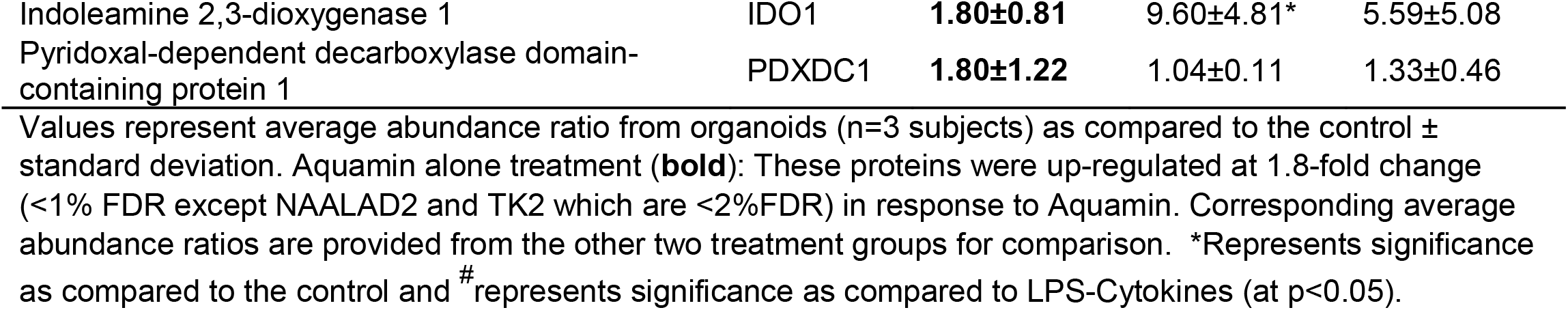
Up-regulated proteins: The effect of Aquamin alone on the proteomic expression.

As also seen in Table 4, virtually none of the differentiation and barrier proteins were up-regulated in response to the LPS-cytokines treatment alone. Several, in fact, demonstrated decreased expression, consistent with the barrier-disrupting effects of inflammation. More importantly, Table 4 shows that even in the presence of the pro-inflammatory stimulus, Aquamin^®^ treatment stimulated expression of proteins critical to differentiation and barrier formation.

To obtain a more complete picture of how the multi-mineral supplement altered differentiation and barrier protein expression and how the concomitant presence of the LPS-cytokine mix may or may not have modified this, we searched the proteomic database specifically for proteins involved in differentiation, tissue strength and barrier function. Proteins of relevance are presented in Figure 4. In addition to the proteins detected in the nonbiased search (Table 4 above), several additional keratins and other differentiation-related proteins were identified along with a number of adherens junctional proteins (cadherin family members), desmosomal proteins, tight junctional proteins, trefoils, mucins and CEACAM components. Consistent with our past reports (20,22), multiple cadherins and desmosomal proteins demonstrated strong up-regulation with Aquamin^®^. Also consistent, most of the tight junctional proteins changed little in response to Aquamin^®^, although claudin 4 and ZO1 (TJP1) were modestly up-regulated. Most importantly, Figure 4 demonstrates that the concomitant presence of the LPS-cytokine mix along with Aquamin^®^ had virtually no dampening effect on the up-regulated proteins including those contributing to the formation of adherens junctions and desmosomes.

**Figure 4.**
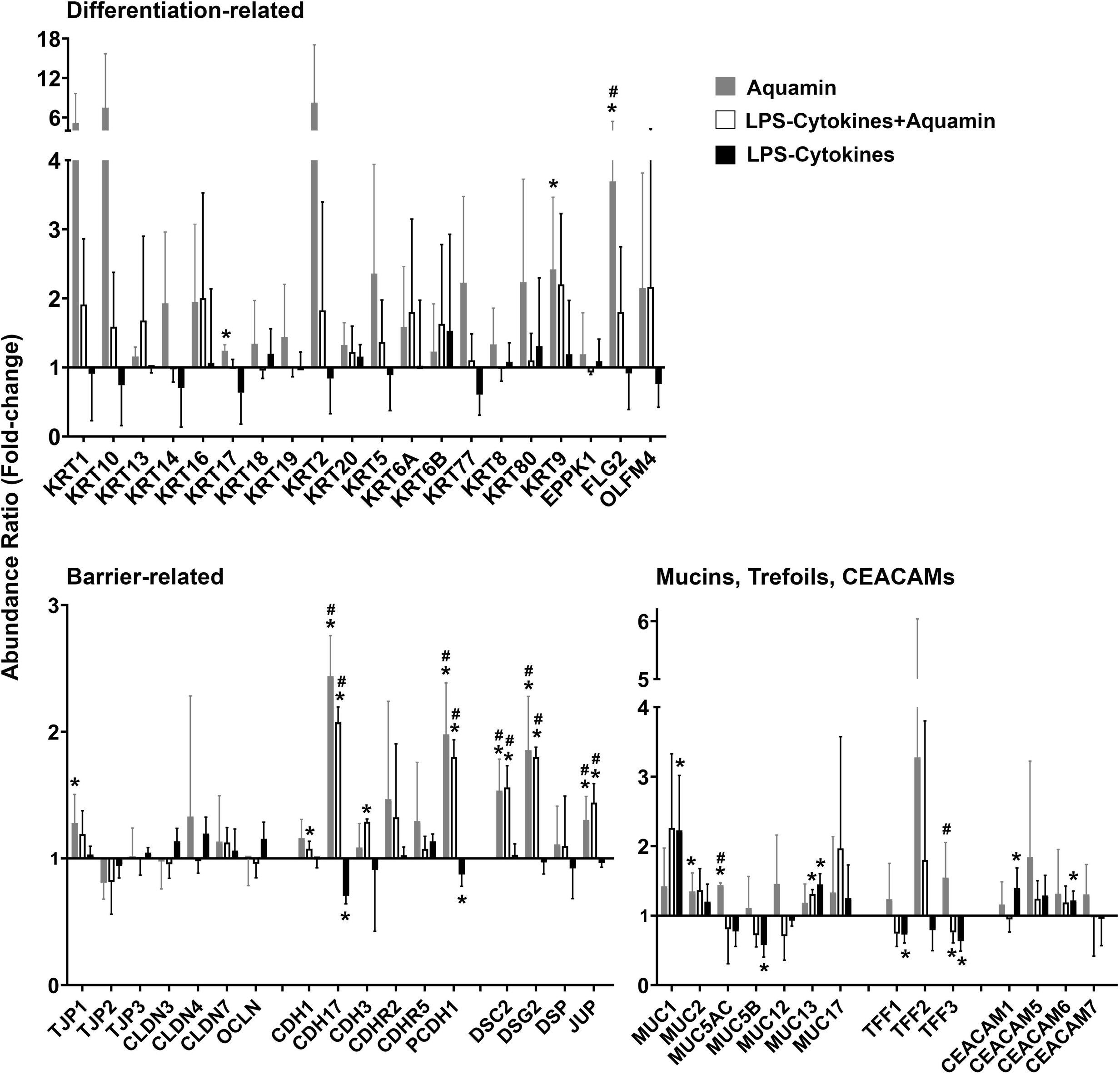
Directed search for proteins involved in differentiation and barrier formation: Effects of Aquamin^®^ alone compared to Aquamin^®^ in the presence of the LPS-cytokine mix. Bar graphs showing the up- or down-regulation of individual proteins of interest (merged data) with each of three interventions compared to control (independent of fold-change but with <2% FDR). Values shown are means and standard deviations. * represents significance (p<0.05) relative to the control and # represents significance (p<0.05) relative to the corresponding LPS-cytokine mix.

In addition to proteins up-regulated in response to Aquamin, 10 down-regulated proteins (nonbiased search; ≥1.8-fold with <2% FDR) were also identified (Table 5A). Effects of LPS-cytokine treatment (alone and in the presence of Aquamin^®^) are presented in the same table for comparison. Among the ten proteins were two of the three peptides that make up intact fibrinogen and the protein, SPARC. None of the three proteins was altered by the LPS-cytokine mix alone and the pro-inflammatory mix did not affect responses to the multi-mineral product. Both fibrinogen (37,38) and SPARC (39) have been shown to promote chronic inflammation, including in the gastrointestinal tract. Their down-regulation by Aquamin^®^ suggests the possibility of an anti-inflammatory effect separate from barrier improvement. A list of pathways altered by the down-regulated moieties with Aquamin^®^ is presented in Table 5B.

**Table 5A.**
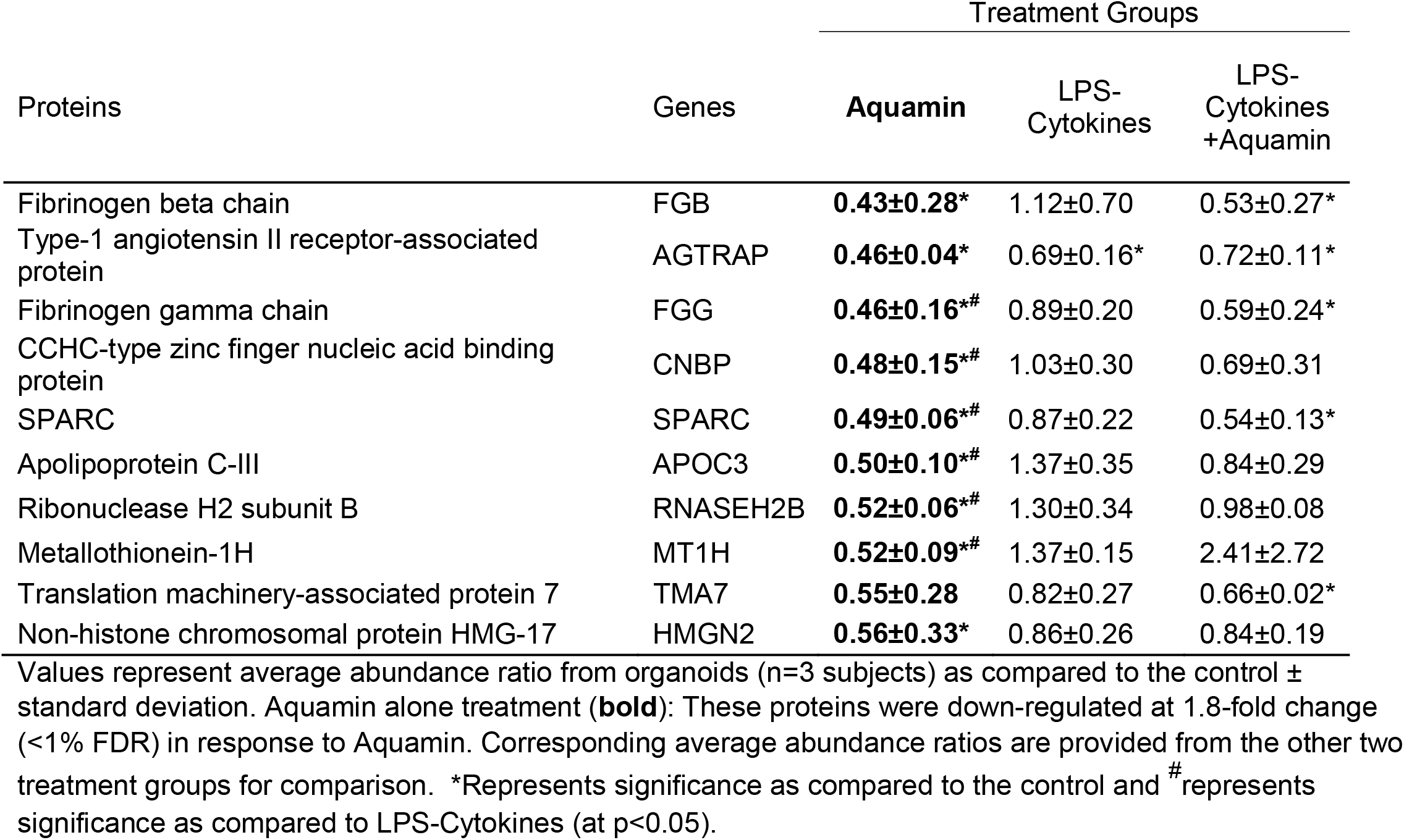
Down-regulated proteins: The effect of Aquamin alone on the proteomic expression.

**Table 5B.** Pathways associated with down-regulated proteins altered with Aquamin.

| Pathways | Entities<br>pValue | Mapped entities |
| --- | --- | --- |
| Signaling by BRAF and RAF1 fusions | 2.14×10<sup>-5</sup> | AGTRAP;FGG;FGB |
| Oncogenic MAPK signaling | 4.37×10<sup>-5</sup> | AGTRAP;FGG;FGB |
| p130Cas linkage to MAPK signaling for integrins | 7.46×10<sup>-5</sup> | FGG;FGB |
| GRB2:SOS provides linkage to MAPK signaling for Integrins | 7.46×10<sup>-5</sup> | FGG;FGB |
| MyD88 deficiency (TLR2/4) | 1.19×10<sup>-4</sup> | FGG;FGB |
| IRAK4 deficiency (TLR2/4) | 1.32×10<sup>-4</sup> | FGG;FGB |
| Regulation of TLR by endogenous ligand | 1.46×10<sup>-4</sup> | FGG;FGB |
| Platelet degranulation | 1.52×10<sup>-4</sup> | FGG;SPARC;FGB |
| Common Pathway of Fibrin Clot Formation | 1.60×10<sup>-4</sup> | FGG;FGB |
| Response to elevated platelet cytosolic Ca2+ | 1.70×10<sup>-4</sup> | FGG;SPARC;FGB |
| Integrin signaling | 2.58×10<sup>-4</sup> | FGG;FGB |
| Diseases of Immune System | 3.80×10<sup>-4</sup> | FGG;FGB |
| Diseases associated with the TLR signaling cascade | 3.80×10<sup>-4</sup> | FGG;FGB |
| Signaling by high-kinase activity BRAF mutants | 4.49×10<sup>-4</sup> | FGG;FGB |
| Formation of Fibrin Clot (Clotting Cascade) | 4.99×10<sup>-4</sup> | FGG;FGB |
| Platelet Aggregation (Plug Formation) | 5.24×10<sup>-4</sup> | FGG;FGB |
| MAP2K and MAPK activation | 5.50×10<sup>-4</sup> | FGG;FGB |
| Signaling by RAF1 mutants | 5.77×10<sup>-4</sup> | FGG;FGB |
| Paradoxical activation of RAF signaling by kinase inactive BRAF | 7.21×10<sup>-4</sup> | FGG;FGB |
| Signaling by moderate kinase activity BRAF mutants | 7.21×10<sup>-4</sup> | FGG;FGB |
| Signaling downstream of RAS mutants | 7.21×10<sup>-4</sup> | FGG;FGB |
| Signaling by RAS mutants | 7.21×10<sup>-4</sup> | FGG;FGB |
| Platelet activation, signaling and aggregation | 0.001 | FGG;SPARC;FGB |
| Extracellular matrix organization | 0.002 | FGG;SPARC;FGB |
| Integrin cell surface interactions | 0.002 | FGG;FGB |
| Scavenging by Class H Receptors | 0.003 | SPARC |
| MyD88:MAL(TIRAP) cascade initiated on plasma membrane | 0.004 | FGG;FGB |
| Toll Like Receptor TLR6:TLR2 Cascade | 0.004 | FGG;FGB |
| Toll Like Receptor TLR1:TLR2 Cascade | 0.004 | FGG;FGB |
| Toll Like Receptor 2 (TLR2) Cascade | 0.004 | FGG;FGB |
| Diseases of signal transduction by growth factor receptors and second messengers | 0.006 | AGTRAP;FGG;FGB |
| Toll Like Receptor 4 (TLR4) Cascade | 0.007 | FGG;FGB |
| ER-Phagosome pathway | 0.008 | FGG;FGB |
| Chylomicron remodeling | 0.009 | APOC3 |
| Chylomicron assembly | 0.009 | APOC3 |
| Toll-like Receptor Cascades | 0.009 | FGG;FGB |
| HDL remodeling | 0.009 | APOC3 |
| Metallothioneins bind metals | 0.009 | MT1H |
| Antigen processing-Cross presentation | 0.010 | FGG;FGB |
| Response to metal ions | 0.012 | MT1H |
| Plasma lipoprotein assembly | 0.016 | APOC3 |
| Hemostasis | 0.021 | FGG;SPARC;FGB |
| RAF/MAP kinase cascade | 0.025 | FGG;FGB |
| MAPK1/MAPK3 signaling | 0.026 | FGG;FGB |
| Nuclear signaling by ERBB4 | 0.029 | SPARC |
| Plasma lipoprotein remodeling | 0.030 | APOC3 |
| MAPK family signaling cascades | 0.033 | FGG;FGB |
| Retinoid metabolism and transport | 0.037 | APOC3 |
| Metabolism of fat-soluble vitamins | 0.041 | APOC3 |
| Signaling by ERBB4 | 0.052 | SPARC |

Based on the findings in the non-biased search, a directed search was carried out to identify additional Aquamin^®^-responsive proteins that may contribute to anti-inflammatory activity through one mechanism or other. Supplement Table 8 provides a list of such proteins. The proteins highlighted in the list were chosen based on i) a potential role in inflammation (previously identified in the scientific literature), ii) a beneficial response to Aquamin^®^ (either up- or down-regulation), and iii) failure of the LPS-cytokine mix to interfere with response to Aquamin^®^. Supplement File 1 provides relevant citations to justify the inclusion of these proteins, and Supplement Table 9 shows pathways (with p<0.05) sensitive to proteins affected by Aquamin^®^.

Among proteins of interest identified in this manner is membrane-associated Phospholipase A2 (PLA2G2A). This Aquamin^®^-down-regulated protein is a secreted form of the enzyme. In the gastrointestinal tract, it is up-regulated in inflammation (40), is responsible for epithelial cell damage and thought to promote tumor formation (41). Also of interest, down-regulation of fibrinogen chains by Aquamin^®^ could be expected to blunt abnormal toll-like receptor (TLR) signaling (Table 5 and Supplement Table 9). Abnormal Toll-like receptor (TLR) signaling and dysfunctional TLR may contribute to the pathogenesis of inflammatory bowel disease (42).

### Desmoglein-2, cadherin-17 and occludin assessment by confocal fluorescence microscopy: Effects of Aquamin^®^, the LPS-cytokine mix and combination of both interventions

Given the strong up-regulation of both desmoglein-2 and cadherin-17 by Aquamin^®^ in the proteomic assessment (Table 4 and Figure 4) and the resistance of both proteins to the LPS-cytokine mix, expression of both proteins was assessed by confocal fluorescence microscopy. Occludin was evaluated in parallel with both proteins. Staining results are presented in Figure 5 (occludin in red; desmoglein-2 in green; cadherin-17 in green). Desmoglein-2 was readily detected under control conditions as punctate (green) fluorescence throughout the section. Boundaries between cells could not be identified. A similar pattern of fluorescence was observed in the section from the LPS-cytokines-treated organoid culture. In contrast, Aquamin^®^ treatment dramatically increased detection of desmoglein-2. Cell-cell boundaries were clearly articulated between adjacent cells. Consistent with proteomic findings, treatment with the mix of LPS and cytokines had little effect on desmoglein-2 staining in the presence of Aquamin^®^.

**Figure 5.**
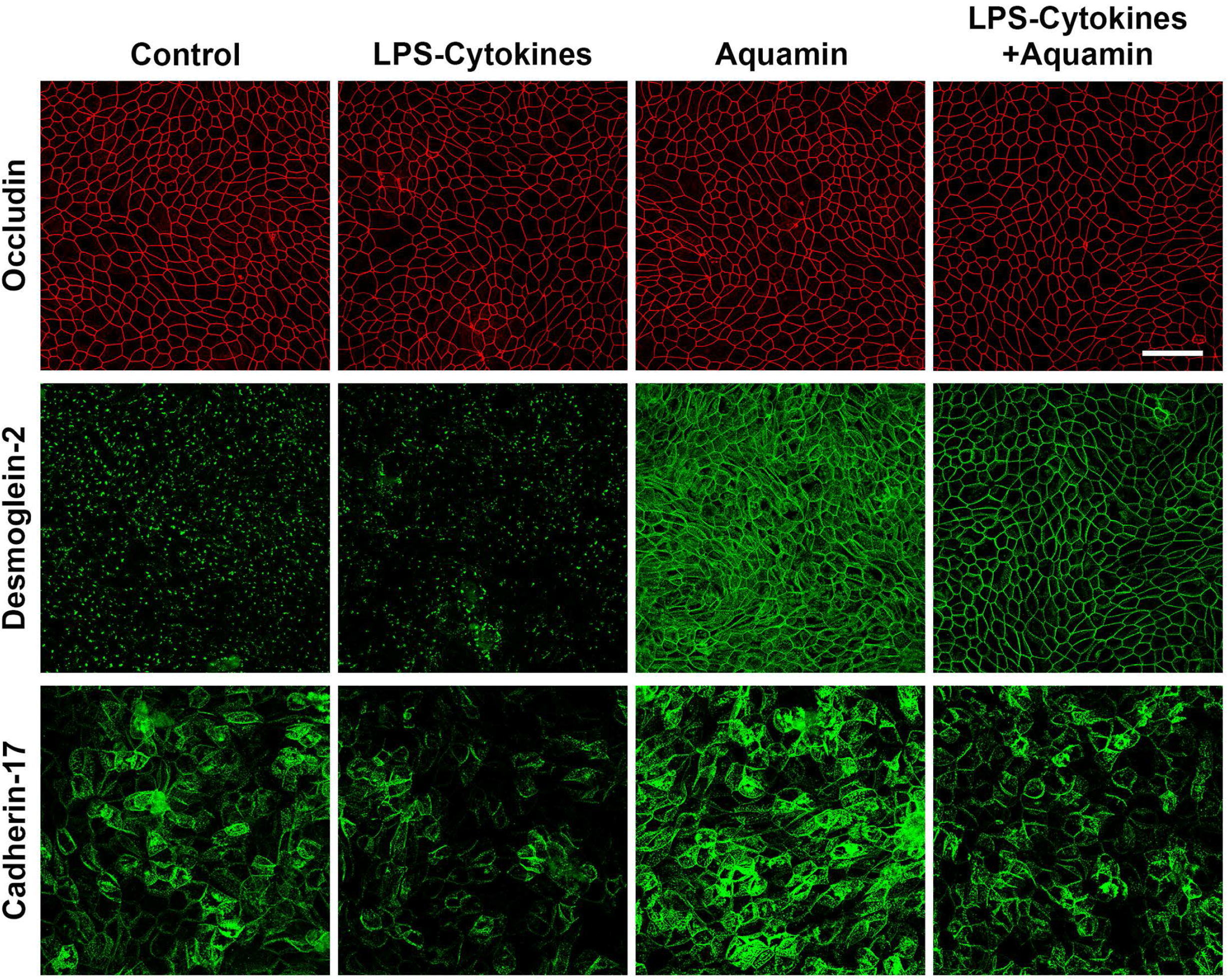
Confocal fluorescent microscopic images of occludin, desmoglein-2 and cadherin-17. Colon organoids were plated on transwell membranes and incubated under the indicated conditions. After TEER assessment at day-3, membranes were prepared and stained. Membranes were stained with a combination of antibody to occludin (red) or desmoglein-2 (green) or a combination of occludin (red) and cadherin-17 green. <u>Upper panels</u>: Occludin (max projected); <u>Middle panels</u>: Desmoglein-2 (max projected); Lower panels: Cadherin-17 (max projected). Scale bar = 60 μm.

With cadherin-17 (Figure 5), cell borders could be identified under all four treatment conditions based on the pattern of green fluorescence observed. Differences in staining intensities distinguished groups. The most intense staining was seen in sections from Aquamin^®^-treated organoids while the weakest was observed in response to the LPS-cytokine mix but without Aquamin^®^. The combination of Aquamin^®^ and the LPS-cytokine mix produced an intermediate staining intensity.

Also, consistent with current proteomic findings and previous confocal assessment (21), neither intervention modulated occludin expression significantly. Distinct cell boundaries (red fluorescence) were consistently seen under all conditions (Figure 5). Variations in intensity were slight.

### Barrier integrity (measured by TEER)

Electrical resistance across the cell layer was used as a functional barrier assay. Substantial electrical resistance across the cell layer was seen under all experimental conditions (1203 ± 752 Ohms per cm^2^ under control conditions; n=6 separate experiments) on Day 3. In the presence of LPS-cytokines treatment, electrical resistance was reduced to 62 ± 23% of control (p<0.01). As expected, based on our previous report (21), The TEER values were increased in the presence of Aquamin^®^ (125 ± 10% of control; p<0.05). The average TEER value obtained in transwell membranes exposed to the combination of interventions was 114 ± 6% of control (p<0.01 compared to LPS-cytokines alone but not significantly different compared to control or to Aquamin^®^ alone).

## DISCUSSION

Colon organoid culture has been utilized in several recent studies to help elucidate mechanisms that drive inflammation in the colon. Functional responses to cytokine stimulation have been described and associated transcriptional changes (based, primarily, on mRNA data) documented (43–46). Colon organoid cultures provide a tool not only for illuminating patho-physiological mechanisms but also provides a way to test potential interventional strategies (46). The approach taken in the present study involved exposing human colon organoids to a combination of LPS and three pro-inflammatory cytokines (TNF-α, IL-1β and IFN-γ) and then utilizing a proteomic approach to identify a broad range of protein changes that may directly or indirectly contribute to the patho-physiology of inflammatory gastrointestinal diseases. Overall, a total of 219 individual proteins were identified with a ≥1.8-fold expression level change in response to the inflammatory challenge as compared to control. Among the most highly up-regulated proteins were HLA class I and II histocompatibility antigens along with additional proteins that support antigen presentation and immune cell recognition. Up-regulation of immune cell signaling molecules accompanied these changes. Proteins involved in intracellular protein processing / breakdown also proved to be responsive to the LPS-cytokine mix, as were moieties found in the specialized proteasome complex known as the immunoproteasome (29). The proteomic data generated in this study, especially those presented in Tables 1,2, 3 and Supplement Tables 4 and 6 as well as deposited in the public database, provide a broad picture of the protein changes occurring in the human colonic epithelium in response to the pro-inflammatory challenge. This database will be a valuable resource to investigators studying inflammatory diseases of the bowel and, especially, the role of pro-inflammatory cytokines.

Although characterizing the protein changes resulting from LPS-cytokines stimulation are of value in their own right, the primary purposes of this study were to i) assess the capacity of a multimineral intervention to mitigate changes brought about in response to the pro-inflammatory challenge, and ii) determine (conversely) if the pro-inflammatory milieu, itself, would counteract the previously-identified beneficial effects of the multi-mineral intervention on barrier formation. Past studies in mice have demonstrated that Aquamin^®^ reduces high-fat diet-induced colonic inflammation in long-term (15-18 month) studies (47,48) and decreases spontaneous colitis in an IL-10^-/-^ mouse model (49). More recent studies in organoid models have demonstrated the beneficial effects of Aquamin^®^ on the gut barrier structure and function and reduction in pro-inflammatory protein expression (20–22). To summarize, the findings presented here demonstrate that the concomitant presence of Aquamin^®^ did not strongly interfere with the colonic epithelial cell response to the robust pro-inflammatory (LPS-cytokine) stimulus. The profile of protein changes in response to the pro-inflammatory stimulus with or without the multi-mineral supplement (with few exceptions) were very similar to support such a conclusion (Table 1). Comparison of the associated pathways altered in conjunction with the protein changes provides further support for this view (Table 2 and Supplement Tables 4 and 5).

While the overall response to the LPS-cytokine mix was not strongly impacted by Aquamin^®^, the multi-mineral supplement did alter expression of several individual proteins that have been linked to inflammation in the gastrointestinal tract (Table 5 and Supplement Tables 8 and 9). This included down-regulation of proteins with known pro-inflammatory roles and up-regulation of moieties with anti-inflammatory, antioxidant and anti-microbial function. These responses to Aquamin^®^ were observed even in the presence of the LPS-cytokine mix. Whether Aquamin^®^’s effect on expression of any of these individual proteins (alone or in combination) contributes in a meaningful way to a direct anti-inflammatory role is not known at this time and will have to await further work. None-the-less, these findings suggest that in addition to barrier improvement (20-22 and this manuscript), multi-mineral intervention could have other activities that counter inflammation in the gastrointestinal tract.

In parallel with assessing Aquamin^®^’s potential for directly countering pro-inflammatory changes resulting from the LPS-cytokines challenge, we also determined the extent to which the pro-inflammatory mix might counteract the beneficial effects of Aquamin^®^ on barrier formation. This is a critical part of the present work because although we have previously demonstrated multi-mineral improvement in barrier structure/function (20–22), whether barrier improvement would persist in the presence of an acute inflammatory challenge was not addressed. The findings presented here suggest that it would. Proteins that provide for mucosal strength and cohesion (primarily desmosomal proteins and adherens junction proteins) as well as differentiation-related keratins, certain tight junctional elements and contributors to the mucinous layer were up-regulated by Aquamin^®^ in the presence of the LPS-cytokine mix almost as effectively as in its absence. This supports the idea that attempts to improve the colonic barrier will not be wasted simply because any barrier improvement would be undone as a consequence of concomitant inflammation.

To be sure, there are limitations to the present work. Most importantly, although organoid culture technology provides a sophisticated model, no *ex vivo* approach can fully mimic the clinical situation. Acute flare-ups in UC are triggered by multiple factors that originate systemically. It is impossible to control for all these factors in any *in vitro* model. It should be noted, however, that the three cytokines provided here – i.e., TNF-α, IL-1β and IFN-γ – are all known to be present in the inflamed colonic mucosa (43–46). Further, bacterial penetration of the normally impenetrable barrier provides a source of endotoxin (50). Recent gene-expression profiling studies in inflammatory bowel disease have shown that multiple cytokines are, typically, responsible for the pattern of changes seen in most individuals (43). It is not unreasonable to assume, therefore, that while organoid culture is not a perfect model, treatment of colonic organoids with a mix of pro-inflammatory stimuli can produce a milieu that approximates the inflamed human colonic mucosa. Therefore, the use of this technology to investigate potential interventions as carried out here with Aquamin^®^ or previously with other agents (46) would seem to be of value.

Ultimately, the capacity of any intervention to bring about disease mitigation has to be established in controlled clinical studies. As a first step toward this goal, we carried out a 90-day biomarker trial in which thirty healthy adult subjects (10 per arm) were randomized to receive Aquamin^®^ at a level providing 800 mg of calcium per day, calcium alone at the same level or placebo (clinicaltrials.gov: NCT02647671) (23). Before and after intervention, replicate colonic biopsies and fecal specimens were obtained from each subject. Proteomic assessment of tissue biopsies demonstrated up-regulation of many of the same differentiation and barrier-related proteins as were stimulated in organoid culture (20,22). While this biomarker study does not establish clinical efficacy, it demonstrates that protein changes associated with barrier improvement are not limited to organoid culture. In addition to changes in protein signature, subjects in the Aquamin^®^ treatment group also demonstrated reduced levels of several primary and secondary bile acids (51), including some that are thought to be toxic and/or carcinogenic.

As a follow-up to the initial biomarker study, a 180-day trial in subjects with UC in remission (clinicaltrials.gov: NCT03869905) was undertaken. In addition to assessing clinical endpoints in the ongoing trial, biopsies obtained before and after treatment are being examined for changes in barrier protein expression and inflammatory markers just as in the recently completed study. In addition, bowel permeability measurements are being obtained in a separate 90-day study (clinicaltrials.gov: NCT04855799). Results from the ongoing studies will go a long way in establishing the potential for benefit with this approach in UC. Protein changes and permeability measurements will provide insight into therapeutic mechanisms.

While UC is the focus of this work, this disease does not constitute the only clinical situation in which barrier defects are likely to be part of the patho-physiology. Barrier dysfunction has been suggested as a contributing factor in Crohn’s disease as well as UC (52) as well as in irritable bowel syndrome (53) and food allergies / malabsorption syndromes (e.g., celiac disease) (52). Recent studies have also demonstrated disruption of the colonic barrier in the setting of allogenic hematopoietic cell transplantation (54). Barrier dysfunction and poor outcomes are linked. Perhaps most interestingly, high-fat intake as well as obesity, itself, are associated with compromised barrier function (55,56). This provides a link to the systemic inflammatory changes seen in obesity and to the myriad of chronic inflammatory conditions associated with obesity. A low-cost, low-to no-toxicity approach to barrier maintenance could be of value in multiple clinical settings, but this will need to be established in future studies. The present work provides a rationale for these studies.

In summary, past studies have demonstrated that a multi-mineral intervention (Aquamin^®^) has the capacity to up-regulate multiple proteins associated with barrier function in the gastrointestinal tract. The present studies suggest that even in the presence of a strong pro-inflammatory challenge, the beneficial effects on the colonic barrier protein expression are maintained. Further, even though the multi-mineral supplement did not substantially alter the profile of pro-inflammatory proteins up-regulated by the LPS-cytokine mix, several proteins that are part of the inflammatory process were altered. Thus, a direct anti-inflammatory role for Aquamin^®^ appears to exist, independent of the beneficial effects on barrier formation.

## ACKNOWLEDGMENTS

We thank the Proteomics Resource Facility (Pathology Department) for help with proteomic data acquisition; the Microscopy and Imaging Laboratory (MIL) for help with confocal fluorescence microscopy; the Translational Tissue Modeling Laboratory (TTML) at the University of Michigan for help with colon organoid propagation transwell filters seeded with colon organoids. We thank Marigot LTD (Cork, Ireland) for providing Aquamin^®^ as a gift.

## DATA AVAILABILITY STATEMENT

All the data are available in the manuscript text or as Supplemental files. The mass spectrometry proteomics data are available on ProteomeXchange Consortium (PRIDE partner repository) – dataset identifier pending.

## SUPPLEMENTAL MATERIAL

Supplement Table 1. Mineral composition of Aquamin^®^

Supplement Table 2. Demographic characteristics

Supplement Table 3. Common Up-regulated (A) & Down-regulated (B) Proteins

Supplement Table 4. Pathways Up-regulated with LPS-Cytokines

Supplement Table 5. Pathways Up-regulated with Aquamin plus LPS-Cytokines

Supplement Table 6. Pathways Down-regulated with LPS-Cytokines

Supplement Table 7. Pathways Up-regulated with Aquamin

Supplement Table 8. Proteins of interest (Inflammation-related)

Supplement Table 9. Pathways altered by proteins of interest (presented in S. Table 8)

Supplemental File 1. Relevant citations to Inflammation-related proteins presented in the S. Table 8

