## Supplement Table 1 for "Red Algae-Derived Mineral Intervention to Counter Pro-inflammatory Activity in Human Colon Organoids"

**Supplement Table 1. A list of minerals and elements detected in Aquamin® Soluble**

| Element | μg/g | Element | μg/g | Element | μg/g |
|---|---|---|---|---|---|
| Aluminum | 21.6 | Hafnium | 0.038 | Rubidium | 0.031 |
| Antimony | 0.69 | Holmium | 0.010 | Ruthenium | 0.137 |
| Arsenic | 0.239 | Indium | <0.001 | Samarium | 0.037 |
| Barium | 1.76 | Iodine | 1.81 | Scandium | 0.469 |
| Beryllium | <0.5 | Iridium | <0.001 | Selenium | <0.5 |
| Bismuth | <0.5 | Iron | 143 | Silicon | 16.8 |
| Boron | 13.7 | Lanthanum | <0.5 | Silver | <0.5 |
| Cadmium | 0.220 | Lead | 0.084 | Sodium | 2,206 |
| Calcium | 117,000 | Lithium | <0.5 | Strontium | 882 |
| Carbon | 26,600 | Lutetium | <0.001 | Sulfur | 1,241 |
| Cerium | 0.314 | Magnesium | 10,210 | Tantalum | 0.043 |
| Cesium | 0.001 | Manganese | 25.4 | Tellurium | <0.5 |
| Chloride | 612 | Mercury | <0.001 | Terbium | 0.007 |
| Chromium | <0.5 | Molybdenum | <0.5 | Thallium | <0.5 |
| Cobalt | <0.5 | Neodymium | 0.170 | Thorium | 1.30 |
| Copper | <0.5 | Nickel | 0.75 | Thulium | 0.004 |
| Dysprosium | 0.045 | Niobium | <0.5 | Tin | 0.029 |
| Erbium | 0.033 | Osmium | <0.001 | Titanium | 11.4 |
| Europium | 0.013 | Palladium | 0.179 | Tungsten | <0.5 |
| Fluoride | 3.57 | Phosphorous | 189 | Vanadium | <0.5 |
| Gadolinium | 0.044 | Platinum | <0.001 | Ytterbium | 0.030 |
| Gallium | 0.307 | Potassium | 70.0 | Yttrium | <0.5 |
| Germanium | <0.001 | Praseodymium | 0.040 | Zinc | 6.07 |
| Gold | <0.5 | Rhenium | 0.001 | Zirconium | <0.5 |
| | | Rhodium | 0.061 | | |

The mineral composition of Aquamin® Soluble was assessed by Advanced Laboratories, Inc. (Salt Lake City), for client Marigot Limited (Ireland) [Source: 2017 Test Certificate for Aquamin® Soluble]. The levels of individual minerals and trace elements were determined by Inductively Coupled Plasma Optical Emission Spectrometry (ICP-OES) except Carbon (determined by LECO), Chloride, Iodine (determined by Titration), and Fluoride (determined by AOAC 939.11).
