## Supplement Table 2 for "Red Algae-Derived Mineral Intervention to Counter Pro-inflammatory Activity in Human Colon Organoids"

**Supplement Table 2. Demographic characteristics of subjects providing tissue.**

| Sample ID | Age (Years) | Sex | Ethnicity | Biopsy Site |
|---|---|---|---|---|
| Colon-87 | 21 | M | White / Caucasian | Ascending colon |
| Colon-104 | 58 | F | White / Caucasian | Sigmoid colon |
| Colon-105 | 62 | M | White / Caucasian | Sigmoid colon |
| Colon-106 | 50 | M | White / Caucasian | Sigmoid colon |
