## Supplement Table 3 for "Red Algae-Derived Mineral Intervention to Counter Pro-inflammatory Activity in Human Colon Organoids"

**Table 3A. Common up-regulated proteins (at 1.8-fold change)**

| Accession number | Protein name | Gene name |
|---|---|---|
| **7 Common proteins among all groups (LPS-Cytokines, Aquamin and LPS-Cytokines with AQ)** | | |
| Q9NRD8 | Dual oxidase 2 | DUOX2 |
| Q8TAX9 | Gasdermin-B | GSDMB |
| P57764 | Gasdermin-D | GSDMD |
| P17693 | HLA class I histocompatibility antigen, alpha chain G | HLA-G |
| Q30154 | HLA class II histocompatibility antigen, DR beta 5 chain | HLA-DRB5 |
| P14902 | Indoleamine 2,3-dioxygenase 1 | IDO1 |
| Q03169 | Tumor necrosis factor alpha-induced protein 2 | TNFAIP2 |
| **0 Common proteins between LPS-Cytokines and Aquamin groups** | | |
| **24 Common proteins between Aquamin and LPS-Cytokines with AQ groups** | | |
| Q12800 | Alpha-globin transcription factor CP2 | TFCP2 |
| Q9UJ72 | Annexin A10 | ANXA10 |
| O15523 | ATP-dependent RNA helicase DDX3Y | DDX3Y |
| Q12864 | Cadherin-17 | CDH17 |
| P81605 | Dermcidin | DCD |
| Q14126 | Desmoglein-2 | DSG2 |
| Q5D862 | Filaggrin-2 | FLG2 |
| P08263 | Glutathione S-transferase A1 | GSTA1 |
| P19525 | Interferon-induced, double-stranded RNA-activated protein kinase | EIF2AK2 |
| Q92876 | Kallikrein-6 | KLK6 |
| P49862 | Kallikrein-7 | KLK7 |
| P08779 | Keratin, type I cytoskeletal 16 | KRT16 |
| P35527 | Keratin, type I cytoskeletal 9 | KRT9 |
| P04264 | Keratin, type II cytoskeletal 1 | KRT1 |
| P35908 | Keratin, type II cytoskeletal 2 epidermal | KRT2 |
| Q99538 | Legumain | LGMN |
| P08582 | Melanotransferrin | MELTF |
| Q6UX06 | Olfactomedin-4 | OLFM4 |
| Q9Y4D7 | Plexin-D1 | PLXND1 |
| Q9BU68 | Proline-rich protein 15-like protein | PRR15L |
| Q969X1 | Protein lifeguard 3 | TMBIM1 |
| Q08174 | Protocadherin-1 | PCDH1 |
| O00506 | Serine/threonine-protein kinase 25 | STK25 |
| Q03403 | Trefoil factor 2 | TFF2 |
| **68 Common proteins between LPS-Cytokines and LPS-Cytokines with AQ groups** | | |
| P00973 | 2'-5'-oligoadenylate synthase 1 | OAS1 |
| Q9Y6K5 | 2'-5'-oligoadenylate synthase 3 | OAS3 |
| P01009 | Alpha-1-antitrypsin | SERPINA1 |
| Q03518 | Antigen peptide transporter 1 | TAP1 |
| Q03519 | Antigen peptide transporter 2 | TAP2 |
| O95786 | Antiviral innate immune response receptor RIG-I | DDX58 |
| Q9BQE5 | Apolipoprotein L2 | APOL2 |
| P61769 | Beta-2-microglobulin | B2M |
| O00478 | Butyrophilin subfamily 3 member A3 | BTN3A3 |
| P29466 | Caspase-1 | CASP1 |
| P55210 | Caspase-7 | CASP7 |

| | | |
|---|---|---|
| O60911 | Cathepsin L2 | CTSV |
| P27701 | CD82 antigen | CD82 |
| P00450 | Ceruloplasmin | CP |
| Q9NZA1 | Chloride intracellular channel protein 5 | CLIC5 |
| P00751 | Complement factor B | CFB |
| P28838 | Cytosol aminopeptidase | LAP3 |
| Q63HN8 | E3 ubiquitin-protein ligase RNF213 | RNF213 |
| P19474 | E3 ubiquitin-protein ligase TRIM21 | TRIM21 |
| Q9NRD1 | F-box only protein 6 | FBXO6 |
| P13284 | Gamma-interferon-inducible lysosomal thiol reductase | IFI30 |
| Q16666 | Gamma-interferon-inducible protein 16 | IFI16 |
| P32455 | Guanylate-binding protein 1 | GBP1 |
| P32456 | Guanylate-binding protein 2 | GBP2 |
| P04439 | HLA class I histocompatibility antigen, A alpha chain | HLA-A |
| P13747 | HLA class I histocompatibility antigen, alpha chain E | HLA-E |
| P30511 | HLA class I histocompatibility antigen, alpha chain F | HLA-F |
| P01889 | HLA class I histocompatibility antigen, B alpha chain | HLA-B |
| P10321 | HLA class I histocompatibility antigen, C alpha chain | HLA-C |
| P04233 | HLA class II histocompatibility antigen gamma chain | CD74 |
| P28068 | HLA class II histocompatibility antigen, DM beta chain | HLA-DMB |
| P20036 | HLA class II histocompatibility antigen, DP alpha 1 chain | HLA-DPA1 |
| P04440 | HLA class II histocompatibility antigen, DP beta 1 chain | HLA-DPB1 |
| P01903 | HLA class II histocompatibility antigen, DR alpha chain | HLA-DRA |
| P79483 | HLA class II histocompatibility antigen, DR beta 3 chain | HLA-DRB3 |
| P13762 | HLA class II histocompatibility antigen, DR beta 4 chain | HLA-DRB4 |
| P01911 | HLA class II histocompatibility antigen, DRB1 beta chain | HLA-DRB1 |
| P05362 | Intercellular adhesion molecule 1 | ICAM1 |
| P20591 | Interferon-induced GTP-binding protein Mx1 | MX1 |
| Q01629 | Interferon-induced transmembrane protein 2 | IFITM2 |
| P24001 | Interleukin-32 | IL32 |
| P21741 | Midkine | MDK |
| Q9H936 | Mitochondrial glutamate carrier 1 | SLC25A22 |
| P15941 | Mucin-1 | MUC1 |
| P35228 | Nitric oxide synthase, inducible | NOS2 |
| P05120 | Plasminogen activator inhibitor 2 | SERPINB2 |
| Q8IY21 | Probable ATP-dependent RNA helicase DDX60 | DDX60 |
| Q06323 | Proteasome activator complex subunit 1 | PSME1 |
| Q9UL46 | Proteasome activator complex subunit 2 | PSME2 |
| P40306 | Proteasome subunit beta type-10 | PSMB10 |
| P28065 | Proteasome subunit beta type-9 | PSMB9 |
| Q8IXQ6 | Protein mono-ADP-ribosyltransferase PARP9 | PARP9 |
| P21980 | Protein-glutamine gamma-glutamyltransferase 2 | TGM2 |
| P49788 | Retinoic acid receptor responder protein 1 | RARRES1 |
| Q9NUL5 | Shiftless antiviral inhibitor of ribosomal frameshifting protein | SHFL |
| P42224 | Signal transducer and activator of transcription 1-alpha/beta | STAT1 |
| Q8IVG5 | Sterile alpha motif domain-containing protein 9-like | SAMD9L |
| O15533 | Tapasin | TAPBP |
| Q9BX59 | Tapasin-related protein | TAPBPL |
| P19971 | Thymidine phosphorylase | TYMP |
| P23381 | Tryptophan--tRNA ligase, cytoplasmic | WARS |

| | | |
|---|---|---|
| P25942 | Tumor necrosis factor receptor superfamily member 5 | CD40 |
| O14933 | Ubiquitin/ISG15-conjugating enzyme E2 L6 | UBE2L6 |
| P41226 | Ubiquitin-like modifier-activating enzyme 7 | UBA7 |
| P05161 | Ubiquitin-like protein ISG15 | ISG15 |
| Q5EBM0 | UMP-CMP kinase 2, mitochondrial | CMPK2 |
| P62760 | Visinin-like protein 1 | VSNL1 |
| Q9UBW7 | Zinc finger MYM-type protein 2 | ZMYM2 |
| | **11 Unique proteins to LPS-Cytokines** | |
| P00966 | Argininosuccinate synthase | ASS1 |
| Q92851 | Caspase-10 | CASP10 |
| P25774 | Cathepsin S | CTSS |
| Q69YN2 | CWF19-like protein 1 | CWF19L1 |
| P02794 | Ferritin heavy chain | FTH1 |
| Q9Y287 | Integral membrane protein 2B | ITM2B |
| Q13325 | Interferon-induced protein with tetratricopeptide repeats 5 | IFIT5 |
| Q08722 | Leukocyte surface antigen CD47 | CD47 |
| Q9Y5A7 | NEDD8 ultimate buster 1 | NUB1 |
| P14555 | Phospholipase A2, membrane associated | PLA2G2A |
| Q460N5 | Protein mono-ADP-ribosyltransferase PARP14 | PARP14 |
| | **50 Unique proteins to Aquamin** | |
| P15428 | 15-hydroxyprostaglandin dehydrogenase [NAD(+)] | HPGD |
| P52895 | Aldo-keto reductase family 1 member C2 | AKR1C2 |
| Q92625 | Ankyrin repeat and SAM domain-containing protein 1A | ANKS1A |
| Q8NFD5 | AT-rich interactive domain-containing protein 1B | ARID1B |
| Q8NI60 | Atypical kinase COQ8A, mitochondrial, | COQ8A |
| Q14CN2 | Calcium-activated chloride channel regulator 4 | CLCA4 |
| P00915 | Carbonic anhydrase 1 | CA1 |
| P06731 | Carcinoembryonic antigen-related cell adhesion molecule 5 | CEACAM5 |
| P40879 | Chloride anion exchanger (Protein DRA) | SLC26A3 |
| Q8NCR9 | Clarin-3 | CLRN3 |
| P99999 | Cytochrome c | CYCS |
| P00395 | Cytochrome c oxidase subunit 1 | COX1 |
| P30046 | D-dopachrome decarboxylase | DDT |
| Q9NZC4 | ETS homologous factor, hEHF | EHF |
| P41214 | Eukaryotic translation initiation factor 2D | EIF2D |
| O94887 | FERM, ARHGEF and pleckstrin domain-containing protein 2 | FARP2 |
| P85037 | Forkhead box protein K1 | FOXK1 |
| Q7L5D6 | Golgi to ER traffic protein 4 homolog | GET4 |
| Q14623 | Indian hedgehog protein | IHH |
| Q8TB37 | Iron-sulfur protein NUBPL | NUBPL |
| P13645 | Keratin, type I cytoskeletal 10 | KRT10 |
| P02533 | Keratin, type I cytoskeletal 14 | KRT14 |
| Q7Z794 | Keratin, type II cytoskeletal 1b | KRT77 |
| P13647 | Keratin, type II cytoskeletal 5 | KRT5 |
| Q6KB66 | Keratin, type II cytoskeletal 80 | KRT80 |
| Q9Y250 | Leucine zipper putative tumor suppressor 1 | LZTS1 |
| Q6UX82 | Ly6/PLAUR domain-containing protein 8 | LYPD8 |
| Q96CM8 | Medium-chain acyl-CoA ligase ACSF2 | ACSF2 |
| Q15555 | Microtubule-associated protein RP/EB family member 2 | MAPRE2 |
| Q9Y3Q0 | N-acetylated-alpha-linked acidic dipeptidase 2 | NAALAD2 |

| | | |
|---|---|---|
| O95497 | Pantetheinase | VNN1 |
| P30039 | Phenazine biosynthesis-like domain-containing protein | PBLD |
| P35558 | Phosphoenolpyruvate carboxykinase, cytosolic [GTP] | PCK1 |
| Q6VY07 | Phosphofurin acidic cluster sorting protein 1 | PACS1 |
| O00592 | Podocalyxin | PODXL |
| O43653 | Prostate stem cell antigen | PSCA |
| Q9BVL4 | Protein adenylyltransferase SelO, mitochondrial, | SELENOO |
| Q6P996 | Pyridoxal-dependent decarboxylase domain-containing protein 1 | PDXDC1 |
| Q12913 | Receptor-type tyrosine-protein phosphatase eta | PTPRJ |
| Q9NX52 | Rhomboid-related protein 2 | RHBDL2 |
| Q9UEW8 | STE20/SPS1-related proline-alanine-rich protein kinase | STK39 |
| P0DMN0 | Sulfotransferase 1A4 | SULT1A4 |
| O43704 | Sulfotransferase 1B1 | SULT1B1 |
| O00142 | Thymidine kinase 2, mitochondrial | TK2 |
| P30408 | Transmembrane 4 L6 family member 1 | TM4SF1 |
| Q9BZV1 | UBX domain-containing protein 6 | UBXN6 |
| O15294 | UDP-N-acetylglucosamine--peptide N-acetylglucosaminyltransferase 110 kDa subunit | OGT |
| O14975 | Very long-chain acyl-CoA synthetase | SLC27A2 |
| P12955 | Xaa-Pro dipeptidase | PEPD |
| Q14966 | Zinc finger protein 638 | ZNF638 |
| | **35 Unique proteins to Aquamin with LPS-Cytokines** | |
| Q92604 | Acyl-CoA:lysophosphatidylglycerol acyltransferase 1 | LPGAT1 |
| P00352 | Aldehyde dehydrogenase 1A1 | ALDH1A1 |
| P15144 | Aminopeptidase N | ANPEP |
| P08174 | Complement decay-accelerating factor | CD55 |
| P08684 | Cytochrome P450 3A4 | CYP3A4 |
| O75891 | Cytosolic 10-formyltetrahydrofolate dehydrogenase | ALDH1L1 |
| O75592 | E3 ubiquitin-protein ligase MYCBP2 | MYCBP2 |
| P05062 | Fructose-bisphosphate aldolase B | ALDOB |
| A2VDF0 | Fucose mutarotase | FUOM |
| Q2TB90 | Hexokinase HKDC1 | HKDC1 |
| Q86YZ3 | Hornerin | HRNR |
| P02538 | Keratin, type II cytoskeletal 6A | KRT6A |
| P04732 | Metallothionein-1E | MT1E |
| P80294 | Metallothionein-1H | MT1H |
| P51608 | Methyl-CpG-binding protein 2 | MECP2 |
| Q685J3 | Mucin-17 | MUC17 |
| Q96HD9 | N-acyl-aromatic-L-amino acid amidohydrolase | ACY3 |
| Q9NTG7 | NAD-dependent protein deacetylase sirtuin-3, mitochondrial | SIRT3 |
| P49281 | Natural resistance-associated macrophage protein 2 | SLC11A2 |
| Q9NZT2 | Opioid growth factor receptor | OGFR |
| A1L390 | Pleckstrin homology domain-containing family G member 3 | PLEKHG3 |
| P29590 | Promyelocytic leukemia protein | PML |
| P31151 | Protein S100-A7 | S100A7 |
| Q5T4F4 | Protrudin | ZFYVE27 |
| P07998 | Ribonuclease pancreatic | RNASE1 |
| Q9NW13 | RNA-binding protein 28 | RBM28 |
| Q14141 | Septin-6 | SEPT6 |
| P50453 | Serpin B9 (Peptidase inhibitor 9) | SERPINB9 |

| | | |
|---|---|---|
| Q8TCT8 | Signal peptide peptidase-like 2A | SPPL2A |
| P48764 | Sodium/hydrogen exchanger 3 | SLC9A3 |
| Q6UX04 | Spliceosome-associated protein CWC27 homolog | CWC27 |
| P13726 | Tissue factor (Thromboplastin) | F3 |
| O14545 | TRAF-type zinc finger domain-containing protein 1 | TRAFD1 |
| P09758 | Tumor-associated calcium signal transducer 2 | TACSTD2 |
| O75795 | UDP-glucuronosyltransferase 2B17 | UGT2B17 |

These data also presented in Figure 3 Venn plots.

**Table 3B. Common down-regulated proteins (at 1.8-fold change)**

| Accession number | Protein name | Gene name |
|---|---|---|
| **0 Common proteins among all groups (LPS-Cytokines, Aquamin and LPS-Cytokines with AQ)** | | |
| **0 Common proteins between LPS-Cytokines and Aquamin groups** | | |
| **2 Common proteins between Aquamin and LPS-Cytokines with AQ groups** | | |
| P02675 | Fibrinogen beta chain | FGB |
| P09486 | SPARC | SPARC |
| **3 Common proteins between LPS-Cytokines and LPS-Cytokines with AQ groups** | | |
| P19957 | Elafin | PI3 |
| P28072 | Proteasome subunit beta type-6 | PSMB6 |
| Q99436 | Proteasome subunit beta type-7 | PSMB7 |
| **5 Unique proteins to LPS-Cytokines** | | |
| P03973 | Antileukoproteinase | SLPI |
| P08263 | Glutathione S-transferase A1 | GSTA1 |
| P35558 | Phosphoenolpyruvate carboxykinase, cytosolic [GTP] | PCK1 |
| P28074 | Proteasome subunit beta type-5 | PSMB5 |
| O15492 | Regulator of G-protein signaling 16 | RGS16 |
| **8 Unique proteins to Aquamin** | | |
| P02656 | Apolipoprotein C-III | APOC3 |
| P62633 | CCHC-type zinc finger nucleic acid binding protein | CNBP |
| P02679 | Fibrinogen gamma chain | FGG |
| P80294 | Metallothionein-1H | MT1H |
| P05204 | Non-histone chromosomal protein HMG-17 | HMGN2 |
| Q5TBB1 | Ribonuclease H2 subunit B | RNASEH2B |
| Q9Y2S6 | Translation machinery-associated protein 7 | TMA7 |
| Q6RW13 | Type-1 angiotensin II receptor-associated protein | AGTRAP |
| **6 Unique proteins to Aquamin with LPS-Cytokines** | | |
| P30711 | Glutathione S-transferase theta-1 | GSTT1 |
| P47902 | Homeobox protein CDX-1 | CDX1 |
| P10586 | Receptor-type tyrosine-protein phosphatase F | PTPRF |
| Q9BYZ8 | Regenerating islet-derived protein 4 | REG4 |
| Q9HCB6 | Spondin-1 | SPON1 |
| Q96IQ7 | V-set and immunoglobulin domain-containing protein 2 | VSIG2 |

These data also presented in Figure 3 Venn plots.
