## Supplement Table 4 for "Red Algae-Derived Mineral Intervention to Counter Pro-inflammatory Activity in Human Colon Organoids"

**Supplement Table 4. Pathways associated with up-regulated proteins altered with LPS-Cytokines.**

| Pathways | Entities pValue | Mapped entities |
|---|---|---|
| Interferon Signaling | $1.11x10^{-16}$ | HLA-E;HLA-A;TRIM21;OAS3;HLA-G;HLA-DRB4;STAT1;HLA-DRB1;HLA-DRB3;HLA-DPB1; B2M;HLA-DPA1;IFITM2;HLA-F;DDX58;IFIT5;MX1;GBP1;GBP2;HLA-C;IFI30;HLA-DRA;ISG15; HLA-B;ICAM1;OAS1;UBE2L6;UBA7;HLA-DRB5 |
| Interferon gamma signaling | $1.11x10^{-16}$ | HLA-F;HLA-E;HLA-A;TRIM21;OAS3;HLA-G;GBP1;GBP2;HLA-DRB4;HLA-C;STAT1;IFI30; HLA-DRA;HLA-DRB1;HLA-B;HLA-DRB3;ICAM1;OAS1;HLA-DPB1;B2M;HLA-DPA1;HLA-DRB5 |
| Immune System | $1.11x10^{-16}$ | IL32;IFI16;HLA-A;HLA-E;CD47;OAS3;GSDMD;PLA2G2A;STAT1;HLA-DRB1;HLA-DRB3;CASP10; HLA-DPB1;HLA-DPA1;HLA-F;IFIT5;CASP1;GBP1;RNF213;GBP2;BTN3A3;HLA-C;IFI30;HLA-DRA; FBXO6;OAS1;CTSS;FTH1;HLA-DMB;PSMB9;TRIM21;HLA-G;PSME1;MUC1;HLA-DRB4;TAPBP; CD40;PSMB10;B2M;IFITM2;DDX58;PSME2;MX1;NOS2;CTSV;ISG15;SERPINA1;HLA-B; SERPINB2;ICAM1;TAP1;CFB;TAP2;UBA7;CD74;UBE2L6;HLA-DRB5 |
| Cytokine Signaling in Immune system | $1.11x10^{-16}$ | IL32;HLA-E;HLA-A;PSMB9;TRIM21;OAS3;HLA-G;GSDMD;PSME1;MUC1;HLA-DRB4;STAT1; HLA-DRB1;HLA-DRB3;CD40;PSMB10;HLA-DPB1;B2M;HLA-DPA1;IFITM2;HLA-F;DDX58; IFIT5;PSME2;CASP1;MX1;GBP1;NOS2;GBP2;HLA-C;IFI30;HLA-DRA;ISG15;HLA-B; SERPINB2;ICAM1;OAS1;UBE2L6;UBA7;HLA-DRB5 |
| Adaptive Immune System | $6.44x10^{-15}$ | HLA-DMB;HLA-E;HLA-A;PSMB9;TRIM21;HLA-G;PSME1;HLA-DRB4;HLA-DRB1;HLA-DRB3; TAPBP;CD40;PSMB10;HLA-DPB1;B2M;HLA-DPA1;HLA-F;PSME2;RNF213;CTSV;HLA-C; BTN3A3;IFI30;HLA-DRA;HLA-B;FBXO6;ICAM1;CTSS;TAP1;TAP2;CD74;UBE2L6;UBA7;HLA-DRB5 |
| Interferon alpha/beta signaling | $1.90x10^{-14}$ | HLA-F;IFIT5;MX1;HLA-E;HLA-A;HLA-G;OAS3;HLA-C;GBP2;STAT1;ISG15;HLA-B;OAS1;IFITM2 |
| Antigen processing-Cross presentation | $4.80x10^{-13}$ | HLA-F;PSME2;HLA-E;HLA-A;PSMB9;HLA-G;PSME1;CTSV;HLA-C;HLA-B;TAPBP; PSMB10;B2M;TAP1;CTSS;TAP2 |
| ER-Phagosome pathway | $2.57x10^{-11}$ | HLA-F;PSME2;HLA-E;HLA-A;PSMB9;HLA-G;PSME1;HLA-C;HLA-B;TAPBP; PSMB10;B2M;TAP1;TAP2 |
| Class I MHC mediated antigen processing & presentation | $2.58x10^{-11}$ | HLA-F;PSME2;HLA-E;HLA-A;PSMB9;TRIM21;HLA-G;PSME1;RNF213;CTSV;HLA-C;HLA-B; TAPBP;FBXO6;PSMB10;CTSS;TAP1;B2M;TAP2;UBE2L6;UBA7 |
| MHC class II antigen presentation | $1.11x10^{-9}$ | HLA-DMB;IFI30;HLA-DRA;HLA-DRB1;HLA-DRB3;HLA-DPB1;CTSS;CD74;CTSV;HLA-DPA1; HLA-DRB4;HLA-DRB5 |
| Downstream TCR signaling | $1.76x10^{-9}$ | PSME2;HLA-DRA;PSMB9;HLA-DRB1;HLA-DRB3;PSMB10;PSME1;HLA-DPB1;HLA-DPA1; HLA-DRB4;HLA-DRB5 |
| Antigen Presentation: Folding, assembly and peptide loading of class I MHC | $5.08x10^{-9}$ | HLA-F;HLA-A;HLA-E;HLA-B;TAPBP;HLA-G;B2M;TAP1;TAP2;HLA-C |
| TCR signaling | $1.04x10^{-8}$ | PSME2;HLA-DRA;PSMB9;HLA-DRB1;HLA-DRB3;PSMB10;PSME1;HLA-DPB1;HLA-DPA1; HLA-DRB4;HLA-DRB5 |
| Endosomal/Vacuolar pathway | $1.21x10^{-8}$ | HLA-F;HLA-A;HLA-E;HLA-B;HLA-G;B2M;CTSS;CTSV;HLA-C |
| Translocation of ZAP-70 to Immunological synapse | $2.19x10^{-8}$ | HLA-DRA;HLA-DRB1;HLA-DRB3;HLA-DPB1;HLA-DPA1;HLA-DRB5;HLA-DRB4 |
| Phosphorylation of CD3 and TCR zeta chains | $3.66x10^{-8}$ | HLA-DRA;HLA-DRB1;HLA-DRB3;HLA-DPB1;HLA-DPA1;HLA-DRB5;HLA-DRB4 |
| PD-1 signaling | $4.30x10^{-8}$ | HLA-DRA;HLA-DRB1;HLA-DRB3;HLA-DPB1;HLA-DPA1;HLA-DRB5;HLA-DRB4 |
| SARS-CoV-2 activates/modulates innate and adaptive immune responses | $8.40x10^{-8}$ | HLA-F;DDX58;STAT1;HLA-A;HLA-E;ISG15;HLA-B;HLA-G;B2M;HLA-C |
| Generation of second messenger molecules | $2.50x10^{-7}$ | HLA-DRA;HLA-DRB1;HLA-DRB3;HLA-DPB1;HLA-DPA1;HLA-DRB5;HLA-DRB4 |
| Antiviral mechanism by IFN-stimulated genes | $4.84x10^{-7}$ | DDX58;STAT1;MX1;ISG15;OAS3;OAS1;UBA7;UBE2L6 |
| Innate Immune System | $6.93x10^{-7}$ | IFI16;HLA-E;PSMB9;TRIM21;CD47;GSDMD;PSME1;PLA2G2A;MUC1;CASP10;PSMB10;B2M; DDX58;PSME2;CASP1;NOS2;CTSV;HLA-C;ISG15;SERPINA1;HLA-B;CTSS; CFB;UBE2L6;UBA7;FTH1 |
| SARS-CoV-2 Infection | $9.81x10^{-7}$ | HLA-F;DDX58;STAT1;HLA-E;HLA-A;ISG15;HLA-B;PARP14;HLA-G;PARP9;B2M;HLA-C |
| SARS-CoV-2-host interactions | $2.81x10^{-6}$ | HLA-F;DDX58;STAT1;HLA-A;HLA-E;ISG15;HLA-B;HLA-G;B2M;HLA-C |

| | | |
|---|---|---|
| Costimulation by the CD28 family | $6.64 \times 10^{-6}$ | HLA-DRA;HLA-DRB1;HLA-DRB3;HLA-DPB1;HLA-DPA1;HLA-DRB5;HLA-DRB4 |
| ISG15 antiviral mechanism | $3.81 \times 10^{-5}$ | DDX58;STAT1;MX1;ISG15;UBA7;UBE2L6 |
| SARS-CoV Infections | $5.75 \times 10^{-5}$ | HLA-F;DDX58;STAT1;HLA-E;HLA-A;ISG15;HLA-B;PARP14;HLA-G;PARP9;B2M;HLA-C |
| Signaling by Interleukins | $6.13 \times 10^{-5}$ | STAT1;PSME2;CASP1;IL32;PSMB9;SERPINB2;ICAM1;PSMB10;GSDMD;PSME1;MUC1;NOS2 |
| OAS antiviral response | $6.86 \times 10^{-5}$ | DDX58;OAS3;OAS1 |
| Negative regulators of DDX58/IFIH1 signaling | $1.64 \times 10^{-4}$ | DDX58;ISG15;UBE2L6;UBA7 |
| Regulation of RUNX2 expression and activity | $2.80 \times 10^{-4}$ | STAT1;PSME2;PSMB9;PSMB10;PSME1 |
| Immunoregulatory interactions between a Lymphoid and a non-Lymphoid cell | $4.71 \times 10^{-4}$ | HLA-F;HLA-A;HLA-E;HLA-B;CD40;ICAM1;HLA-G;B2M;HLA-C |
| DDX58/IFIH1-mediated induction of interferon-alpha/beta | $5.24 \times 10^{-4}$ | DDX58;ISG15;CASP10;UBA7;UBE2L6 |
| Antigen processing: Ubiquitination & Proteasome degradation | $6.24 \times 10^{-4}$ | PSME2;PSMB9;FBXO6;TRIM21;PSMB10;PSME1;RNF213;UBA7;UBE2L6 |
| Regulation of activated PAK-2p34 by proteasome mediated degradation | $6.27 \times 10^{-4}$ | PSME2;PSMB9;PSMB10;PSME1 |
| Cross-presentation of soluble exogenous antigens (endosomes) | $6.27 \times 10^{-4}$ | PSME2;PSMB9;PSMB10;PSME1 |
| Regulation of ornithine decarboxylase (ODC) | $6.75 \times 10^{-4}$ | PSME2;PSMB9;PSMB10;PSME1 |
| p53-Independent G1/S DNA damage checkpoint | $7.25 \times 10^{-4}$ | PSME2;PSMB9;PSMB10;PSME1 |
| Ubiquitin Mediated Degradation of Phosphorylated Cdc25A | $7.25 \times 10^{-4}$ | PSME2;PSMB9;PSMB10;PSME1 |
| p53-Independent DNA Damage Response | $7.25 \times 10^{-4}$ | PSME2;PSMB9;PSMB10;PSME1 |
| GSK3B and BTRC:CUL1-mediated-degradation of NFE2L2 | $7.25 \times 10^{-4}$ | PSME2;PSMB9;PSMB10;PSME1 |
| Ubiquitin-dependent degradation of Cyclin D | $7.25 \times 10^{-4}$ | PSME2;PSMB9;PSMB10;PSME1 |
| Autodegradation of the E3 ubiquitin ligase COP1 | $7.25 \times 10^{-4}$ | PSME2;PSMB9;PSMB10;PSME1 |
| Infectious disease | $7.62 \times 10^{-4}$ | HLA-F;DDX58;PSME2;CASP1;HLA-E;HLA-A;PSMB9;PARP14;HLA-G;GSDMD; PSME1;PARP9;NOS2;RNF213;HLA-C;STAT1;ISG15;HLA-B;PSMB10;B2M |
| Vpu mediated degradation of CD4 | $7.78 \times 10^{-4}$ | PSME2;PSMB9;PSMB10;PSME1 |
| Regulation of Apoptosis | $7.78 \times 10^{-4}$ | PSME2;PSMB9;PSMB10;PSME1 |
| FBXL7 down-regulates AURKA during mitotic entry and in early mitosis | $8.92 \times 10^{-4}$ | PSME2;PSMB9;PSMB10;PSME1 |
| SCF-beta-TrCP mediated degradation of Emi1 | $8.92 \times 10^{-4}$ | PSME2;PSMB9;PSMB10;PSME1 |
| Degradation of AXIN | $8.92 \times 10^{-4}$ | PSME2;PSMB9;PSMB10;PSME1 |
| Negative regulation of NOTCH4 signaling | $8.92 \times 10^{-4}$ | PSME2;PSMB9;PSMB10;PSME1 |
| Regulation of RUNX3 expression and activity | $8.92 \times 10^{-4}$ | PSME2;PSMB9;PSMB10;PSME1 |
| AUF1 (hnRNP D0) binds and destabilizes mRNA | $9.53 \times 10^{-4}$ | PSME2;PSMB9;PSMB10;PSME1 |

| | | |
|---|---|---|
| Vif-mediated degradation of APOBEC3G | $9.53x10^{-4}$ | PSME2;PSMB9;PSMB10;PSME1 |
| Hh mutants are degraded by ERAD | $9.53x10^{-4}$ | PSME2;PSMB9;PSMB10;PSME1 |
| Degradation of DVL | 0.001 | PSME2;PSMB9;PSMB10;PSME1 |
| Stabilization of p53 | 0.001 | PSME2;PSMB9;PSMB10;PSME1 |
| Hh mutants abrogate ligand secretion | 0.001 | PSME2;PSMB9;PSMB10;PSME1 |
| NIK-->noncanonical NF-kB signaling | 0.001 | PSME2;PSMB9;PSMB10;PSME1 |
| Metabolism of polyamines | 0.001 | PSME2;PSMB9;PSMB10;PSME1 |
| TNFR2 non-canonical NF-kB pathway | 0.001 | PSME2;PSMB9;CD40;PSMB10;PSME1 |
| Degradation of GLI1 by the proteasome | 0.001 | PSME2;PSMB9;PSMB10;PSME1 |
| Degradation of GLI2 by the proteasome | 0.001 | PSME2;PSMB9;PSMB10;PSME1 |
| GLI3 is processed to GLI3R by the proteasome | 0.001 | PSME2;PSMB9;PSMB10;PSME1 |
| SCF(Skp2)-mediated degradation of p27/p21 | 0.001 | PSME2;PSMB9;PSMB10;PSME1 |
| Dectin-1 mediated noncanonical NF-kB signaling | 0.001 | PSME2;PSMB9;PSMB10;PSME1 |
| KEAP1-NFE2L2 pathway | 0.001 | PSME2;PSMB9;TRIM21;PSMB10;PSME1 |
| Interleukin-1 family signaling | 0.001 | PSME2;CASP1;PSMB9;PSMB10;GSDMD;PSME1 |
| Defective CFTR causes cystic fibrosis | 0.001 | PSME2;PSMB9;PSMB10;PSME1 |
| Programmed Cell Death | 0.001 | PSME2;CASP1;CASP7;PSMB9;PSMB10;GSDMD;PSME1 |
| Autodegradation of Cdh1 by Cdh1:APC/C | 0.002 | PSME2;PSMB9;PSMB10;PSME1 |
| Asymmetric localization of PCP proteins | 0.002 | PSME2;PSMB9;PSMB10;PSME1 |
| Hedgehog ligand biogenesis | 0.002 | PSME2;PSMB9;PSMB10;PSME1 |
| Oxygen-dependent proline hydroxylation of Hypoxia-inducible Factor Alpha | 0.002 | PSME2;PSMB9;PSMB10;PSME1 |
| p53-Dependent G1 DNA Damage Response | 0.002 | PSME2;PSMB9;PSMB10;PSME1 |
| p53-Dependent G1/S DNA damage checkpoint | 0.002 | PSME2;PSMB9;PSMB10;PSME1 |
| Activation of NF-kappaB in B cells | 0.002 | PSME2;PSMB9;PSMB10;PSME1 |
| APC/C:Cdc20 mediated degradation of Securin | 0.002 | PSME2;PSMB9;PSMB10;PSME1 |
| G1/S DNA Damage Checkpoints | 0.002 | PSME2;PSMB9;PSMB10;PSME1 |
| Termination of translesion DNA synthesis | 0.002 | ISG15;UBE2L6;UBA7 |
| Regulation of RAS by GAPs | 0.002 | PSME2;PSMB9;PSMB10;PSME1 |
| Regulation of PTEN stability and activity | 0.002 | PSME2;PSMB9;PSMB10;PSME1 |
| Interleukin-1 processing | 0.002 | CASP1;GSDMD |
| Orc1 removal from chromatin | 0.002 | PSME2;PSMB9;PSMB10;PSME1 |
| Transcriptional regulation by RUNX2 | 0.002 | STAT1;PSME2;PSMB9;PSMB10;PSME1 |
| Cdc20:Phospho-APC/C mediated degradation of Cyclin A | 0.002 | PSME2;PSMB9;PSMB10;PSME1 |
| CDK-mediated phosphorylation and removal of Cdc6 | 0.002 | PSME2;PSMB9;PSMB10;PSME1 |

| | | |
|---|---|---|
| APC/C:Cdh1 mediated degradation of Cdc20 and other APC/C:Cdh1 targeted proteins in late mitosis/early G1 | 0.003 | PSME2;PSMB9;PSMB10;PSME1 |
| APC:Cdc20 mediated degradation of cell cycle proteins prior to satisfation of the cell cycle checkpoint | 0.003 | PSME2;PSMB9;PSMB10;PSME1 |
| Neddylation | 0.003 | PSME2;PSMB9;FBXO6;PSMB10;PSME1;NUB1 |
| Apoptosis | 0.003 | PSME2;CASP7;PSMB9;PSMB10;GSDMD;PSME1 |
| APC/C:Cdc20 mediated degradation of mitotic proteins | 0.003 | PSME2;PSMB9;PSMB10;PSME1 |
| Cellular response to hypoxia | 0.003 | PSME2;PSMB9;PSMB10;PSME1 |
| Activation of APC/C and APC/C:Cdc20 mediated degradation of mitotic proteins | 0.003 | PSME2;PSMB9;PSMB10;PSME1 |
| ABC transporter disorders | 0.003 | PSME2;PSMB9;PSMB10;PSME1 |
| The role of GTSE1 in G2/M progression after G2 checkpoint | 0.003 | PSME2;PSMB9;PSMB10;PSME1 |
| Nuclear events mediated by NFE2L2 | 0.003 | PSME2;PSMB9;PSMB10;PSME1 |
| Translesion synthesis by Y family DNA polymerases bypasses lesions on DNA template | 0.003 | ISG15;UBE2L6;UBA7 |
| Regulation of APC/C activators between G1/S and early anaphase | 0.004 | PSME2;PSMB9;PSMB10;PSME1 |
| Downstream signaling events of B Cell Receptor (BCR) | 0.004 | PSME2;PSMB9;PSMB10;PSME1 |
| Cyclin E associated events during G1/S transition | 0.004 | PSME2;PSMB9;PSMB10;PSME1 |
| Signaling by NOTCH4 | 0.004 | PSME2;PSMB9;PSMB10;PSME1 |
| Degradation of beta-catenin by the destruction complex | 0.004 | PSME2;PSMB9;PSMB10;PSME1 |
| TP53 Regulates Transcription of Caspase Activators and Caspases | 0.004 | CASP1;CASP10 |
| Maturation of nucleoprotein | 0.004 | PARP14;PARP9 |
| Hedgehog 'on' state | 0.004 | PSME2;PSMB9;PSMB10;PSME1 |
| Cyclin A:Cdk2-associated events at S phase entry | 0.004 | PSME2;PSMB9;PSMB10;PSME1 |
| NF-kB activation through FADD/RIP-1 pathway mediated by caspase-8 and -10 | 0.005 | DDX58;CASP10 |
| Trafficking and processing of endosomal TLR | 0.005 | CTSS;CTSV |
| Regulation of mitotic cell cycle | 0.005 | PSME2;PSMB9;PSMB10;PSME1 |
| APC/C-mediated degradation of cell cycle proteins | 0.005 | PSME2;PSMB9;PSMB10;PSME1 |
| Regulation of mRNA stability by proteins that bind AU-rich elements | 0.005 | PSME2;PSMB9;PSMB10;PSME1 |
| MAPK6/MAPK4 signaling | 0.005 | PSME2;PSMB9;PSMB10;PSME1 |

| | | |
|---|---|---|
| Host Interactions of HIV factors | 0.005 | PSME2;PSMB9;PSMB10;PSME1;B2M |
| Signaling by NOTCH | 0.005 | STAT1;PSME2;PSMB9;PSMB10;PSME1;MDK |
| Switching of origins to a post-replicative state | 0.006 | PSME2;PSMB9;PSMB10;PSME1 |
| PCP/CE pathway | 0.006 | PSME2;PSMB9;PSMB10;PSME1 |
| Interleukin-6 signaling | 0.006 | STAT1 |
| DNA Damage Bypass | 0.006 | ISG15;UBE2L6;UBA7 |
| UCH proteinases | 0.007 | PSME2;PSMB9;PSMB10;PSME1 |
| Transcriptional regulation by RUNX3 | 0.007 | PSME2;PSMB9;PSMB10;PSME1 |
| STING mediated induction of host immune responses | 0.007 | IFI16;TRIM21 |
| Maturation of nucleoprotein | 0.007 | PARP14;PARP9 |
| CLEC7A (Dectin-1) signaling | 0.007 | PSME2;PSMB9;PSMB10;PSME1 |
| RUNX1 regulates transcription of genes involved in differentiation of HSCs | 0.007 | PSME2;PSMB9;PSMB10;PSME1 |
| ABC-family proteins mediated transport | 0.008 | PSME2;PSMB9;PSMB10;PSME1 |
| Signaling by cytosolic FGFR1 fusion mutants | 0.009 | STAT1;ZMYM2 |
| Interleukin-12 family signaling | 0.009 | STAT1;PSME2;SERPINB2 |
| Nicotinamide salvaging | 0.010 | PARP14;PARP9 |
| Apoptotic factor-mediated response | 0.011 | CASP7;GSDMD |
| Interleukin-4 and Interleukin-13 signaling | 0.011 | STAT1;ICAM1;NOS2;MUC1 |
| Assembly of the pre-replicative complex | 0.011 | PSME2;PSMB9;PSMB10;PSME1 |
| C-type lectin receptors (CLRs) | 0.011 | PSME2;PSMB9;PSMB10;PSME1;MUC1 |
| Hedgehog 'off' state | 0.012 | PSME2;PSMB9;PSMB10;PSME1 |
| Interleukin-1 signaling | 0.012 | PSME2;PSMB9;PSMB10;PSME1 |
| Signaling by phosphorylated juxtamembrane, extracellular and kinase domain KIT mutants | 0.013 | STAT1 |
| Signaling by KIT in disease | 0.013 | STAT1 |
| Disorders of transmembrane transporters | 0.013 | CP;PSME2;PSMB9;PSMB10;PSME1 |
| Synthesis of DNA | 0.014 | PSME2;PSMB9;PSMB10;PSME1 |
| Defective SLC40A1 causes hemochromatosis 4 (HFE4) (macrophages) | 0.015 | CP |
| Defective CP causes aceruloplasminemia (ACERULOP) | 0.015 | CP |
| Regulation of IFNA/IFNB signaling | 0.017 | STAT1 |
| DNA Replication Pre-Initiation | 0.018 | PSME2;PSMB9;PSMB10;PSME1 |
| Cellular response to chemical stress | 0.018 | PSME2;PSMB9;TRIM21;PSMB10;PSME1 |
| Pyroptosis | 0.019 | CASP1;GSDMD |
| Purinergic signaling in leishmaniasis infection | 0.019 | CASP1;GSDMD |
| Cell recruitment (pro-inflammatory response) | 0.019 | CASP1;GSDMD |
| G1/S Transition | 0.019 | PSME2;PSMB9;PSMB10;PSME1 |

| | | |
|---|---|---|
| Interleukin-6 family signaling | 0.020 | STAT1 |
| Ub-specific processing proteases | 0.021 | DDX58;PSME2;PSMB9;PSMB10;PSME1 |
| Transcriptional regulation by RUNX1 | 0.022 | PSME2;PSMB9;PSMB10;PSME1;CTSV |
| PTEN Regulation | 0.022 | PSME2;PSMB9;PSMB10;PSME1 |
| DAP12 signaling | 0.023 | HLA-E;B2M |
| The AIM2 inflammasome | 0.023 | CASP1 |
| Nicotinate metabolism | 0.024 | PARP14;PARP9 |
| Diseases of signal transduction by growth factor receptors and second messengers | 0.024 | STAT1;PSME2;ZMYM2;PSMB9;PSMB10;PSME1;RNF213 |
| Amyloid fiber formation | 0.025 | ITM2B;B2M;UBE2L6 |
| Beta-catenin independent WNT signaling | 0.025 | PSME2;PSMB9;PSMB10;PSME1 |
| Metabolism of amino acids and derivatives | 0.025 | DUOX2;PSME2;IDO1;PSMB9;ASS1;PSMB10;PSME1 |
| Infection with Mycobacterium tuberculosis | 0.026 | B2M;NOS2;RNF213 |
| Disease | 0.026 | HLA-F;DDX58;CP;PSME2;CASP1;HLA-E;HLA-A;PSMB9;PARP14;HLA-G;GSDMD;PARP9; PSME1;NOS2;MUC1;RNF213;HLA-C;STAT1;ZMYM2;ISG15;HLA-B;PSMB10;B2M |
| FGFR1 mutant receptor activation | 0.027 | STAT1;ZMYM2 |
| Mitotic G1 phase and G1/S transition | 0.028 | PSME2;PSMB9;PSMB10;PSME1 |
| Translation of Structural Proteins | 0.028 | PARP14;PARP9 |
| Signaling by Hedgehog | 0.029 | PSME2;PSMB9;PSMB10;PSME1 |
| G2/M Checkpoints | 0.029 | PSME2;PSMB9;PSMB10;PSME1 |
| Inhibition of nitric oxide production | 0.030 | NOS2 |
| Neutrophil degranulation | 0.031 | SERPINA1;HLA-B;CD47;GSDMD;B2M;CTSS;FTH1;HLA-C |
| DNA Replication | 0.033 | PSME2;PSMB9;PSMB10;PSME1 |
| Gene and protein expression by JAK-STAT signaling after Interleukin-12 stimulation | 0.033 | PSME2;SERPINB2 |
| Response of Mtb to phagocytosis | 0.036 | NOS2;RNF213 |
| S Phase | 0.037 | PSME2;PSMB9;PSMB10;PSME1 |
| Alternative complement activation | 0.038 | CFB |
| The IPAF inflammasome | 0.038 | CASP1 |
| FasL/ CD95L signaling | 0.038 | CASP10 |
| Signaling by FGFR1 in disease | 0.040 | STAT1;ZMYM2 |
| FCERI mediated NF-kB activation | 0.040 | PSME2;PSMB9;PSMB10;PSME1 |
| HIV Infection | 0.040 | PSME2;PSMB9;PSMB10;PSME1;B2M |
| Regulation of expression of SLITs and ROBOs | 0.044 | PSME2;PSMB9;PSMB10;PSME1 |
| Activation of caspases through apoptosome-mediated cleavage | 0.045 | CASP7 |
| TP53 Regulates Transcription of Cell Death Genes | 0.045 | CASP1;CASP10 |
| Signaling by the B Cell Receptor (BCR) | 0.047 | PSME2;PSMB9;PSMB10;PSME1 |
| Signaling by SCF-KIT | 0.047 | STAT1 |
| Interleukin-12 signaling | 0.049 | PSME2;SERPINB2 |

The pathway analysis report was created by employing Reactome pathway database (v82) for species “Homo sapiens.” The listed pathways were curated by submitting up-regulated proteins altered with LPS-Cytokines and presented in Table 1.
