## Supplement Table 5 for "Red Algae-Derived Mineral Intervention to Counter Pro-inflammatory Activity in Human Colon Organoids"

**Supplement Table 5. Pathways associated with up-regulated proteins altered with Aquamin in the presence of LPS-Cytokines.**

| Pathways | Entities pValue | Mapped entities |
|---|---|---|
| Interferon Signaling | $1.11x10^{-16}$ | HLA-E;HLA-A;TRIM21;OAS3;HLA-G;PML;HLA-DRB4;EIF2AK2;STAT1;HLA-DRB1;HLA-DRB3;HLA-DPB1;B2M; HLA-DPA1;IFITM2;HLA-F;DDX58;MX1;GBP1;GBP2;HLA-C;IFI30;HLA-DRA;ISG15; HLA-B;ICAM1;OAS1;UBE2L6;UBA7;HLA-DRB5 |
| Interferon gamma signaling | $1.11x10^{-16}$ | HLA-F;HLA-E;HLA-A;TRIM21;OAS3;HLA-G;PML;GBP1;GBP2;HLA-DRB4;HLA-C;STAT1;IFI30;HLA-DRA; HLA-DRB1;HLA-B;HLA-DRB3;ICAM1;OAS1;HLA-DPB1;B2M;HLA-DPA1;HLA-DRB5 |
| Cytokine Signaling in Immune system | $1.11x10^{-16}$ | IL32;HLA-E;HLA-A;PSMB9;TRIM21;OAS3;HLA-G;PML;GSDMD;PSME1;MUC1;HLA-DRB4;EIF2AK2;STAT1; HLA-DRB1;HLA-DRB3;CD40;PSMB10;HLA-DPB1;B2M;HLA-DPA1;IFITM2;HLA-F;DDX58;PSME2;CASP1; MX1;GBP1;NOS2;GBP2;HLA-C;IFI30;HLA-DRA;ISG15;HLA-B;ICAM1;SERPINB2;OAS1;UBE2L6;UBA7; HLA-DRB5 |
| Immune System | $5.93x10^{-14}$ | IL32;IFI16;HLA-A;HLA-E;OAS3;GSDMD;EIF2AK2;TMBIM1;STAT1;HLA-DRB1;HLA-DRB3;HLA-DPB1; HLA-DPA1;MUC17;HLA-F;CASP1;HRNR;GBP1;RNF213;GBP2;BTN3A3;HLA-C;ANPEP;IFI30;CD55; HLA-DRA;OLFM4;FBXO6;OAS1;HLA-DMB;PSMB9;TRIM21;HLA-G;PML;PSME1;MUC1;HLA-DRB4; TAPBP;CD40;PSMB10;KRT1;B2M;IFITM2;DDX58;PSME2;MX1;NOS2;CTSV;S100A7;ISG15;FLG2; SERPINA1;HLA-B;ICAM1;SERPINB2;TAP1;CFB;TAP2;DCD;UBA7;CD74;UBE2L6;HLA-DRB5;LGMN |
| Interferon alpha/beta signaling | $2.16x10^{-10}$ | HLA-F;MX1;HLA-E;HLA-A;HLA-G;OAS3;HLA-C;GBP2;STAT1;ISG15;HLA-B;OAS1;IFITM2 |
| Antigen processing-Cross presentation | $3.85x10^{-9}$ | HLA-F;PSME2;HLA-E;HLA-A;PSMB9;HLA-G;PSME1;CTSV;HLA-C;HLA-B;TAPBP;PSMB10;B2M;TAP1;TAP2 |
| Adaptive Immune System | $5.75x10^{-9}$ | HLA-DMB;HLA-E;HLA-A;PSMB9;TRIM21;HLA-G;PSME1;HLA-DRB4;HLA-DRB1;HLA-DRB3;TAPBP;CD40;PSMB10; HLA-DPB1;B2M;HLA-DPA1;HLA-F;PSME2;RNF213;CTSV;HLA-C;BTN3A3;IFI30;HLA-DRA;HLA-B;FBXO6; ICAM1;TAP1;TAP2;CD74;UBE2L6;UBA7;HLA-DRB5;LGMN |
| ER-Phagosome pathway | $9.46x10^{-9}$ | HLA-F;PSME2;HLA-E;HLA-A;PSMB9;HLA-G;PSME1;HLA-C;HLA-B;TAPBP;PSMB10;B2M;TAP1;TAP2 |
| MHC class II antigen presentation | $1.64x10^{-7}$ | HLA-DMB;IFI30;HLA-DRA;HLA-DRB1;HLA-DRB3;HLA-DPB1;CD74;CTSV;HLA-DPA1;HLA-DRB4;LGMN;HLA-DRB5 |
| Downstream TCR signaling | $1.78x10^{-7}$ | PSME2;HLA-DRA;PSMB9;HLA-DRB1;HLA-DRB3;PSMB10;PSME1;HLA-DPB1;HLA-DPA1;HLA-DRB4;HLA-DRB5 |
| Antigen Presentation: Folding, assembly and peptide loading of class I MHC | $3.39x10^{-7}$ | HLA-F;HLA-A;HLA-E;HLA-B;TAPBP;HLA-G;B2M;TAP1;TAP2;HLA-C |
| Translocation of ZAP-70 to Immunological synapse | $4.48x10^{-7}$ | HLA-DRA;HLA-DRB1;HLA-DRB3;HLA-DPB1;HLA-DPA1;HLA-DRB5;HLA-DRB4 |
| Class I MHC mediated antigen processing & presentation | $4.76x10^{-7}$ | HLA-F;PSME2;HLA-E;HLA-A;PSMB9;TRIM21;HLA-G;PSME1;RNF213;CTSV;HLA-C; HLA-B;TAPBP;FBXO6;PSMB10;B2M;TAP1;TAP2;UBA7;UBE2L6 |
| Phosphorylation of CD3 and TCR zeta chains | $7.40x10^{-7}$ | HLA-DRA;HLA-DRB1;HLA-DRB3;HLA-DPB1;HLA-DPA1;HLA-DRB5;HLA-DRB4 |
| PD-1 signaling | $8.67x10^{-7}$ | HLA-DRA;HLA-DRB1;HLA-DRB3;HLA-DPB1;HLA-DPA1;HLA-DRB5;HLA-DRB4 |
| TCR signaling | $9.62x10^{-7}$ | PSME2;HLA-DRA;PSMB9;HLA-DRB1;HLA-DRB3;PSMB10;PSME1;HLA-DPB1;HLA-DPA1;HLA-DRB4;HLA-DRB5 |
| Antiviral mechanism by IFN-stimulated genes | $1.44x10^{-6}$ | EIF2AK2;DDX58;STAT1;MX1;ISG15;OAS3;OAS1;UBA7;UBE2L6 |
| Generation of second messenger molecules | $3.96x10^{-6}$ | HLA-DRA;HLA-DRB1;HLA-DRB3;HLA-DPB1;HLA-DPA1;HLA-DRB5;HLA-DRB4 |
| Endosomal/Vacuolar pathway | $5.55x10^{-6}$ | HLA-F;HLA-A;HLA-E;HLA-B;HLA-G;B2M;CTSV;HLA-C |
| SARS-CoV-2 activates/modulates innate and adaptive immune responses | $9.49x10^{-6}$ | HLA-F;DDX58;STAT1;HLA-A;HLA-E;ISG15;HLA-B;HLA-G;B2M;HLA-C |
| Innate Immune System | $9.95x10^{-6}$ | IFI16;HLA-E;PSMB9;TRIM21;GSDMD;PSME1;MUC1;TMBIM1;PSMB10;KRT1;B2M;MUC17;DDX58;PSME2;CASP1; HRNR;NOS2;CTSV;HLA-C;ANPEP;CD55;S100A7;ISG15;FLG2;SERPINA1;HLA-B;OLFM4;CFB;DCD;UBE2L6; UBA7;LGMN |
| ISG15 antiviral mechanism | $5.43x10^{-5}$ | EIF2AK2;DDX58;STAT1;MX1;ISG15;UBA7;UBE2L6 |

| | | |
|---|---|---|
| Costimulation by the CD28 family | $1.12x10^{-4}$ | HLA-DRA;HLA-DRB1;HLA-DRB3;HLA-DPB1;HLA-DPA1;HLA-DRB5;HLA-DRB4 |
| SARS-CoV-2-host interactions | $1.83x10^{-4}$ | HLA-F;DDX58;STAT1;HLA-A;HLA-E;ISG15;HLA-B;HLA-G;B2M;HLA-C |
| OAS antiviral response | $2.51x10^{-4}$ | DDX58;OAS3;OAS1 |
| Loss of MECP2 binding ability to 5hmC-DNA | $2.76x10^{-4}$ | MECP2 |
| SARS-CoV-2 Infection | $5.42x10^{-4}$ | HLA-F;DDX58;STAT1;HLA-E;HLA-A;ISG15;HLA-B;HLA-G;PARP9;B2M;HLA-C |
| Negative regulators of DDX58/IFIH1 signaling | $8.65x10^{-4}$ | DDX58;ISG15;UBE2L6;UBA7 |
| MECP2 regulates transcription of genes involved in GABA signaling | 0.001 | MECP2 |
| Loss of MECP2 binding ability to 5mC-DNA | 0.001 | MECP2 |
| KEAP1-NFE2L2 pathway | 0.002 | PSME2;GSTA1;PSMB9;TRIM21;PSMB10;PSME1 |
| Loss of phosphorylation of MECP2 at T308 | 0.002 | MECP2 |
| Fructose catabolism | 0.002 | ALDOB;ALDH1A1 |
| Regulation of RUNX2 expression and activity | 0.002 | STAT1;PSME2;PSMB9;PSMB10;PSME1 |
| MECP2 regulates transcription factors | 0.002 | MECP2 |
| Nuclear events mediated by NFE2L2 | 0.003 | PSME2;GSTA1;PSMB9;PSMB10;PSME1 |
| Regulation of activated PAK-2p34 by proteasome mediated degradation | 0.003 | PSME2;PSMB9;PSMB10;PSME1 |
| Cross-presentation of soluble exogenous antigens (endosomes) | 0.003 | PSME2;PSMB9;PSMB10;PSME1 |
| Fructose metabolism | 0.003 | ALDOB;ALDH1A1 |
| Regulation of ornithine decarboxylase (ODC) | 0.003 | PSME2;PSMB9;PSMB10;PSME1 |
| p53-Independent DNA Damage Response | 0.004 | PSME2;PSMB9;PSMB10;PSME1 |
| Ubiquitin Mediated Degradation of Phosphorylated Cdc25A | 0.004 | PSME2;PSMB9;PSMB10;PSME1 |
| p53-Independent G1/S DNA damage checkpoint | 0.004 | PSME2;PSMB9;PSMB10;PSME1 |
| GSK3B and BTRC:CUL1-mediated-degradation of NFE2L2 | 0.004 | PSME2;PSMB9;PSMB10;PSME1 |
| Autodegradation of the E3 ubiquitin ligase COP1 | 0.004 | PSME2;PSMB9;PSMB10;PSME1 |
| Ubiquitin-dependent degradation of Cyclin D | 0.004 | PSME2;PSMB9;PSMB10;PSME1 |
| Vpu mediated degradation of CD4 | 0.004 | PSME2;PSMB9;PSMB10;PSME1 |
| Regulation of Apoptosis | 0.004 | PSME2;PSMB9;PSMB10;PSME1 |
| Signaling by Interleukins | 0.004 | STAT1;PSME2;CASP1;IL32;PSMB9;ICAM1;SERPINB2;PSMB10;GSDMD;PSME1;MUC1;NOS2 |
| Loss of MECP2 binding ability to the NCoR/SMRT complex | 0.004 | MECP2 |
| FBXL7 down-regulates AURKA during mitotic entry and in early mitosis | 0.004 | PSME2;PSMB9;PSMB10;PSME1 |
| SCF-beta-TrCP mediated degradation of Emi1 | 0.004 | PSME2;PSMB9;PSMB10;PSME1 |
| Degradation of AXIN | 0.004 | PSME2;PSMB9;PSMB10;PSME1 |

| | | |
|---|---|---|
| Negative regulation of NOTCH4 signaling | 0.004 | PSME2;PSMB9;PSMB10;PSME1 |
| Regulation of RUNX3 expression and activity | 0.004 | PSME2;PSMB9;PSMB10;PSME1 |
| Programmed Cell Death | 0.005 | DSG2;PSME2;CASP1;CASP7;PSMB9;PSMB10;GSDMD;PSME1 |
| Formation of the cornified envelope | 0.005 | DSG2;KRT16;KRT9;KRT2;KRT6A;KRT1 |
| AUF1 (hnRNP D0) binds and destabilizes mRNA | 0.005 | PSME2;PSMB9;PSMB10;PSME1 |
| Vif-mediated degradation of APOBEC3G | 0.005 | PSME2;PSMB9;PSMB10;PSME1 |
| Hh mutants are degraded by ERAD | 0.005 | PSME2;PSMB9;PSMB10;PSME1 |
| Neutrophil degranulation | 0.005 | HRNR;GSDMD;HLA-C;TMBIM1;ANPEP;CD55;S100A7;FLG2;OLFM4;SERPINA1;HLA-B;B2M;KRT1 |
| Degradation of DVL | 0.005 | PSME2;PSMB9;PSMB10;PSME1 |
| Stabilization of p53 | 0.005 | PSME2;PSMB9;PSMB10;PSME1 |
| MECP2 regulates transcription of neuronal ligands | 0.005 | MECP2 |
| Interleukin-1 processing | 0.005 | CASP1;GSDMD |
| Hh mutants abrogate ligand secretion | 0.006 | PSME2;PSMB9;PSMB10;PSME1 |
| NIK-->noncanonical NF-kB signaling | 0.006 | PSME2;PSMB9;PSMB10;PSME1 |
| Metabolism of polyamines | 0.006 | PSME2;PSMB9;PSMB10;PSME1 |
| Degradation of GLI1 by the proteasome | 0.006 | PSME2;PSMB9;PSMB10;PSME1 |
| GLI3 is processed to GLI3R by the proteasome | 0.006 | PSME2;PSMB9;PSMB10;PSME1 |
| Degradation of GLI2 by the proteasome | 0.006 | PSME2;PSMB9;PSMB10;PSME1 |
| SCF(Skp2)-mediated degradation of p27/p21 | 0.006 | PSME2;PSMB9;PSMB10;PSME1 |
| Dectin-1 mediated noncanonical NF-kB signaling | 0.006 | PSME2;PSMB9;PSMB10;PSME1 |
| Defective CFTR causes cystic fibrosis | 0.006 | PSME2;PSMB9;PSMB10;PSME1 |
| Apoptosis | 0.006 | DSG2;PSME2;CASP7;PSMB9;PSMB10;GSDMD;PSME1 |
| Termination of translesion DNA synthesis | 0.007 | ISG15;UBE2L6;UBA7 |
| Autodegradation of Cdh1 by Cdh1:APC/C | 0.007 | PSME2;PSMB9;PSMB10;PSME1 |
| Asymmetric localization of PCP proteins | 0.007 | PSME2;PSMB9;PSMB10;PSME1 |
| TNFR2 non-canonical NF-kB pathway | 0.008 | PSME2;PSMB9;CD40;PSMB10;PSME1 |
| Metallothioneins bind metals | 0.008 | MT1E;MT1H |
| Hedgehog ligand biogenesis | 0.008 | PSME2;PSMB9;PSMB10;PSME1 |
| Oxygen-dependent proline hydroxylation of Hypoxia-inducible Factor Alpha | 0.008 | PSME2;PSMB9;PSMB10;PSME1 |
| p53-Dependent G1 DNA Damage Response | 0.008 | PSME2;PSMB9;PSMB10;PSME1 |
| p53-Dependent G1/S DNA damage checkpoint | 0.008 | PSME2;PSMB9;PSMB10;PSME1 |
| Activation of NF-kappaB in B cells | 0.009 | PSME2;PSMB9;PSMB10;PSME1 |
| APC/C:Cdc20 mediated degradation of Securin | 0.009 | PSME2;PSMB9;PSMB10;PSME1 |

| | | |
|---|---|---|
| G1/S DNA Damage Checkpoints | 0.009 | PSME2;PSMB9;PSMB10;PSME1 |
| Immunoregulatory interactions between a Lymphoid and a non-Lymphoid cell | 0.009 | HLA-F;HLA-A;HLA-E;HLA-B;CD40;ICAM1;HLA-G;B2M;HLA-C |
| Regulation of RAS by GAPs | 0.010 | PSME2;PSMB9;PSMB10;PSME1 |
| Regulation of PTEN stability and activity | 0.010 | PSME2;PSMB9;PSMB10;PSME1 |
| Orc1 removal from chromatin | 0.011 | PSME2;PSMB9;PSMB10;PSME1 |
| Disorders of Developmental Biology | 0.011 | MECP2 |
| Pervasive developmental disorders | 0.011 | MECP2 |
| Loss of function of MECP2 in Rett syndrome | 0.011 | MECP2 |
| Disorders of Nervous System Development | 0.011 | MECP2 |
| Trafficking and processing of endosomal TLR | 0.011 | CTSV;LGMN |
| Interleukin-1 family signaling | 0.011 | PSME2;CASP1;PSMB9;PSMB10;GSDMD;PSME1 |
| Translesion synthesis by Y family DNA polymerases bypasses lesions on DNA template | 0.012 | ISG15;UBE2L6;UBA7 |
| Cdc20:Phospho-APC/C mediated degradation of Cyclin A | 0.012 | PSME2;PSMB9;PSMB10;PSME1 |
| CDK-mediated phosphorylation and removal of Cdc6 | 0.012 | PSME2;PSMB9;PSMB10;PSME1 |
| SARS-CoV Infections | 0.012 | HLA-F;DDX58;STAT1;HLA-E;HLA-A;ISG15;HLA-B;HLA-G;PARP9;B2M;HLA-C |
| Hereditary fructose intolerance | 0.012 | ALDOB |
| Defective SLC11A2 causes hypochromic microcytic anemia, with iron overload 1 (AHMIO1) | 0.012 | SLC11A2 |
| Antigen processing: Ubiquitination & Proteasome degradation | 0.012 | PSME2;PSMB9;FBXO6;TRIM21;PSMB10;PSME1;RNF213;UBA7;UBE2L6 |
| Signaling by NOTCH | 0.012 | STAT1;PSME2;PLXND1;PSMB9;PSMB10;PSME1;MDK |
| APC/C:Cdh1 mediated degradation of Cdc20 and other APC/C:Cdh1 targeted proteins in late mitosis/early G1 | 0.012 | PSME2;PSMB9;PSMB10;PSME1 |
| APC:Cdc20 mediated degradation of cell cycle proteins prior to satisfation of the cell cycle checkpoint | 0.012 | PSME2;PSMB9;PSMB10;PSME1 |
| Response to metal ions | 0.012 | MT1E;MT1H |
| APC/C:Cdc20 mediated degradation of mitotic proteins | 0.013 | PSME2;PSMB9;PSMB10;PSME1 |
| Cellular response to hypoxia | 0.013 | PSME2;PSMB9;PSMB10;PSME1 |
| Activation of APC/C and APC/C:Cdc20 mediated degradation of mitotic proteins | 0.014 | PSME2;PSMB9;PSMB10;PSME1 |
| Interleukin-6 signaling | 0.014 | STAT1 |
| ABC transporter disorders | 0.014 | PSME2;PSMB9;PSMB10;PSME1 |

| | | |
|---|---|---|
| The role of GTSE1 in G2/M progression after G2 checkpoint | 0.015 | PSME2;PSMB9;PSMB10;PSME1 |
| Transcriptional regulation by RUNX2 | 0.015 | STAT1;PSME2;PSMB9;PSMB10;PSME1 |
| STING mediated induction of host immune responses | 0.016 | IFI16;TRIM21 |
| Regulation of APC/C activators between G1/S and early anaphase | 0.016 | PSME2;PSMB9;PSMB10;PSME1 |
| Downstream signaling events of B Cell Receptor (BCR) | 0.017 | PSME2;PSMB9;PSMB10;PSME1 |
| Cyclin E associated events during G1/S transition | 0.018 | PSME2;PSMB9;PSMB10;PSME1 |
| Signaling by NOTCH4 | 0.018 | PSME2;PSMB9;PSMB10;PSME1 |
| Degradation of beta-catenin by the destruction complex | 0.018 | PSME2;PSMB9;PSMB10;PSME1 |
| C-type lectin receptors (CLRs) | 0.018 | MUC17;PSME2;PSMB9;PSMB10;PSME1;MUC1 |
| DDX58/IFIH1-mediated induction of interferon-alpha/beta | 0.019 | DDX58;ISG15;UBE2L6;UBA7 |
| Hedgehog 'on' state | 0.019 | PSME2;PSMB9;PSMB10;PSME1 |
| Cyclin A:Cdk2-associated events at S phase entry | 0.019 | PSME2;PSMB9;PSMB10;PSME1 |
| MECP2 regulates neuronal receptors and channels | 0.020 | MECP2 |
| Signaling by cytosolic FGFR1 fusion mutants | 0.020 | STAT1;ZMYM2 |
| DNA Damage Bypass | 0.020 | ISG15;UBE2L6;UBA7 |
| APC/C-mediated degradation of cell cycle proteins | 0.021 | PSME2;PSMB9;PSMB10;PSME1 |
| Regulation of mitotic cell cycle | 0.021 | PSME2;PSMB9;PSMB10;PSME1 |
| Regulation of mRNA stability by proteins that bind AU-rich elements | 0.021 | PSME2;PSMB9;PSMB10;PSME1 |
| Disorders of transmembrane transporters | 0.022 | CP;PSME2;PSMB9;PSMB10;PSME1;SLC11A2 |
| Defective GALNT3 causes HFTC | 0.022 | MUC17;MUC1 |
| Defective GALNT12 causes CRCS1 | 0.022 | MUC17;MUC1 |
| MAPK6/MAPK4 signaling | 0.022 | PSME2;PSMB9;PSMB10;PSME1 |
| Defective SLC40A1 causes hemochromatosis 4 (HFE4) (macrophages) | 0.023 | CP |
| Defective CP causes aceruloplasminemia (ACERULOP) | 0.023 | CP |
| Inhibition of PKR | 0.023 | EIF2AK2 |
| Defective C1GALT1C1 causes TNPS | 0.024 | MUC17;MUC1 |
| Aflatoxin activation and detoxification | 0.024 | CYP3A4;ACY3 |

| | | |
|---|---|---|
| Nuclear events stimulated by ALK signaling in cancer | 0.024 | MECP2 |
| Apoptotic factor-mediated response | 0.024 | CASP7;GSDMD |
| Switching of origins to a post-replicative state | 0.025 | PSME2;PSMB9;PSMB10;PSME1 |
| PCP/CE pathway | 0.025 | PSME2;PSMB9;PSMB10;PSME1 |
| PTEN Regulation | 0.026 | PSME2;PSMB9;PML;PSMB10;PSME1 |
| UCH proteinases | 0.028 | PSME2;PSMB9;PSMB10;PSME1 |
| Transcriptional regulation by RUNX3 | 0.028 | PSME2;PSMB9;PSMB10;PSME1 |
| Signaling by phosphorylated juxtamembrane, extracellular and kinase domain KIT mutants | 0.029 | STAT1 |
| Signaling by KIT in disease | 0.029 | STAT1 |
| Host Interactions of HIV factors | 0.029 | PSME2;PSMB9;PSMB10;PSME1;B2M |
| Interleukin-12 family signaling | 0.030 | STAT1;PSME2;SERPINB2 |
| CLEC7A (Dectin-1) signaling | 0.030 | PSME2;PSMB9;PSMB10;PSME1 |
| Cellular response to chemical stress | 0.030 | PSME2;GSTA1;PSMB9;TRIM21;PSMB10;PSME1 |
| Signaling by ALK fusions and activated point mutants | 0.031 | MECP2;RNF213 |
| Signaling by ALK in cancer | 0.031 | MECP2;RNF213 |
| RUNX1 regulates transcription of genes involved in differentiation of HSCs | 0.031 | PSME2;PSMB9;PSMB10;PSME1 |
| The AIM2 inflammasome | 0.035 | CASP1 |
| ABC-family proteins mediated transport | 0.035 | PSME2;PSMB9;PSMB10;PSME1 |
| NOTCH3 Intracellular Domain Regulates Transcription | 0.036 | STAT1;PLXND1 |
| Transcriptional regulation by RUNX1 | 0.039 | PSME2;PSMB9;PML;PSMB10;PSME1;CTSV |
| Termination of O-glycan biosynthesis | 0.039 | MUC17;MUC1 |
| Regulation of IFNA/IFNB signaling | 0.039 | STAT1 |
| Metal ion SLC transporters | 0.039 | CP;SLC11A2 |
| Pyroptosis | 0.041 | CASP1;GSDMD |
| Cell recruitment (pro-inflammatory response) | 0.041 | CASP1;GSDMD |
| Purinergic signaling in leishmaniasis infection | 0.041 | CASP1;GSDMD |
| Interleukin-4 and Interleukin-13 signaling | 0.044 | STAT1;ICAM1;NOS2;MUC1 |
| Interleukin-6 family signaling | 0.044 | STAT1 |
| Assembly of the pre-replicative complex | 0.045 | PSME2;PSMB9;PSMB10;PSME1 |
| Keratinization | 0.046 | DSG2;KRT16;KRT9;KRT2;KRT6A;KRT1 |
| Inhibition of nitric oxide production | 0.046 | NOS2 |

| | | |
|---|---|---|
| Diseases of signal transduction by growth factor receptors and second messengers | 0.047 | STAT1;PSME2;ZMYM2;PSMB9;MECP2;PSMB10;PSME1;RNF213 |
| Hedgehog 'off' state | 0.048 | PSME2;PSMB9;PSMB10;PSME1 |
| Interleukin-1 signaling | 0.049 | PSME2;PSMB9;PSMB10;PSME1 |
| DAP12 signaling | 0.050 | HLA-E;B2M |

The pathway analysis report was created by employing Reactome pathway database (v82) for species “Homo sapiens.” The listed pathways were curated by submitting up-regulated proteins altered with LPS-Cytokines in the presence of Aquamin.
