## Supplement Table 6 for "Red Algae-Derived Mineral Intervention to Counter Pro-inflammatory Activity in Human Colon Organoids"

**Supplement Table 6. Pathways associated with down-regulated proteins altered with LPS-Cytokines.**

| Pathways | Entities pValue | Mapped entities |
|---|---|---|
| Nuclear events mediated by NFE2L2 | $1.47\times10^{-7}$ | PSMB7;GSTA1;PSMB5;PSMB6 |
| KEAP1-NFE2L2 pathway | $4.21\times10^{-7}$ | PSMB7;GSTA1;PSMB5;PSMB6 |
| Regulation of activated PAK-2p34 by proteasome mediated degradation | $4.40\times10^{-6}$ | PSMB7;PSMB5;PSMB6 |
| Cross-presentation of soluble exogenous antigens (endosomes) | $4.40\times10^{-6}$ | PSMB7;PSMB5;PSMB6 |
| Regulation of ornithine decarboxylase (ODC) | $4.67\times10^{-6}$ | PSMB7;PSMB5;PSMB6 |
| p53-Independent G1/S DNA damage checkpoint | $4.94\times10^{-6}$ | PSMB7;PSMB5;PSMB6 |
| GSK3B and BTRC:CUL1-mediated-degradation of NFE2L2 | $4.94\times10^{-6}$ | PSMB7;PSMB5;PSMB6 |
| Ubiquitin Mediated Degradation of Phosphorylated Cdc25A | $4.94\times10^{-6}$ | PSMB7;PSMB5;PSMB6 |
| p53-Independent DNA Damage Response | $4.94\times10^{-6}$ | PSMB7;PSMB5;PSMB6 |
| Autodegradation of the E3 ubiquitin ligase COP1 | $4.94\times10^{-6}$ | PSMB7;PSMB5;PSMB6 |
| Ubiquitin-dependent degradation of Cyclin D | $4.94\times10^{-6}$ | PSMB7;PSMB5;PSMB6 |
| Vpu mediated degradation of CD4 | $5.23\times10^{-6}$ | PSMB7;PSMB5;PSMB6 |
| Regulation of Apoptosis | $5.23\times10^{-6}$ | PSMB7;PSMB5;PSMB6 |
| Cellular response to chemical stress | $5.38\times10^{-6}$ | PSMB7;GSTA1;PSMB5;PSMB6 |
| FBXL7 down-regulates AURKA during mitotic entry and in early mitosis | $5.84\times10^{-6}$ | PSMB7;PSMB5;PSMB6 |
| SCF-beta-TrCP mediated degradation of Emi1 | $5.84\times10^{-6}$ | PSMB7;PSMB5;PSMB6 |
| Degradation of AXIN | $5.84\times10^{-6}$ | PSMB7;PSMB5;PSMB6 |
| Negative regulation of NOTCH4 signaling | $5.84\times10^{-6}$ | PSMB7;PSMB5;PSMB6 |
| Regulation of RUNX3 expression and activity | $5.84\times10^{-6}$ | PSMB7;PSMB5;PSMB6 |
| AUF1 (hnRNP D0) binds and destabilizes mRNA | $6.17\times10^{-6}$ | PSMB7;PSMB5;PSMB6 |
| Vif-mediated degradation of APOBEC3G | $6.17\times10^{-6}$ | PSMB7;PSMB5;PSMB6 |
| Hh mutants are degraded by ERAD | $6.17\times10^{-6}$ | PSMB7;PSMB5;PSMB6 |
| Degradation of DVL | $6.50\times10^{-6}$ | PSMB7;PSMB5;PSMB6 |
| Stabilization of p53 | $6.50\times10^{-6}$ | PSMB7;PSMB5;PSMB6 |
| Hh mutants abrogate ligand secretion | $7.20\times10^{-6}$ | PSMB7;PSMB5;PSMB6 |
| NIK-->noncanonical NF-kB signaling | $7.20\times10^{-6}$ | PSMB7;PSMB5;PSMB6 |
| Metabolism of polyamines | $7.20\times10^{-6}$ | PSMB7;PSMB5;PSMB6 |
| Degradation of GLI1 by the proteasome | $7.57\times10^{-6}$ | PSMB7;PSMB5;PSMB6 |
| GLI3 is processed to GLI3R by the proteasome | $7.57\times10^{-6}$ | PSMB7;PSMB5;PSMB6 |
| Degradation of GLI2 by the proteasome | $7.57\times10^{-6}$ | PSMB7;PSMB5;PSMB6 |
| SCF(Skp2)-mediated degradation of p27/p21 | $7.57\times10^{-6}$ | PSMB7;PSMB5;PSMB6 |
| Dectin-1 mediated noncanonical NF-kB signaling | $7.57\times10^{-6}$ | PSMB7;PSMB5;PSMB6 |
| Defective CFTR causes cystic fibrosis | $7.96\times10^{-6}$ | PSMB7;PSMB5;PSMB6 |
| Autodegradation of Cdh1 by Cdh1:APC/C | $9.18\times10^{-6}$ | PSMB7;PSMB5;PSMB6 |
| Asymmetric localization of PCP proteins | $9.18\times10^{-6}$ | PSMB7;PSMB5;PSMB6 |
| Hedgehog ligand biogenesis | $9.62\times10^{-6}$ | PSMB7;PSMB5;PSMB6 |
| Oxygen-dependent proline hydroxylation of Hypoxia-inducible Factor Alpha | $1.01\times10^{-5}$ | PSMB7;PSMB5;PSMB6 |
| p53-Dependent G1 DNA Damage Response | $1.01\times10^{-5}$ | PSMB7;PSMB5;PSMB6 |
| p53-Dependent G1/S DNA damage checkpoint | $1.01\times10^{-5}$ | PSMB7;PSMB5;PSMB6 |
| Activation of NF-kappaB in B cells | $1.05\times10^{-5}$ | PSMB7;PSMB5;PSMB6 |
| APC/C:Cdc20 mediated degradation of Securin | $1.10\times10^{-5}$ | PSMB7;PSMB5;PSMB6 |
| G1/S DNA Damage Checkpoints | $1.10\times10^{-5}$ | PSMB7;PSMB5;PSMB6 |
| Regulation of RAS by GAPs | $1.15\times10^{-5}$ | PSMB7;PSMB5;PSMB6 |

| | | |
|---|---|---|
| Regulation of PTEN stability and activity | $1.15\times10^{-5}$ | PSMB7;PSMB5;PSMB6 |
| Orc1 removal from chromatin | $1.25\times10^{-5}$ | PSMB7;PSMB5;PSMB6 |
| Cdc20:Phospho-APC/C mediated degradation of Cyclin A | $1.36\times10^{-5}$ | PSMB7;PSMB5;PSMB6 |
| CDK-mediated phosphorylation and removal of Cdc6 | $1.36\times10^{-5}$ | PSMB7;PSMB5;PSMB6 |
| APC/C:Cdh1 mediated degradation of Cdc20 and other APC/C:Cdh1 targeted proteins in late mitosis/early G1 | $1.41\times10^{-5}$ | PSMB7;PSMB5;PSMB6 |
| APC:Cdc20 mediated degradation of cell cycle proteins prior to satisfation of the cell cycle checkpoint | $1.41\times10^{-5}$ | PSMB7;PSMB5;PSMB6 |
| Regulation of RUNX2 expression and activity | $1.41\times10^{-5}$ | PSMB7;PSMB5;PSMB6 |
| APC/C:Cdc20 mediated degradation of mitotic proteins | $1.53\times10^{-5}$ | PSMB7;PSMB5;PSMB6 |
| Cellular response to hypoxia | $1.53\times10^{-5}$ | PSMB7;PSMB5;PSMB6 |
| Activation of APC/C and APC/C:Cdc20 mediated degradation of mitotic proteins | $1.59\times10^{-5}$ | PSMB7;PSMB5;PSMB6 |
| ABC transporter disorders | $1.65\times10^{-5}$ | PSMB7;PSMB5;PSMB6 |
| The role of GTSE1 in G2/M progression after G2 checkpoint | $1.72\times10^{-5}$ | PSMB7;PSMB5;PSMB6 |
| Regulation of APC/C activators between G1/S and early anaphase | $1.85\times10^{-5}$ | PSMB7;PSMB5;PSMB6 |
| Downstream signaling events of B Cell Receptor (BCR) | $1.92\times10^{-5}$ | PSMB7;PSMB5;PSMB6 |
| Cyclin E associated events during G1/S transition | $1.99\times10^{-5}$ | PSMB7;PSMB5;PSMB6 |
| Signaling by NOTCH4 | $1.99\times10^{-5}$ | PSMB7;PSMB5;PSMB6 |
| Degradation of beta-catenin by the destruction complex | $1.99\times10^{-5}$ | PSMB7;PSMB5;PSMB6 |
| Hedgehog 'on' state | $2.14\times10^{-5}$ | PSMB7;PSMB5;PSMB6 |
| Cyclin A:Cdk2-associated events at S phase entry | $2.14\times10^{-5}$ | PSMB7;PSMB5;PSMB6 |
| Regulation of mitotic cell cycle | $2.37\times10^{-5}$ | PSMB7;PSMB5;PSMB6 |
| APC/C-mediated degradation of cell cycle proteins | $2.37\times10^{-5}$ | PSMB7;PSMB5;PSMB6 |
| Regulation of mRNA stability by proteins that bind AU-rich elements | $2.37\times10^{-5}$ | PSMB7;PSMB5;PSMB6 |
| MAPK6/MAPK4 signaling | $2.45\times10^{-5}$ | PSMB7;PSMB5;PSMB6 |
| Switching of origins to a post-replicative state | $2.70\times10^{-5}$ | PSMB7;PSMB5;PSMB6 |
| PCP/CE pathway | $2.70\times10^{-5}$ | PSMB7;PSMB5;PSMB6 |
| UCH proteinases | $3.07\times10^{-5}$ | PSMB7;PSMB5;PSMB6 |
| Transcriptional regulation by RUNX3 | $3.07\times10^{-5}$ | PSMB7;PSMB5;PSMB6 |
| CLEC7A (Dectin-1) signaling | $3.26\times10^{-5}$ | PSMB7;PSMB5;PSMB6 |
| RUNX1 regulates transcription of genes involved in differentiation of HSCs | $3.36\times10^{-5}$ | PSMB7;PSMB5;PSMB6 |
| TNFR2 non-canonical NF-kB pathway | $3.67\times10^{-5}$ | PSMB7;PSMB5;PSMB6 |
| ABC-family proteins mediated transport | $3.78\times10^{-5}$ | PSMB7;PSMB5;PSMB6 |
| Assembly of the pre-replicative complex | $4.84\times10^{-5}$ | PSMB7;PSMB5;PSMB6 |
| Hedgehog 'off' state | $5.11\times10^{-5}$ | PSMB7;PSMB5;PSMB6 |
| Interleukin-1 signaling | $5.24\times10^{-5}$ | PSMB7;PSMB5;PSMB6 |
| Downstream TCR signaling | $5.38\times10^{-5}$ | PSMB7;PSMB5;PSMB6 |
| Synthesis of DNA | $6.09\times10^{-5}$ | PSMB7;PSMB5;PSMB6 |
| Transcriptional regulation by RUNX2 | $6.09\times10^{-5}$ | PSMB7;PSMB5;PSMB6 |
| DNA Replication Pre-Initiation | $7.36\times10^{-5}$ | PSMB7;PSMB5;PSMB6 |
| G1/S Transition | $7.70\times10^{-5}$ | PSMB7;PSMB5;PSMB6 |
| TCR signaling | $8.99\times10^{-5}$ | PSMB7;PSMB5;PSMB6 |
| PTEN Regulation | $9.18\times10^{-5}$ | PSMB7;PSMB5;PSMB6 |
| Host Interactions of HIV factors | $1.02\times10^{-4}$ | PSMB7;PSMB5;PSMB6 |
| Beta-catenin independent WNT signaling | $1.02\times10^{-4}$ | PSMB7;PSMB5;PSMB6 |
| Mitotic G1 phase and G1/S transition | $1.13\times10^{-4}$ | PSMB7;PSMB5;PSMB6 |
| Signaling by Hedgehog | $1.15\times10^{-4}$ | PSMB7;PSMB5;PSMB6 |
| G2/M Checkpoints | $1.15\times10^{-4}$ | PSMB7;PSMB5;PSMB6 |

| | | |
|---|---|---|
| Interleukin-1 family signaling | $1.27\times10^{-4}$ | PSMB7;PSMB5;PSMB6 |
| DNA Replication | $1.32\times10^{-4}$ | PSMB7;PSMB5;PSMB6 |
| ER-Phagosome pathway | $1.39\times10^{-4}$ | PSMB7;PSMB5;PSMB6 |
| S Phase | $1.47\times10^{-4}$ | PSMB7;PSMB5;PSMB6 |
| FCERI mediated NF-kB activation | $1.58\times10^{-4}$ | PSMB7;PSMB5;PSMB6 |
| Regulation of expression of SLITs and ROBOs | $1.72\times10^{-4}$ | PSMB7;PSMB5;PSMB6 |
| C-type lectin receptors (CLRs) | $1.78\times10^{-4}$ | PSMB7;PSMB5;PSMB6 |
| Antigen processing-Cross presentation | $1.84\times10^{-4}$ | PSMB7;PSMB5;PSMB6 |
| Signaling by the B Cell Receptor (BCR) | $1.84\times10^{-4}$ | PSMB7;PSMB5;PSMB6 |
| Disorders of transmembrane transporters | $2.00\times10^{-4}$ | PSMB7;PSMB5;PSMB6 |
| Apoptosis | $2.03\times10^{-4}$ | PSMB7;PSMB5;PSMB6 |
| Separation of Sister Chromatids | $2.34\times10^{-4}$ | PSMB7;PSMB5;PSMB6 |
| G2/M Transition | $2.60\times10^{-4}$ | PSMB7;PSMB5;PSMB6 |
| Mitotic G2-G2/M phases | $2.68\times10^{-4}$ | PSMB7;PSMB5;PSMB6 |
| TCF dependent signaling in response to WNT | $2.76\times10^{-4}$ | PSMB7;PSMB5;PSMB6 |
| Ub-specific processing proteases | $2.88\times10^{-4}$ | PSMB7;PSMB5;PSMB6 |
| Signaling by NOTCH | $2.92\times10^{-4}$ | PSMB7;PSMB5;PSMB6 |
| Transcriptional regulation by RUNX1 | $3.01\times10^{-4}$ | PSMB7;PSMB5;PSMB6 |
| Programmed Cell Death | $3.41\times10^{-4}$ | PSMB7;PSMB5;PSMB6 |
| Signaling by ROBO receptors | $3.45\times10^{-4}$ | PSMB7;PSMB5;PSMB6 |
| Fc epsilon receptor (FCERI) signaling | $3.45\times10^{-4}$ | PSMB7;PSMB5;PSMB6 |
| Developmental Biology | $3.62\times10^{-4}$ | PSMB7;PI3;PSMB5;PSMB6;PCK1 |
| Mitotic Anaphase | $4.41\times10^{-4}$ | PSMB7;PSMB5;PSMB6 |
| Mitotic Metaphase and Anaphase | $4.46\times10^{-4}$ | PSMB7;PSMB5;PSMB6 |
| Neddylation | $4.74\times10^{-4}$ | PSMB7;PSMB5;PSMB6 |
| HIV Infection | $4.86\times10^{-4}$ | PSMB7;PSMB5;PSMB6 |
| Innate Immune System | $5.06\times10^{-4}$ | PSMB7;SLPI;PI3;PSMB5;PSMB6 |
| Cell Cycle Checkpoints | $6.73\times10^{-4}$ | PSMB7;PSMB5;PSMB6 |
| PIP3 activates AKT signaling | $7.24\times10^{-4}$ | PSMB7;PSMB5;PSMB6 |
| Deubiquitination | $7.39\times10^{-4}$ | PSMB7;PSMB5;PSMB6 |
| RAF/MAP kinase cascade | $8.10\times10^{-4}$ | PSMB7;PSMB5;PSMB6 |
| Signaling by WNT | $8.67\times10^{-4}$ | PSMB7;PSMB5;PSMB6 |
| MAPK1/MAPK3 signaling | $8.67\times10^{-4}$ | PSMB7;PSMB5;PSMB6 |
| Antigen processing: Ubiquitination & Proteasome degradation | $9.54\times10^{-4}$ | PSMB7;PSMB5;PSMB6 |
| Cellular responses to stress | 0.001 | PSMB7;GSTA1;PSMB5;PSMB6 |
| Intracellular signaling by second messengers | 0.001 | PSMB7;PSMB5;PSMB6 |
| Cellular responses to stimuli | 0.001 | PSMB7;GSTA1;PSMB5;PSMB6 |
| MAPK family signaling cascades | 0.001 | PSMB7;PSMB5;PSMB6 |
| Metabolism of amino acids and derivatives | 0.002 | PSMB7;PSMB5;PSMB6 |
| M Phase | 0.002 | PSMB7;PSMB5;PSMB6 |
| Drug ADME | 0.002 | GSTA1;PCK1 |
| Class I MHC mediated antigen processing & presentation | 0.003 | PSMB7;PSMB5;PSMB6 |
| Diseases of signal transduction by growth factor receptors and second messengers | 0.003 | PSMB7;PSMB5;PSMB6 |
| Signaling by Interleukins | 0.003 | PSMB7;PSMB5;PSMB6 |
| NR1H2 & NR1H3 regulate gene expression linked to gluconeogenesis | 0.003 | PCK1 |
| Abacavir metabolism | 0.003 | PCK1 |
| Cell Cycle, Mitotic | 0.005 | PSMB7;PSMB5;PSMB6 |
| Axon guidance | 0.005 | PSMB7;PSMB5;PSMB6 |
| Nervous system development | 0.006 | PSMB7;PSMB5;PSMB6 |

| | | |
|---|---|---|
| Generic Transcription Pathway | 0.007 | PSMB7;PSMB5;PSMB6;PCK1 |
| Abacavir ADME | 0.007 | PCK1 |
| Metabolism | 0.007 | PSMB7;GSTA1;PSMB5;PSMB6;PCK1 |
| Cell Cycle | 0.009 | PSMB7;PSMB5;PSMB6 |
| Metabolism of RNA | 0.009 | PSMB7;PSMB5;PSMB6 |
| Immune System | 0.009 | PSMB7;SLPI;PI3;PSMB5;PSMB6 |
| RNA Polymerase II Transcription | 0.009 | PSMB7;PSMB5;PSMB6;PCK1 |
| Heme degradation | 0.011 | GSTA1 |
| Transport of small molecules | 0.011 | PSMB7;PSMB5;PSMB6 |
| Gene expression (Transcription) | 0.013 | PSMB7;PSMB5;PSMB6;PCK1 |
| Cytokine Signaling in Immune system | 0.015 | PSMB7;PSMB5;PSMB6 |
| Azathioprine ADME | 0.016 | GSTA1 |
| Signal Transduction | 0.017 | RGS16;PSMB7;PSMB5;PSMB6;PCK1 |
| Metabolism of porphyrins | 0.020 | GSTA1 |
| FOXO-mediated transcription of oxidative stress, metabolic and neuronal genes | 0.020 | PCK1 |
| Adaptive Immune System | 0.023 | PSMB7;PSMB5;PSMB6 |
| Gluconeogenesis | 0.023 | PCK1 |
| Glutathione conjugation | 0.025 | GSTA1 |
| NR1H2 and NR1H3-mediated signaling | 0.033 | PCK1 |
| G alpha (z) signalling events | 0.033 | RGS16 |
| Neutrophil degranulation | 0.041 | PSMB7;SLPI |
| FOXO-mediated transcription | 0.045 | PCK1 |

The pathway analysis report was created by employing Reactome pathway database (v82) for species “Homo sapiens.” The listed pathways were curated by submitting down-regulated proteins altered with LPS-Cytokines and presented in Table 3.
