## Supplement Table 7 for "Red Algae-Derived Mineral Intervention to Counter Pro-inflammatory Activity in Human Colon Organoids"

**Supplement Table 7. Top pathways associated with up-regulated proteins altered with Aquamin.**

| Pathways | Entities pValue | Mapped entities |
|---|---|---|
| Formation of the cornified envelope | $2.62\times10^{-8}$ | DSG2;KRT16;KRT9;KRT2;KRT77;KRT80;KRT5;KRT10;KRT1;KRT14 |
| Keratinization | $2.91\times10^{-6}$ | DSG2;KRT16;KRT9;KRT2;KRT77;KRT80;KRT5;KRT10;KRT1;KRT14 |
| Post-translational modification: synthesis of GPI-anchored proteins | $5.09\times10^{-4}$ | PSCA;MELTF;LYPD8;VNN1;CEACAM5 |
| Release of apoptotic factors from the mitochondria | 0.001 | CYCS;GSDMD |
| Type I hemidesmosome assembly | 0.003 | KRT5;KRT14 |
| Synthesis of bile acids and bile salts via 24-hydroxycholesterol | 0.004 | AKR1C2;SLC27A2 |
| Defective SLC26A3 causes congenital secretory chloride diarrhea 1 | 0.007 | SLC26A3 |
| Apoptotic factor-mediated response | 0.009 | CYCS;GSDMD |
| Regulated Necrosis | 0.010 | OGT;CYCS;GSDMD |
| Synthesis of bile acids and bile salts via 7alpha-hydroxycholesterol | 0.012 | AKR1C2;SLC27A2 |
| Cytosolic sulfonation of small molecules | 0.013 | SULT1A4;SULT1B1 |
| Inhibition of PKR | 0.014 | EIF2AK2 |
| Pyroptosis | 0.016 | CYCS;GSDMD |
| Neutrophil degranulation | 0.019 | TMBIM1;OLFM4;FLG2;GSDMD;PTPRJ;KRT1;SLC27A2;VNN1 |
| Developmental Biology | 0.022 | DSG2;KRT16;KRT9;KRT2;ZNF638;FARP2;KRT80;KRT5;KRT10; PCK1;PLXND1;KRT77;KRT1;KRT14 |
| Synthesis of bile acids and bile salts | 0.024 | AKR1C2;SLC27A2 |
| HHAT G278V doesn't palmitoylate Hh-Np | 0.028 | IHH |
| Biosynthesis of D-series resolvins | 0.028 | HPGD |
| Cell junction organization | 0.028 | CDH17;KRT5;KRT14 |
| Drug ADME | 0.033 | SULT1A4;GSTA1;PCK1 |
| Phosphorylation of CD3 and TCR zeta chains | 0.034 | PTPRJ;HLA-DRB5 |
| NR1H2 & NR1H3 regulate gene expression linked to gluconeogenesis | 0.034 | PCK1 |
| Abacavir metabolism | 0.034 | PCK1 |
| Biosynthesis of E-series 18(S)-resolvins | 0.034 | HPGD |
| Respiratory electron transport | 0.036 | COX1;CYCS;NUBPL |
| Bile acid and bile salt metabolism | 0.040 | AKR1C2;SLC27A2 |
| Activation of caspases through apoptosome-mediated cleavage | 0.041 | CYCS |
| Alpha-oxidation of phytanate | 0.041 | SLC27A2 |
| RUNX2 regulates chondrocyte maturation | 0.041 | IHH |
| Synthesis of Lipoxins (LX) | 0.041 | HPGD |
| Biosynthesis of EPA-derived SPMs | 0.041 | HPGD |
| Phase II - Conjugation of compounds | 0.043 | SULT1A4;GSTA1;SULT1B1 |
| SMAC(DIABLO)-mediated dissociation of IAP:caspase complexes | 0.048 | CYCS |
| SMAC (DIABLO) binds to IAPs | 0.048 | CYCS |

The pathway analysis report was created by employing Reactome pathway database (v82) for species “Homo sapiens.” The listed pathways were curated by submitting up-regulated proteins altered with Aquamin treatment and presented in Table 4.
