## Supplement Table 8 for "Red Algae-Derived Mineral Intervention to Counter Pro-inflammatory Activity in Human Colon Organoids"

**Supplement Table 8. Proteins of Interest (Inflammation-related)**

| Proteins | Genes | Treatment Groups | | |
|---|---|---|---|---|
| | | **Aquamin** | **LPS-Cytokines +Aquamin** | LPS-Cytokines |
| Fibrinogen beta chain | FGB | **0.43±0.28*** | **0.53±0.27*** | 1.12±0.70 |
| Fibrinogen gamma chain | FGG | **0.46±0.16*#** | **0.59±0.24*** | 0.89±0.20 |
| SPARC | SPARC | **0.49±0.06*#** | **0.54±0.13*** | 0.87±0.22 |
| Phospholipase A2, membrane associated | PLA2G2A | **0.72±0.25#** | **1.74±1.81#** | 5.60±1.11* |
| Phosphoglycerate kinase 1 | PGK1 | **1.31±0.40** | **1.33±0.54** | 0.99±0.02 |
| Protein kinase C alpha type | PRKCA | **1.31±0.42** | **1.37±0.39** | 0.94±0.11 |
| Glutamine synthetase | GLUL | **1.40±0.15*** | **1.66±0.67** | 1.05±0.34 |
| Homeobox protein CDX-2 | CDX2 | **1.41±0.21*#** | **1.35±0.52** | 0.97±0.11 |
| Fructose-1,6-bisphosphatase isozyme 2 | FBP2 | **1.47±0.74** | **1.46±0.85** | 0.97±0.09 |
| Natural resistance-associated macrophage protein 2 | SLC11A2 | **1.50±0.64** | **2.09±1.21** | 0.93±0.10 |
| NAD-dependent protein deacetylase sirtuin-3, mitochondrial | SIRT3 | **1.64±1.23** | **2.36±2.36** | 0.98±0.24 |
| Sialidase-1 | NEU1 | **1.66±0.91** | **1.71±0.93** | 1.00±0.26 |
| Protein S100-A7 | S100A7 | **1.68±0.23*#** | **2.26±1.87** | 0.95±0.47 |
| Phenazine biosynthesis-like domain-containing protein | PBLD | **1.95±0.62*#** | **1.63±0.29*#** | 0.87±0.08* |
| 15-hydroxyprostaglandin dehydrogenase [NAD(+)] | HPGD | **2.05±0.69*#** | **1.50±0.61** | 1.00±0.16 |
| ETS homologous factor | EHF | **2.12±2.18** | **1.44±0.66** | 1.17±0.28 |
| Glutathione S-transferase A1 | GSTA1 | **2.45±1.87** | **2.42±3.12** | 0.56±0.24* |
| Dermcidin | DCD | **3.27±1.41*#** | **3.05±3.12** | 1.00±0.54 |

Values represent average abundance ratio from organoids (n=3 subjects) as compared to the control ± standard deviation. Aquamin and Aquamin plus LPS-cytokine treatments (**bold**): These proteins were up- or down-regulated regardless of the fold change (<1% FDR) in response to both Aquamin treatments. Corresponding average abundance ratios are provided from the LPS-Cytokines treatment group for comparison. *Represents significance as compared to the control and #represents significance as compared to LPS-Cytokines (at $p<0.05$).
