## Supplement Table 9 for "Red Algae-Derived Mineral Intervention to Counter Pro-inflammatory Activity in Human Colon Organoids"

**Supplement Table 9. Pathways associated with proteins of interest (presented in Supplement Table 8)**

| Pathways | Entities pValue | Mapped entities |
|---|---|---|
| Response to elevated platelet cytosolic Ca2+ | $4.61 \times 10^{-05}$ | PRKCA;FGG;SPARC;FGB |
| p130Cas linkage to MAPK signaling for integrins | $2.51 \times 10^{-04}$ | FGG;FGB |
| GRB2:SOS provides linkage to MAPK signaling for Integrins | $2.51 \times 10^{-04}$ | FGG;FGB |
| MyD88 deficiency (TLR2/4) | $4.02 \times 10^{-04}$ | FGG;FGB |
| Antimicrobial peptides | $4.06 \times 10^{-04}$ | S100A7;PLA2G2A;DCD |
| IRAK4 deficiency (TLR2/4) | $4.45 \times 10^{-04}$ | FGG;FGB |
| Regulation of TLR by endogenous ligand | $4.90 \times 10^{-04}$ | FGG;FGB |
| Common Pathway of Fibrin Clot Formation | $5.37 \times 10^{-04}$ | FGG;FGB |
| Platelet activation, signaling and aggregation | $6.39 \times 10^{-04}$ | PRKCA;FGG;SPARC;FGB |
| Integrin signaling | $8.65 \times 10^{-04}$ | FGG;FGB |
| Platelet degranulation | $9.62 \times 10^{-04}$ | FGG;SPARC;FGB |
| Extracellular matrix organization | 0.001 | PRKCA;FGG;SPARC;FGB |
| Diseases associated with the TLR signaling cascade | 0.001 | FGG;FGB |
| Diseases of Immune System | 0.001 | FGG;FGB |
| Gluconeogenesis | 0.001 | FBP2;PGK1 |
| Signaling by high-kinase activity BRAF mutants | 0.001 | FGG;FGB |
| Defective SLC11A2 causes hypochromic microcytic anemia, with iron overload 1 (AHMIO1) | 0.002 | SLC11A2 |
| Formation of Fibrin Clot (Clotting Cascade) | 0.002 | FGG;FGB |
| Platelet Aggregation (Plug Formation) | 0.002 | FGG;FGB |
| MAP2K and MAPK activation | 0.002 | FGG;FGB |
| Signaling by RAF1 mutants | 0.002 | FGG;FGB |
| Signaling downstream of RAS mutants | 0.002 | FGG;FGB |
| Paradoxical activation of RAF signaling by kinase inactive BRAF | 0.002 | FGG;FGB |
| Signaling by moderate kinase activity BRAF mutants | 0.002 | FGG;FGB |
| Signaling by RAS mutants | 0.002 | FGG;FGB |
| Defective NEU1 causes sialidosis | 0.005 | NEU1 |
| Manipulation of host energy metabolism | 0.005 | PGK1 |
| Signaling by BRAF and RAF1 fusions | 0.005 | FGG;FGB |
| Scavenging by Class H Receptors | 0.006 | SPARC |
| Neurotransmitter uptake and metabolism in glial cells | 0.006 | GLUL |
| Astrocytic Glutamate-Glutamine Uptake and Metabolism | 0.006 | GLUL |
| Biosynthesis of D-series resolvins | 0.006 | HPGD |
| Oncogenic MAPK signaling | 0.007 | FGG;FGB |
| Innate Immune System | 0.008 | S100A7;FGG;NEU1;PLA2G2A;DCD;FGB |
| Integrin cell surface interactions | 0.008 | FGG;FGB |
| Disinhibition of SNARE formation | 0.008 | PRKCA |
| Biosynthesis of E-series 18(S)-resolvins | 0.008 | HPGD |
| Metal sequestration by antimicrobial proteins | 0.009 | S100A7 |
| Synthesis of Lipoxins (LX) | 0.009 | HPGD |
| Biosynthesis of EPA-derived SPMs | 0.009 | HPGD |
| Glucose metabolism | 0.010 | FBP2;PGK1 |
| Metabolism | 0.010 | FBP2;GSTA1;PRKCA;PGK1;HPGD;NEU1;PLA2G2A;GLUL |
| HuR (ELAVL1) binds and stabilizes mRNA | 0.012 | PRKCA |

| | | |
|---|---|---|
| MyD88:MAL(TIRAP) cascade initiated on plasma membrane | 0.013 | FGG;FGB |
| Toll Like Receptor TLR6:TLR2 Cascade | 0.013 | FGG;FGB |
| ROBO receptors bind AKAP5 | 0.014 | PRKCA |
| Toll Like Receptor TLR1:TLR2 Cascade | 0.014 | FGG;FGB |
| Toll Like Receptor 2 (TLR2) Cascade | 0.014 | FGG;FGB |
| EGFR Transactivation by Gastrin | 0.015 | PRKCA |
| Acetylcholine regulates insulin secretion | 0.015 | PRKCA |
| Regulation of FOXO transcriptional activity by acetylation | 0.015 | SIRT3 |
| Toll Like Receptor 4 (TLR4) Cascade | 0.021 | FGG;FGB |
| Hemostasis | 0.023 | PRKCA;FGG;SPARC;FGB |
| WNT5A-dependent internalization of FZD4 | 0.023 | PRKCA |
| Glutamate and glutamine metabolism | 0.023 | GLUL |
| Synthesis of Prostaglandins (PG) and Thromboxanes (TX) | 0.023 | HPGD |
| Depolymerisation of the Nuclear Lamina | 0.024 | PRKCA |
| Regulation of KIT signaling | 0.024 | PRKCA |
| ER-Phagosome pathway | 0.025 | FGG;FGB |
| Trafficking of GluR2-containing AMPA receptors | 0.026 | PRKCA |
| Acyl chain remodelling of PI | 0.026 | PLA2G2A |
| Biosynthesis of DHA-derived SPMs | 0.026 | HPGD |
| Acyl chain remodelling of PG | 0.028 | PLA2G2A |
| Toll-like Receptor Cascades | 0.028 | FGG;FGB |
| Gastrin-CREB signalling pathway via PKC and MAPK | 0.029 | PRKCA |
| Biosynthesis of specialized proresolving mediators (SPMs) | 0.029 | HPGD |
| Antigen processing-Cross presentation | 0.030 | FGG;FGB |
| VEGFR2 mediated cell proliferation | 0.031 | PRKCA |
| Diseases associated with N-glycosylation of proteins | 0.031 | NEU1 |
| Synthesis, secretion, and inactivation of Glucagon-like Peptide-1 (GLP-1) | 0.032 | CDX2 |
| Acyl chain remodelling of PS | 0.034 | PLA2G2A |
| Azathioprine ADME | 0.035 | GSTA1 |
| RHO GTPases Activate NADPH Oxidases | 0.037 | PRKCA |
| Incretin synthesis, secretion, and inactivation | 0.037 | CDX2 |
| Metal ion SLC transporters | 0.040 | SLC11A2 |
| Acyl chain remodelling of PC | 0.041 | PLA2G2A |
| Syndecan interactions | 0.041 | PRKCA |
| SHC1 events in ERBB2 signaling | 0.042 | PRKCA |
| Acyl chain remodelling of PE | 0.044 | PLA2G2A |
| Metabolism of porphyrins | 0.044 | GSTA1 |
| FOXO-mediated transcription of oxidative stress, metabolic and neuronal genes | 0.045 | SIRT3 |
| Trafficking of AMPA receptors | 0.047 | PRKCA |
| Glutamate binding, activation of AMPA receptors and synaptic plasticity | 0.047 | PRKCA |
| Sialic acid metabolism | 0.050 | NEU1 |
| Inactivation, recovery and regulation of the phototransduction cascade | 0.050 | PRKCA |
| Nuclear signaling by ERBB4 | 0.051 | SPARC |
| Calmodulin induced events | 0.053 | PRKCA |
| Glutathione conjugation | 0.056 | GSTA1 |

The pathway analysis report was created by employing Reactome pathway database (v82) for species “Homo sapiens.” The listed pathways were curated by submitting altered proteins compiled after a targeted search and are presented in Supplement Table 8.
