## Supplemental File 1 for "Red Algae-Derived Mineral Intervention to Counter Pro-inflammatory Activity in Human Colon Organoids"

**Supplemental File 1. Additional References (Relevant to the moieties presented in the Supplement Table 8).**

**Fibrinogen beta chain (FGB) / Fibrinogen gamma chain (FGG):**


- Zhang C, Chen H, He Q, Luo Y, He A, Tao A, Yan J. Fibrinogen/AKT/Microfilament Axis Promotes Colitis by Enhancing Vascular Permeability. Cell Mol Gastroenterol Hepatol. 2021;11(3):683-696. doi: 10.1016/j.jcmgh.2020.10.007. Epub 2020 Oct 17. PMID: 33075564; PMCID: PMC7843406.

- Luyendyk JP, Schoenecker JG, Flick MJ. The multifaceted role of fibrinogen in tissue injury and inflammation. Blood. 2019 Feb 7;133(6):511-520. doi: 10.1182/blood-2018-07-818211. Epub 2018 Dec 6. PMID: 30523120; PMCID: PMC6367649.


**SPARC:**


- Fonseca-Camarillo G, Furuzawa-Carballeda J, Razo-López N, Barreto-Zúñiga R, Martínez-Benítez B, Yamamoto-Furusho JK. Intestinal production of secreted protein acidic and rich in cysteine (SPARC) in patients with ulcerative colitis. Immunobiology. 2021 May;226(3):152095. doi: 10.1016/j.imbio.2021.152095. Epub 2021 May 8. PMID: 34000572.
- Ng Y-Lm Klopcic B, Lloyd F, Forrst C, Greene W, Lawrance IC. Secreted protein acidic and rich in cysteine (SPARC) exacerbates colnic inflammatory symptoms in dextran sodium sulphate-induced murine colitis. PLoS One 8 (10) e77575 doi:10:1371/journal.pone.0077575.


**Phospholipase A2, membrane associated (PLA2G2A):**


- Escudero-Hernández C, van Beelen Granlund A, Bruland T, Sandvik AK, Koch S, Østvik AE, Münch A. Transcriptomic Profiling of Collagenous Colitis Identifies Hallmarks of Nondestructive Inflammatory Bowel Disease. Cell Mol Gastroenterol Hepatol. 2021;12(2):665-687. doi: 10.1016/j.jcmgh.2021.04.011. Epub 2021 Apr 27. PMID: 33930606; PMCID: PMC8267496.
- Doré E, Joly-Beauparlant C, Morozumi S, Mathieu A, Lévesque T, Allaeys I, Duchez AC, Cloutier N, Leclercq M, Bodein A, Payré C, Martin C, Petit-Paitel A, Gelb MH, Rangachari M, Murakami M, Davidovic L, Flamand N, Arita M, Lambeau G, Droit A, Boilard E. The interaction of secreted phospholipase A2-IIA with the microbiota alters its lipidome and promotes inflammation. JCI Insight. 2022 Jan 25;7(2):e152638. doi: 10.1172/jci.insight.152638. PMID: 35076027; PMCID: PMC8855825.


**Phosphoglycerate kinase 1 (PGK1):**


- Liu J, Zhao W, Li C, Wu T, Han L, Hu Z, Li X, Zhou J, Chen X. Terazosin Stimulates Pgk1 to Remedy Gastrointestinal Disorders. Int J Mol Sci. 2021 Dec 30;23(1):416. doi: 10.3390/ijms23010416. PMID: 35008842; PMCID: PMC8745693.


**Protein kinase C alpha type (PRKCA):**


- Wang M, Zhong H, Zhang X, Huang X, Wang J, Li Z, Chen M, Xiao Z. EGCG promotes PRKCA expression to alleviate LPS-induced acute lung injury and inflammatory response. Sci Rep. 2021 May 26;11(1):11014. doi: 10.1038/s41598-021-90398-x. PMID: 34040072; PMCID: PMC8154949.


**Glutamine synthetase (GLUL):**


- Zhou Q, Souba WW, Croce CM, Verne GN. MicroRNA-29a regulates intestinal membrane permeability in patients with irritable bowel syndrome. Gut. 2010 Jun;59(6):775-84. doi: 10.1136/gut.2009.181834. Epub 2009 Dec 1. PMID: 19951903; PMCID: PMC2891786.

- Yu J, Zhang J, Shi M, Ding H, Ma L, Zhang H, Liu J. Maintenance of glutamine synthetase expression alleviates endotoxin-induced sepsis via alpha-ketoglutarate-mediated demethylation. FASEB J. 2022 May;36(5):e22281. doi: 10.1096/fj.202200059R. PMID: 35344214.


**Homeobox protein CDX-2 (CDX2):**


- Stolfi C, Maresca C, Monteleone G, Laudisi F. Implication of Intestinal Barrier Dysfunction in Gut Dysbiosis and Diseases. Biomedicines. 2022 Jan 27;10(2):289. doi: 10.3390/biomedicines10020289. PMID: 35203499; PMCID: PMC8869546.
- Coskun M, Olsen AK, Holm TL, Kvist PH, Nielsen OH, Riis LB, Olsen J, Troelsen JT. TNF-α-induced down-regulation of CDX2 suppresses MEP1A expression in colitis. Biochim Biophys Acta. 2012 Jun;1822(6):843-51. doi: 10.1016/j.bbadis.2012.01.012. Epub 2012 Feb 3. PMID: 22326557.


**Fructose-1,6-bisphosphatase isozyme 2 (FBP2):**


- Li H, Wang J, Xu H, Xing R, Pan Y, Li W, Cui J, Zhang H, Lu Y. Decreased fructose-1,6-bisphosphatase-2 expression promotes glycolysis and growth in gastric cancer cells. Mol Cancer. 2013 Sep 25;12(1):110. doi: 10.1186/1476-4598-12-110. PMID: 24063558; PMCID: PMC3849177.
- Danckwardt S, Gantzert AS, Macher-Goeppinger S, Probst HC, Gentzel M, Wilm M, Gröne HJ, Schirmacher P, Hentze MW, Kulozik AE. p38 MAPK controls prothrombin expression by regulated RNA 3' end processing. Mol Cell. 2011 Feb 4;41(3):298-310. doi: 10.1016/j.molcel.2010.12.032. PMID: 21292162.


**Natural resistance-associated macrophage protein 2 (SLC11A2 or NRAMP2):**


- Forbes JR, Gros P. Divalent-metal transport by NRAMP proteins at the interface of host-pathogen interactions. Trends Microbiol. 2001 Aug;9(8):397-403. doi: 10.1016/s0966-842x(01)02098-4. PMID: 11514223.
- Wardrop SL, Wells C, Ravasi T, Hume DA, Richardson DR. Induction of Nramp2 in activated mouse macrophages is dissociated from regulation of the Nramp1, classical inflammatory genes, and genes involved in iron metabolism. J Leukoc Biol. 2002 Jan;71(1):99-106. PMID: 11781385.


**NAD-dependent protein deacetylase sirtuin-3, mitochondrial (SIRT3):**


- Zhang, Y., Wang, X. L., Zhou, M., Kang, C., Lang, H. D., Chen, M. T., Hui, S. C., Wang, B., & Mi, M. T. (2018). Crosstalk between gut microbiota and Sirtuin-3 in colonic inflammation and tumorigenesis. Experimental & molecular medicine, 50(4), 1–11. https://doi.org/10.1038/s12276-017-0002-0.
- Heinonen, T., Ciarlo, E., Le Roy, D., & Roger, T. (2019). Impact of the Dual Deletion of the Mitochondrial Sirtuins SIRT3 and SIRT5 on Anti-microbial Host Defenses. Frontiers in immunology, 10, 2341. https://doi.org/10.3389/fimmu.2019.02341


**Sialidase-1 (NEU1):**


- Lillehoj EP, Luzina IG, Atamas SP. Mammalian Neuraminidases in Immune-Mediated Diseases: Mucins and Beyond. Front Immunol. 2022 Apr 11;13:883079. doi: 10.3389/fimmu.2022.883079. PMID: 35479093; PMCID: PMC9035539.
- Bonten E, van der Spoel A, Fornerod M, Grosveld G, d'Azzo A. Characterization of human lysosomal neuraminidase defines the molecular basis of the metabolic storage disorder sialidosis. Genes Dev. 1996 Dec 15;10(24):3156-69. doi: 10.1101/gad.10.24.3156. PMID: 8985184.


**Protein S100-A7 (S100A7):**

- Michalek, M., Gelhaus, C., Hecht, O., Podschun, R., Schröder, J. M., Leippe, M., & Grötzinger, J. (2009). The human antimicrobial protein psoriasin acts by permeabilization of bacterial membranes. Developmental and comparative immunology, 33(6), 740–746. https://doi.org/10.1016/j.dci.2008.12.005
- Gläser, R., Harder, J., Lange, H., Bartels, J., Christophers, E., & Schröder, J. M. (2005). Antimicrobial psoriasin (S100A7) protects human skin from Escherichia coli infection. Nature immunology, 6(1), 57–64. https://doi.org/10.1038/ni1142


**Phenazine biosynthesis-like domain-containing protein (PBLD):**


- Chen, S., Liu, H., Li, Z., Tang, J., Huang, B., Zhi, F., & Zhao, X. (2021). Epithelial PBLD attenuates intestinal inflammatory response and improves intestinal barrier function by inhibiting NF-κB signaling. Cell death & disease, 12(6), 563. https://doi.org/10.1038/s41419-021-03843-0
- Zhao, X., Kang, B., Lu, C., Liu, S., Wang, H., Yang, X., Chen, Y., Jiang, B., Zhang, J., Lu, Y., & Zhi, F. (2011). Evaluation of p38 MAPK pathway as a molecular signature in ulcerative colitis. Journal of proteome research, 10(5), 2216–2225. https://doi.org/10.1021/pr100969w


**15-hydroxyprostaglandin dehydrogenase [NAD(+)] (HPGD):**


- Gibbs DC, Fedirko V, Baron JA, Barry EL, Flanders WD, McCullough ML, Yacoub R, Raavi T, Rutherford RE, Seabrook ME, Bostick RM. Inflammation Modulation by Vitamin D and Calcium in the Morphologically Normal Colorectal Mucosa of Patients with Colorectal Adenoma in a Clinical Trial. Cancer Prev Res (Phila). 2021 Jan;14(1):65-76. doi: 10.1158/1940-6207.CAPR-20-0140. Epub 2020 Sep 11. PMID: 32917645; PMCID: PMC7947029.
- Yan, M., Rerko, R. M., Platzer, P., Dawson, D., Willis, J., Tong, M., Lawrence, E., Lutterbaugh, J., Lu, S., Willson, J. K., Luo, G., Hensold, J., Tai, H. H., Wilson, K., & Markowitz, S. D. (2004). 15-Hydroxyprostaglandin dehydrogenase, a COX-2 oncogene antagonist, is a TGF-beta-induced suppressor of human gastrointestinal cancers. Proceedings of the National Academy of Sciences of the United States of America, 101(50), 17468–17473. https://doi.org/10.1073/pnas.0406142101


**ETS homologous factor (EHF):**


- Reehorst CM, Nightingale R, Luk IY, Jenkins L, Koentgen F, Williams DS, Darido C, Tan F, Anderton H, Chopin M, Schoffer K, Eissmann MF, Buchert M, Mouradov D, Sieber OM, Ernst M, Dhillon AS, Mariadason JM. EHF is essential for epidermal and colonic epithelial homeostasis, and suppresses Apc-initiated colonic tumorigenesis. Development. 2021 Jun 15;148(12):dev199542. doi: 10.1242/dev.199542. Epub 2021 Jun 28. PMID: 34180969.


**Glutathione S-transferase A1 (GSTA1):**


- Clapper ML, Adrian RH, Pfeiffer GR, Kido K, Everley L, Cooper HS, Murthy S. Depletion of colonic detoxication enzyme activity in mice with dextran sulphate sodium-induced colitis. Aliment Pharmacol Ther. 1999 Mar;13(3):389-96. doi: 10.1046/j.1365-2036.1999.00475.x. PMID: 10102973.
- Langmann T, Moehle C, Mauerer R, Scharl M, Liebisch G, Zahn A, Stremmel W, Schmitz G. Loss of detoxification in inflammatory bowel disease: dysregulation of pregnane X receptor target genes. Gastroenterology. 2004 Jul;127(1):26-40. doi: 10.1053/j.gastro.2004.04.019. PMID: 15236169.


**Dermcidin (DCD):**


- Schittek B, Hipfel R, Sauer B, Bauer J, Kalbacher H, Stevanovic S, Schirle M, Schroeder K, Blin N, Meier F, Rassner G, Garbe C. Dermcidin: a novel human antibiotic peptide secreted by sweat glands. Nat Immunol. 2001 Dec;2(12):1133-7. doi: 10.1038/ni732. PMID: 11694882.

- Song C, Weichbrodt C, Salnikov ES, Dynowski M, Forsberg BO, Bechinger B, Steinem C, de Groot BL, Zachariae U, Zeth K. Crystal structure and functional mechanism of a human antimicrobial membrane channel. Proc Natl Acad Sci U S A. 2013 Mar 19;110(12):4586-91. doi: 10.1073/pnas.1214739110. Epub 2013 Feb 20. PMID: 23426625; PMCID: PMC3607029.
